# Nerve-associated macrophages control adipose homeostasis across lifespan and restrain age-related inflammation

**DOI:** 10.1101/2024.10.12.618004

**Authors:** Elsie Gonzalez-Hurtado, Claire Leveau, Keyi Li, Rihao Qu, Manish Mishra, Emily L. Goldberg, Sviatoslav Sidorov, Stephen T. Yeung, Camille Khairallah, David Gonzalez, Taverlyn M. Shepard, Christina Camell, Maxim N. Artyomov, Yuval Kluger, Kamal M. Khanna, Vishwa Deep Dixit

**Author notes:** Authors with equal contribution (Elsie Gonzalez-Hurtado, Claire Leveau).

## Abstract

Age-related inflammation or ‘inflammaging’ is a key mechanism that increases disease burden and may control lifespan. How adipose tissue macrophages (ATMs) control inflammaging is not well understood in part because the molecular identities of niche-specific ATMs are incompletely known. Using intravascular labeling to exclude circulating myeloid cells and subsequent single-cell sequencing with orthogonal validation, we define the diversity and alterations in niche resident ATMs through lifespan. Aging led to depletion of vessel-associated macrophages (VAMs), expansion of lipid-associated macrophages (LAMs), and emergence of a unique subset of CD38^+^ age-associated macrophages (AAMs) in visceral white adipose tissue (VAT). Interestingly, CD169^+^CD11c^−^ ATMs are enriched in a subpopulation of nerve-associated macrophages (NAMs) that declines with age. Depletion of CD169^+^ NAMs in aged mice increases inflammaging and impairs lipolysis suggesting that they are necessary for preventing catecholamine resistance in VAT. These findings reveal specialized ATMs control adipose homeostasis and link inflammation to tissue dysfunction during aging.

## Introduction

The sympathetic nervous system (SNS) along with innate immune system are the first lines of defense in sensing danger and protection of tissues from internal and external threats.The emergence of chronic inflammation is considered a hallmark feature of the aging process that is directly linked to a decline in tissue function ^1,2^. Adipose tissue macrophages (ATMs) are predominant immune cells in the adipose tissue. ATMs do not display binary M1 or M2-like phenotype and are known to decline with age ^3–5^. Notably, ATMs reside in specialized niches such as fat-associated lymphoid clusters, crown-like structures that contain necrotic adipocytes, and near vasculature and/or sympathetic nerves. Recent studies of ATMs have revealed the presence of diverse macrophage states such as lipid-associated macrophages (LAMs) and vascular-associated macrophages (VAMs) ^6,7^. However, the changes that may occur amongst distinct ATM subsets in aging, and the functional role these cells play through lifespan remain largely unknown.

A population of tissue resident macrophages referred to here as nerve-associated macrophages (NAMs) have also been described in several tissues including: the gut, lung, skin, and adipose tissue ^3,8–13^. NAMs have been linked to a wide array of functions including the regulation of peristalsis (gut), the immunoregulatory response to influenza (lung), nerve surveillance and repair (skin and sciatic nerve), and catecholamine degradation (adipose) ^3,8–14^. In the adipose tissue, NAMs appear to regulate obesity and may play a role in fasting-induced lipolysis during aging ^3,12^. However, given the lack of specific markers or Cre drivers that target NAMs, or distinguish them from other ATMs, the precise functional role of NAMs in adipose tissue homeostasis and aging remains unclear. Moreover, despite several subsequent comprehensive single-cell transcriptomic analyses, the exact transcriptional identity of adipose tissue NAMs continues to remain elusive, as so far, no study has yet identified a NAM-specific sub-cluster or transcriptomic signature using an unbiased scRNAseq approach.

Here, we implemented an unbiased approach to meticulously dissect the resident F4/80^+^CD11b^+^ population in young and aged visceral white adipose tissue (VAT). Using intra-vascular (iv) labeling in combination with bulk RNA-sequencing we show that resident F4/80^+^CD11b^+^ cells are transcriptionally distinct from circulating F4/80^+^CD11b^+^ cells. Furthermore, using single cell RNA-sequencing (scRNAseq) we find that the resident F4/80^+^CD11b^+^ population can be subdivided into 13 subsets that include vascular-associated macrophages (VAMs), lipid-associated macrophages (LAMs), interferon-associated macrophages (IAMs), CD169^+^ nerve-associated macrophages (NAMs), and CD38^+^ age-associated macrophages (AAMs) that emerge only in aging. Using CD169 diphtheria toxin receptor (CD169-DTR) mice, we demonstrate that CD169^+^ NAMs are pivotal in regulating lipolysis and are essential for preventing catecholamine resistance and adipose tissue dysfunction. We further reveal that CD169^+^ NAMs diminish in adipose tissue with age and are crucial for mitigating age-induced inflammation.

## Results

### Singe-cell characterization of resident adipose tissue macrophages from the visceral white adipose tissue (VAT)

The VAT expands with aging, is highly vascularized and contains circulating myeloid cells that are likely transcriptionally distinct from tissue-resident macrophages or monocytes that have recently undergone extravasation. To investigate this, we performed intravascular (iv) labeling with anti-CD45.2 antibody to label circulating (CD45^+^CD45.2iv^+^) and resident (CD45^+^CD45.2iv^−^) cells ^15,16^. Resident and circulating F4/80^+^CD11b^+^ cells were sorted from the VAT of young (2-month) and aged (22-month) male and female mice (**Fig. 1A**). Flow cytometry analysis showed that in the VAT, approximately 20-30% of F4/80^+^CD11b^+^ cells were in the vasculature, and 70-80% of F4/80^+^CD11b^+^ cells resided in the tissue (**Fig. 1B**). Subsequent bulk RNA-sequencing and principal component analysis (PCA) confirmed that the resident F4/80^+^CD11b^+^ population exhibits a distinct gene expression profile compared to the circulating F4/80^+^CD11b^+^ population (**Fig. 1C**). Together, these data highlight that approximately 75% of the F4/80^+^CD11b^+^ cells isolated from the VAT stromal vascular fraction (SVF) are in the tissue parenchyma.

**Figure 1.**
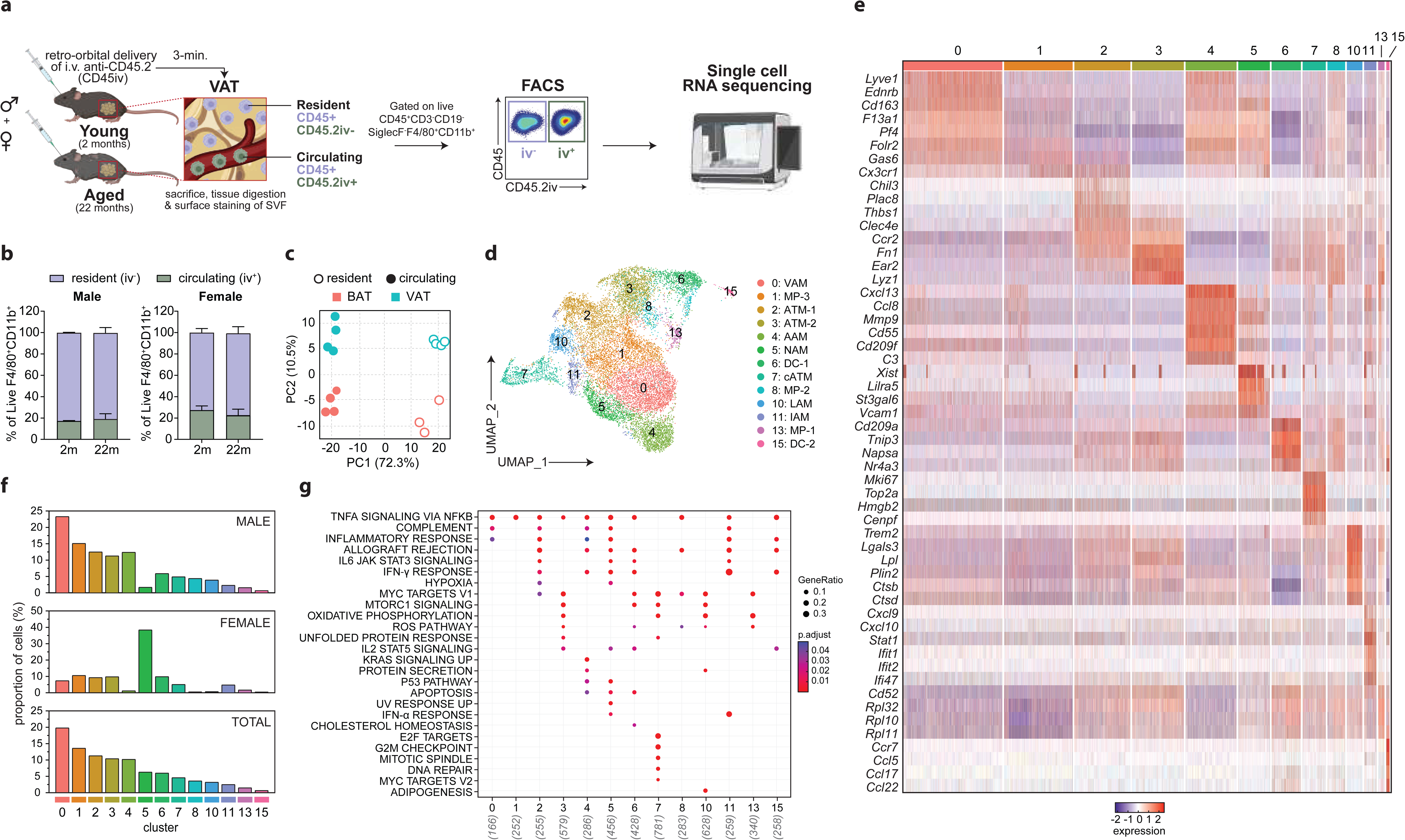
Singe-cell characterization of adipose tissue macrophages from the visceral white adipose tissue (VAT). (A) Schematic of experimental procedure for intra-vascular (iv) labeling of circulating cells with CD45.2 antibody (CD45.2iv) via retro-orbital delivery in 2-month-old and 22-month-old male and female mice. CD45.2 antibody was allowed to circulate for 3-minutes, after which animals were sacrificed. The VAT stromal vascular fraction (SVF) was collected and stained with antibodies against CD45, CD3, CD19, SiglecF, CD11b, and F4/80. Resident and circulating F4/80^+^CD11b^+^ cells (Resident: live CD45^+^CD45.2iv^−^CD3^−^CD19^−^SiglecF^−^, Circulating: live CD45^+^CD45.2iv^+^CD3^−^CD19^−^SiglecF^−^) from the VAT were sorted using fluorescence-activated cell sorting (FACS) and sequenced via bulk- and single cell RNA-sequencing (scRNAseq). (B) Quantification of resident (CD45^+^CD45.2iv^−^CD3^−^CD19^−^SiglecF^−^) and circulating (CD45^+^CD45.2iv^+^CD3^−^CD19^−^SiglecF^−^) F4/80^+^CD11b^+^ cells from VAT of 2-month and 22-month-old male and female mice via flow cytometry. Data is representative of 3 independent experiments with n=3-4 per group. (C) Principle Component Analysis (PCA) plot of resident (CD45^+^CD45.2iv^−^CD3^−^CD19^−^SiglecF^−^) and circulating (CD45^+^CD45.2iv^+^CD3^−^CD19^−^SiglecF^−^) F4/80^+^CD11b^+^ cells from the brown adipose tissue (BAT) and VAT of 2- and 22-month-old males and females. (D) Uniform Manifold Approximation and Projection (UMAP) representation of unsupervised clustering of 14,697 resident F4/80^+^CD11b^+^ (CD45^+^CD45.2iv^−^CD3^−^CD19^−^SiglecF^−^) cells sorted from VAT of young and aged males (12,685 cells) and females (2,012 cells). UMAP excludes clusters that lacked significant expression of *Ptprc* (CD45), *Itgam* (CD11b), and *Adgre1* (F4/80). Cluster abbreviations are as follows: Myeloid Precursors (MP), Adipose tissue macrophages (ATM), Dendritic Cells (DC), Cycling ATMs (cATM), Vessel-Associated Macrophages (VAM), Age-associated Macrophages (VAM), Nerve-associated Macrophages (NAM), Interferon-associated Macrophages (IAM), and Lipid-associated Macrophages (LAM). (E) Heatmap of each cell’s (column) scaled expression of differentially expressed genes (DEGs) (row) per cluster, with exemplar genes labeled. (F) Repartition of cells in each cluster as a fraction of total cells profiled (*top*), fraction of cells from males (*middle*), and fraction of cells from females (*bottom*). (G) ‘Hallmark’ gene enrichment analysis per cluster.

To examine the age-related changes in the tissue resident compartment (CD45^+^CD45.2iv^−^), we conducted single cell RNA-sequencing (scRNAseq) of F4/80^+^CD11b^+^ cells sorted from the VAT of young and aged males and females (CD45^+^CD45.2iv^−^CD3^−^CD19^−^SiglecF^−^) (**Fig. 1A**). Unsupervised clustering of 15,709 quality control (QC)-positive cells revealed 16 clusters as visualized by Uniform Manifold Approximation and Projection (UMAP) (**Fig. S1A-B & Table S1**). Clusters 9, 12, and 14 did not express significant levels of *Ptprc* (CD45), *Itgam* (CD11b), and *Adgre1* (F4/80), suggesting these clusters were likely contaminating cells and were therefore excluded from future analyses (**Fig. 1D-E & S1C**). Approximately 60-70% of cells clustered into 5 main populations (Clusters 0-4) and 30-40% of cells clustered into the remaining 8 populations (Clusters 5-8, 10-13, 15) (**Fig. 1F)**. The proportion of cells in each cluster was similar in males and females except for two clusters: cluster 0 was the dominant cluster in males and cluster 5 in females (**Fig. 1F**).

Clusters 0, 1, 4, and 5 represented approximately 20%, 13%, 10%, and 6% of all cells profiled, respectively, and exhibited high expression levels of *Adgre1* (F4/80) relative to other clusters (**Fig. 1F, S1C**). Only a few genes such as *Cx3cr1* were differentially expressed in Cluster 1 compared to other clusters (**Fig. 1E &Table S2**). RNA velocity inference analysis suggested Cluster 1 represented an early differentiation and/or precursor state that could give rise to macrophages from Cluster 0 (**Fig. S1D**). Visualization of the top 25 differentially expressed genes (DEGs) for each cluster showed that Clusters 0, 4, and 5 were enriched for unique as well as a shared subset of genes that included several complement genes (*C1qa*, *C1qb*, *C1qc*, *C4b*, *C3ar1*, *C5ar1*), and genes associated with tissue resident macrophages such as *Cd163*, *Cd209f*, *Cd209g*, *Csf1r*, *Folr2*, and *Gas6* (**Fig. 1E, S1B, & Table S2**) ^8,17,18^. To delineate between Clusters 0, 4, and 5, we compared the DEGs between Cluster 0 versus Clusters 4 and 5 (**Fig. S1E**). This analysis showed that Cluster 0 was enriched for *Lyve1*, *Cd163*, *Ednrb*, *F13a1*, *Apoe*, *Cd36*, and *Ccl24* expression (**Fig. 1E, S1E, & Table S2**). The gene signature for Cluster 0 corresponded to a population of tissue-resident vessel-associated macrophages (VAMs) previously described in the adipose tissue, lungs, and heart ^6,7,13,17,19^. Cluster 4 was enriched for *Cxcl13*, *C3*, *Cd55*, *Ly6e*, *Ccl8*, *Colq*, *Mmp9* genes, and was unique in that it only emerged in the aged condition; hence we refer to this cluster as age-associated macrophages (AAMs) (**Fig. 1E, S1E, Table S2, & 2A**). Cluster 5 was enriched for expression of genes such as *Lilra5*, *H2-Eb1*, *Cd74*, *Cd86*, *Vcam1*, *Ccl12*, *Csf1r*, *St3gal6*, and *Xist* (**Fig. 1E, S1E, & Table S2**). Notably, Cluster 5 showed enrichment for monoamine oxidase-A (*Maoa*), an enzyme required for the breakdown and inactivation of catecholamines that was previously shown to be expressed by sympathetic nerve-associated macrophages (SAMs, referred to here as NAMs) ^3,12^ (**Fig. S2A**). Moreover, relative to Cluster 0 and 4, Cluster 5 expressed low levels of *Lyve1*, a gene marker that has previously been used to discriminate between macrophages adjacent to vessels (Lyve1^hi^MHCII^hi^) or nerves (Lyve1^lo^MHCII^hi^) ^8^ (**Fig. 1E, S2A**). Thus, to assess if Cluster 5 was a *bona fide* NAM cluster, we calculated a gene set enrichment score (GSES) for each cell using a list of NAM-associated genes from studies characterizing NAMs in other tissues ^3,8–13,20,21^ (**Table S3**). Our analysis showed Cluster 5 had the highest GSES, suggesting Cluster 5 represented NAMs in the adipose tissue (**Fig. S2B**). Interestingly, we noted that Clusters 0, 4, and 5 exhibited low levels of *Ccr2* expression relative to other clusters, with *Ccr2* expression levels being the lowest in Cluster 4 (**Fig. S2A**). Conversely, expression of *Timd4*, *Folr2*, and *Lyve1* was largely restricted to cells in Clusters 0 and 4, and to a lesser extent Cluster 5, suggesting as a previous report did, that these macrophage subsets might be maintained via self-renewal with minimal monocyte input (**Fig. S2C**) ^18^.

Cluster 2 represented approximately 11% of all cells profiled (**Fig. 1F**). Analysis of the DEGs for Cluster 2 showed it was enriched for *Ccr2*, *Chil3*, *Plac8*, *Thbs1*, *Clec4e*, *Mpeg1*, *Il1b* and *Ly6c2* genes, suggesting these adipose tissue macrophages maybe of monocyte origin (ATM-1) (**Fig. 1D-E & Table S2**) ^17,22^. Clusters 3, 7, 8, and 13 represented approximately 10%, 5%, 4%, and 1.5% of all cells profiled, respectively, and exhibited intermediate expression levels of *Adgre1* (F4/80) relative to other clusters (**Fig. 1F, S1C, & Table S2**). Analysis of DEGs for Clusters 3, 7, 8, and 13 showed these clusters were enriched for *Fn1*, *Lyz1*, *Ear2*, and *Cd52* expression, with Clusters 3 and 7 exhibiting the highest and lowest expression levels of these genes, respectively (**Fig. 1E & Table S2**). Cluster 3 was distinguished by high expression of *Retnla*, *Lpl*, *Cd226*, and *Socs6*, a gene signature similar to that of small peritoneal macrophages and to a monocyte-derived macrophage cluster previously described in the adipose and heart (ATM-2) (**Fig. 1D-E & Table S2**) ^6,17,22,23^. Cluster 7 was enriched for cell cycle genes such as *Mki67*, *Top2a*, *Hmgb2*, *Ube2c*, and Cluster 13 for ribosomal genes such as *Rpl32*, *Rpl10*, *Rpl11*, *Rps18* (**Fig. 1D-E & Table S2**). The gene signature for Cluster 7 suggested this cluster represented cycling ATMs (cATM), which have previously been reported in the lung and heart ^13,19,22^ (**Fig. 1E & Table S2**). Cluster 10 and 11 each represented approximately 3% and 2.5% of all cells profiled, respectively (**Fig. 1F**). Cluster 10 was characterized by high expression of *Trem2*, *Spp1*, *Lgals3*, *Lpl*, *Ctsl*, *Cstb*, *Cd9*, and *Fabp5*, a gene signature previously associated with lipid-associated macrophages (LAMs) (**Fig. 1D-E & Table S2**) ^6,17^. Cluster 11 expressed high levels of *Cxcl9*, *Cxcl10*, *Rsad2*, *Stat1* and interferon-stimulated genes (ISGs) such as *Ifit1*, *Ifit2*, *Irf7*, *Ifi47*, *Isg15* (**Fig. 1D-E & Table S2**). The gene signature for Cluster 11 has been observed in a subset of macrophages from the heart, hence we refer to them here as interferon-associated macrophages (IAMs) ^22^. On the other hand, Clusters 6 and 15 represented approximately 6% and 0.7% of all cells profiled and expressed the lowest levels of *Adgre1* (F4/80) relative to all clusters (**Fig. 1E, S1B**). While only Cluster 6 expressed statistically significant levels of *Cd209a* and *Itgam* (CD11b), both Clusters 6 and 15 were enriched for several genes associated with dendritic cells including *Itgax*, *Flt3*, *Napsa*, *Nr4a3*, *Zbtb46*, *Ccl17, Ccl22,* and *Dpp4* suggesting these clusters represented CD11b^+^ DCs (DC-1 and DC-2) (**Fig. 1D-E & Table S2**) ^17^. Together, the observed gene signatures along with RNA velocity inference analysis suggest Clusters 13, 8, and 1 represent early myeloid precursor populations (MP-1, MP-2, and MP3) that may give rise to more terminally differentiated tissue resident macrophages (Clusters 0, 4, and 5) or monocyte-derived-macrophages (Cluster 2, 3, 7, 10, 11) and monocyte-derived dendritic cells (Clusters 6 and 15) (**Fig. 1D, S1D & Table S2**).

Next, we calculated a gene set enrichment score (GSES) for each cell using the Hallmark gene collection from the Molecular Signatures Database (MSigDB) ^24–26^. Our analysis confirmed upregulation of complement genes in Clusters 0, 4, and 5, cell cycle genes in Cluster 7, oxidative phosphorylation and adipogenesis genes in Cluster 10, and interferon response genes in Cluster 11 (**Fig. 1G**). Furthermore, Clusters 3, 6, 7, and 10 all showed enrichment for MYC target and MTORC1 signaling genes, while Clusters 4 and 5 were both enriched for genes involved in the p53 pathway and apoptosis (**Fig. 1G**). In addition, Cluster 4 was enriched for genes involved in KRAS signaling and protein secretion, while Cluster 5 showed enrichment for genes involved in Hypoxia and UV response (**Fig. 1G**).

Altogether, our analysis unveils unique ATM states with distinct transcriptional signatures such as VAMs (Cluster 0) and LAMs (Cluster 10), and two tissue resident macrophage clusters not previously identified in the adipose using unbiased scRNAseq profiling of F4/80^+^CD11b^+^ cells: Cluster 5 (NAMs) and Cluster 4 (AAMs).

### Macrophage subsets in the VAT undergo remodeling during aging

The impact of aging on different resident ATM populations and their transcriptome remains to be established. Our unbiased scRNAseq profiling of total resident (non-circulating) F4/80^+^CD11b^+^ cells showed that aging induced perturbations in greater than half of the clusters identified (**Fig. 2A-B, S3A**). The proportion of cells in clusters likely representing precursor or proliferating populations, namely MP-1 and cATM, remained largely unaffected by age (**Fig. 2A-B**). The exceptions were MP-2 and MP-3, which showed a 0.5-fold reduction with age (**Fig. 2A-B**). The proportion of cells in ATM-1 and ATM-2 remained largely constant between conditions, suggesting these clusters might be continually seeded from infiltrating monocytes (**Fig. 2A-B**). VAMs exhibited a 0.3-fold decrease with age, while AAMs, representing 1% of all cells in the young condition represented almost 20% of cells in the aged condition (**Fig. 2A-B**). NAMs represented approximately 5% and 8% of all cells in the young and aged conditions, while the proportion of cells in DC-1 decreased by approximately 0.5-fold in the aged condition (**Fig. 2A-B**). Both LAMs and IAMs exhibited increases with age (**Fig. 2A-B**).

**Figure 2.**
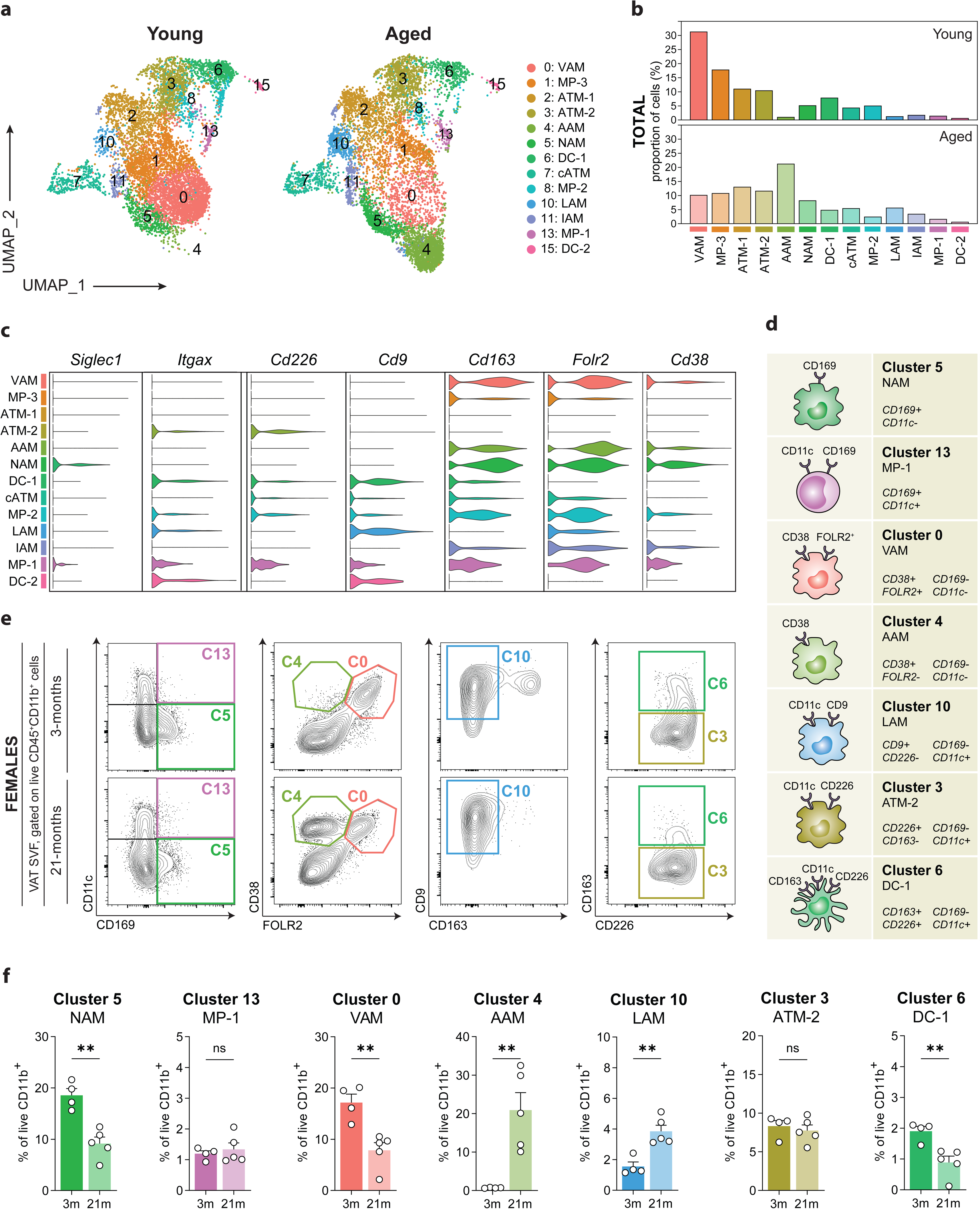
Macrophage subsets in the visceral white adipose tissue (VAT) undergo remodeling during aging. (A) Uniform Manifold Approximation and Projection (UMAP) representation of unsupervised clustering of resident F4/80^+^CD11b^+^ (CD45^+^CD45.2iv^−^CD3^−^CD19^−^SiglecF^−^) cells sorted from the VAT of young (7,558 cells) and aged (7,139 cells) mice from males and females. (B) Repartition of cells in each cluster as a fraction of total cells (males and females) from young and aged VAT. (C) Violin plots depicting the expression of *Siglec1*, *Itgax*, *Cd226*, *Cd9*, *Cd163*, *Folr2*, and *Cd38* genes by cluster. (D) Schematic summarizing the cell surface markers used to define seven clusters identified via single-cell RNA-sequencing of resident F4/80^+^CD11b^+^ cells (CD45^+^CD45.2iv^−^CD3^−^ CD19^−^SiglecF^−^) from the VAT. Abbreviations for each subset and defining cell surface markers are as follows: Nerve-associated-macrophages (NAM, C5): CD169^+^CD11c^−^, Myeloid Precursors-1 (MP-1, C13): CD169^+^CD11c^+^, Vessel-associated Macrophages (VAM, C0): CD38^+^FOLR2^+^CD11c^−^CD38^+^, Age-associated Macrophages (AAM, C4): CD38^+^FOLR2^−^CD169^−^CD11c^−^, Lipid-associated Macrophages (LAM, C10): CD9^+^CD11c^+^CD226^−^CD163^−^CD169^−^, Adipose Tissue Macrophages-1 (ATM-1, C3): CD226^+^CD11c^+^CD163^−^CD169^−^, Dendritic Cells-1 (DC-1, C6): CD163^+^CD226^+^CD11c^+^CD163^+^CD169^−^. (E) Representative flow cytometry plots depicting the proportion of NAM, MP-1, VAM, AAM, LAM, ATM-2, and DC-1 populations in 3-month (n=4) and 21-month (n=5) old females. (F) Quantification of the proportion of NAM, MP-1, VAM, AAM, LAM, ATM-2, and DC-1in the VAT from young and aged females. Data is displayed as the mean +/− SEM and is representative of one independent experiment from 3-month (n=4) and 21-month-old (n=5) females. Statistical significance was determined via Student’s two-tailed t-test with α=0.05 and *, p < 0.05; **, p < 0.01, n.s. = not significant.

To orthogonally validate the age-related changes observed in our scRNAseq data set, we mined cell surface markers to build a panel (CD45, CD11b, CD11c, CD226, FOLR2, CD163, CD38, CD169, and CD9) and used it to investigate the VAM, ATM-2, AAM, NAM, DC-1, LAM, and MP-1 clusters via multiparametric flow cytometry (Clusters 0, 3, 4, 5, 6, 10, and 13) (**Fig. 2C-D, S3B**). Our scRNAseq data showed Cluster 5 (NAMs: CD169^+^CD11c^−^) was enriched for *Siglec1* (CD169) expression, but not *Itgax* (CD11c) expression (**Fig. 2C-D, S3B**). Cluster 13 (MP-1: CD169^+^CD11c^+^) also expressed *Siglec1*, but unlike Cluster 5, exhibited *Itgax* (CD11c) expression (**Fig. 2C-D, S3B**). Furthermore, Clusters 3 and 6 were both enriched for *Cd226* (CD226) expression. Cluster 6 (DC-1: CD169^−^CD226^+^CD163^+^CD11c^+^), but not Cluster 3 (ATM-2: CD169^−^ CD226^+^CD163^−^CD11c^+^) showed significant enrichment for *Cd163* (CD163) expression (**Fig. 2C-D, S3B**). Cluster 10 (LAMs: CD169^−^CD226^−^CD9^+^CD11c^+^) lacked CD226 and CD163 expression but was enriched for CD9 (*Cd9*) (**Fig. 2C-D, S3B**). Both Cluster 0 (VAMs: CD38^+^FOLR2^+^CD169^−^ CD226^−^CD11c^−^) and Cluster 4 (AAMs: CD38^+^FOLR2^−^CD169^−^CD226^−^ CD11c^−^) expressed *Cd38* (CD38) and *Folr2* (Folate Receptor Beta, FOLR2) (**Fig. 2C-D, S3B**). To validate the changes observed via scRNAseq, flow cytometric analysis of VAT showed a decrease in Cluster 0 (VAMs) and Cluster 6 (DC-1) with age (**Fig. 2E-F**). Furthermore, Cluster 3 (ATM-1) and Cluster 13 (MP-1) were not affected with age, while Cluster 10 (LAM) and Cluster 4 (AAM) increased with aging (**Fig. 2E-F**). Notably, Cluster 5 (NAM) decreased with age in females (**Fig. 2E-F**). Similar results, except for Cluster 10, were obtained when the data was normalized for cell number and grams of tissue (**Fig. S3C**).

Next, we analyzed the genes that were differentially expressed with age in each cluster (**Fig. S4**). We excluded Cluster 4 from this analysis since this cluster was virtually absent in the young condition. Analysis of all DEGs showed that there was a subset of shared genes in many of the clusters that were up- or downregulated by age in our dataset (**Fig. S4A, Table S4**). Genes that were upregulated with age in several of the clusters included *Cxcl13*, *Ccl8*, *Cd55*, *C3*, *Mmp9*, *Gda*, and *Ly6e*, while genes downregulated with age included *Hspa1a*, *Hsp1ab*, *Hpgd*, *Ccl24*, *Cx3cr1*, and *Apoe* (**Fig. S4A**). Interestingly, VAMs (Cluster 0) downregulated expression of key marker genes *Folr2*, *Lyve1*, and *Ednrb* (**Fig. S4B**). *Timd4,* though not significant after correcting for multiple comparisons, decreased 3-fold with age in VAMs, suggesting these macrophages may show a reduced capacity for lysosomal activation with age (**Fig. S4B & Table S4**). NAMs (Cluster 5) significantly upregulated genes involved in antigen presentation (*Cd74*, *H2-Eb2*), compliment signaling (*C4b*, *C6*), and the growth factor *Gdf3* (**Fig. S4B & Table S4**). Aged NAMs downregulated the expression of *Abca1*, which mediates the efflux of cholesterol and phospholipids, and the scavenger receptor *Colec12*, which internalizes and degrades oxidatively modified low-density lipoprotein (**Fig. S4B & Table S4**). Of note, both *Abca1* and *Colec12* have been shown to be important for proper myelin clearance/uptake by phagocytes ^27,28^. DC-1 (Cluster 6) and MP-2 (Cluster 8) showed a significant downregulation of several ribosomal protein genes, while LAMs (Cluster 10) and IAMs (Cluster 11) both downregulated *Pmepa1*, a negative regulator of TGF-beta signaling (**Fig. S4B & Table S4**). Consistent with the latter, aged LAMs also upregulated *Anxa1* expression, which acts to inhibit inflammation by suppressing phospholipase A2 (**Fig. S4B**).

In summary, these data show that ATMs undergo remodeling in response to aging and support the existence of a transcriptionally distinct CD38^+^ population (AAMs) in the VAT that expands significantly during aging ^29,30^.

### Adipose nerve-associated macrophages (NAMs) are enriched for CD169 and display unique morphology and localization

Since cellular senescence is linked to aging, we wanted to investigate whether we could detect senescent cell signatures at the single cell level in tissue resident macrophages. To investigate senescence, we used the published SenMayo gene set to calculate a gene set enrichment score (GSES) for each cell ^31^. Our analysis suggested NAMs (Cluster 5), and to a lesser extent AAMs (Cluster 4) and MP-1 (Cluster 13) were enriched for senescence/senescence-associated secretory phenotype (SASP) genes (**Fig. 3A**). Gene Ontology (GO) analysis of all clusters also suggested NAMs were enriched for cellular senescence genes, with NAMs also showing enrichment for genes involved in the cellular response to unfolded protein, phagolysosome assembly, myeloid cell differentiation, positive regulation of gliogenesis, regulation of fat cell differentiation, and type 2 immune response (**Fig. 3B**). Given these results, we sought to further characterize the nerve-associated macrophage (NAM) cluster. Analysis of DEGs for Cluster 5 showed that NAMs were indeed enriched for genes involved in beta-adrenergic signaling (*Adrb2, Arrb2, Maoa, Aldh2*), IL-10 signaling (*Il10rb, Il10*), sialic-acid modifying enzymes and binding proteins (*St3gal6, Neu1, Siglece, Siglec1, Siglech),* chemokines/chemoattractants (*Cxcl2, Ccl12, Cxcl12, Ccl2, Ccl4, Ccl3*) and various other genes enriched in NAMs from the lung and sciatic nerve such as *Csf1r*, *Ms4a7*, *Mgl2*, *Sdc4*, *Aoah*, *Xist*, *St3gal6*, *Tanc2*, and *Cdr2* (**Fig. 3C & Table S2**) ^3,12,13,20,21^. Our analysis also revealed that NAMs were enriched for several stress response genes (*Hspa1a*, *Hspa1b*, *Hspa8*, *Dnaja1, Dnajb1, Hsph1*, *Ier3*, *Ier5*, *Atf3*), mitochondrial genes (*mt-Nd3, mt-Cyb, mt-Atp6, mt-Nd2, mt-Nd4*), complement & complement receptors (*C4b, C5ar1, C5ar1, C5ar2*), genes involved in prostaglandin metabolism (*Hpgd*, *Hpgds*), and genes involved in iron binding/transport (*Trf, Cp, Slc40a1, Slco2b1*), suggesting a potential role for NAMs in the regulation of cellular stress, regulation of inflammation, and iron, which is component of myelin (**Fig. 3C, Table S2**). Of note, NAMs also showed enrichment for the pattern recognition receptor genes *Nlrp3* and *Tlr7*, suggesting NAMs may be better equipped to respond to cellular damage and/or neurotropic viruses (**Fig. 3C**). Interestingly, NAMs express leukocyte immunoglobulin-like receptor genes *Lilr4b* and *Lilra5*, the latter of which has been associated with late-onset sporadic Alzheimer’s disease (LOAD) (**Fig. 3C**) ^32^.

**Figure 3.**
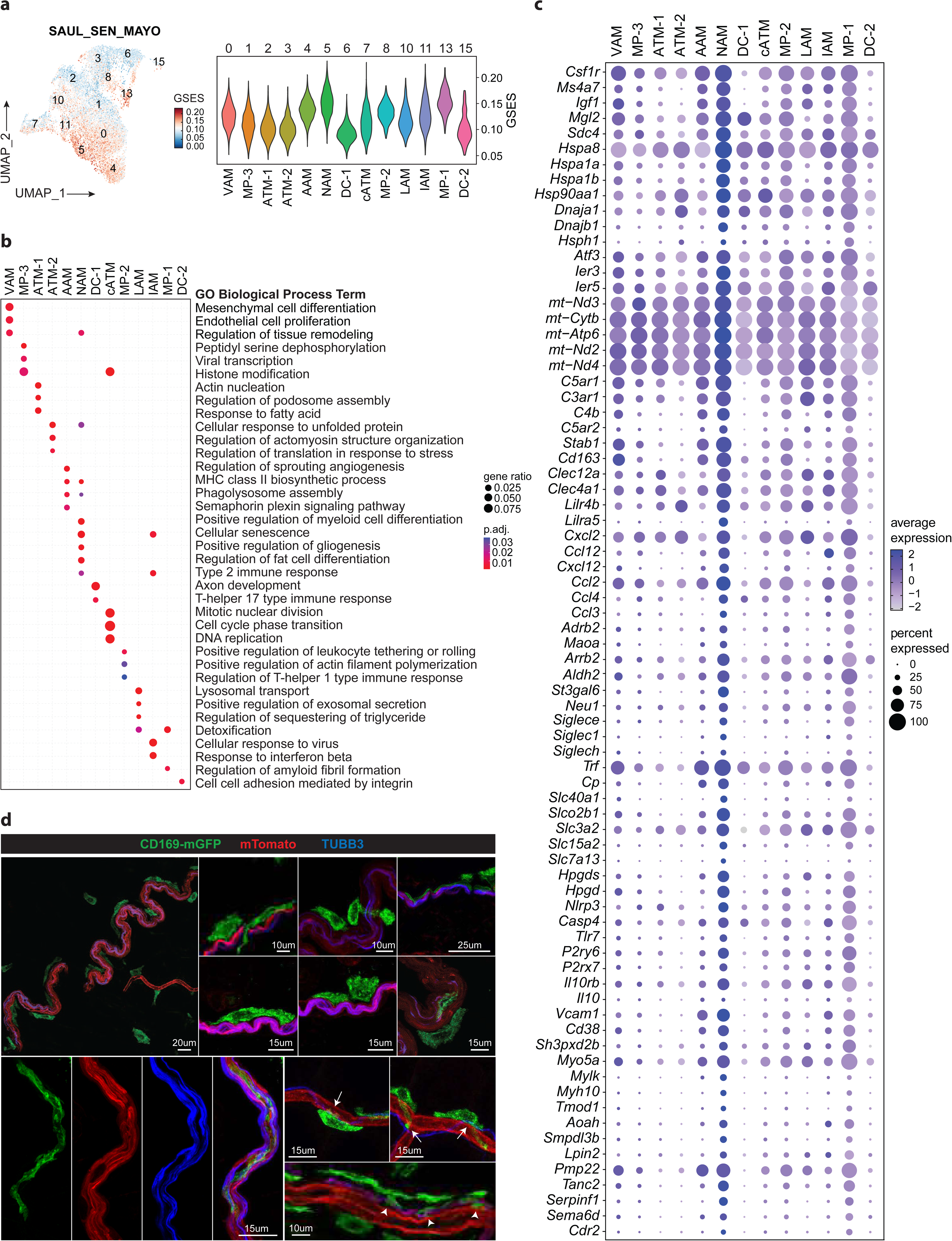
Adipose nerve-associated macrophages (NAMs) are characterized by CD169-expression and display distinct morphology and localization in the adipose tissue. (A) Uniform Manifold Approximation and Projection (UMAP) (*left*) and violin plot (*right*) showing enrichment of senescence associated gene transcripts and gene set enrichment score (GSES) per cell. Senescence associated gene transcripts were defined using the SenMayo gene set (SAUL_SEN_MAYO, MM16098). (B) Gene Ontology (GO) analysis showing enrichment of unique biological process (BP) terms by cluster. (C) Dot plot depicting the average expression of genes enriched in NAMs (Cluster 5). Dot size represents the percentage of cells expressing a corresponding gene in a given cluster. (D) Whole mount immunofluorescence confocal microscopy images of visceral adipose tissue (VAT) from adult CD169:mTmG dual reporter mice labeled with the pan-neuronal marker TUBB3. CD169 is labeled with membrane EGFP (mGFP^+^) (*green*) while non-recombined structures such as adipocytes, nerves, and blood vessels are labeled with membrane Tomato (mTomato^+^) (*red*). Non-recombined structures such as adipocytes, nerves, and blood vessels and can be distinguished by their morphology with adipocytes being round, nerves appearing dense and curved/sinusoidal, and blood vessels appearing straight and hollow. *Bottom Left*: Single channel and merged image of a CD169^+^ cell embedded between a mTomato^+^TUBB3^+^ nerve. *Bottom Right*: CD169^+^ cells on the surface of mTomato^+^ blood vessels and thin TUBB3^+^ nerve fibers. Data is representative of n = 4 independent experiments using 3- or 4-month-old females.

To examine the biology of NAMs further we performed *in vivo* two-photon microscopy using dual reporter LysM^Cre^:Rosa26^mT/mG^ mice (referred to here as LysM:mTmG). mT/mG is a cell membrane-targeted, two-color fluorescent Cre-reporter allele which only expresses membrane tdTomato (mT) in cells/tissues prior to Cre recombination. Following Cre recombination, the mT sequence is excised from Cre-expressing cells/tissues and is replaced with membrane EGFP (mG). This can be visualized as change of color from red (in non-recombined cells) to green ^33^. Using LysM:mTmG dual reporter mice, we were able to visually appreciate the diversity of adipose tissue myeloid cells (LysM:mGFP^+^) surrounding different adipose structures (mTomato^+^) such as adipocytes, blood vessels, and nerves that can easily be distinguished by their morphology, with adipocytes being round, nerves appearing dense and curved/sinusoidal, and vessels appearing hollow and straight. (**Video S1**). Via *in vivo* two-photon microscopy, we were able to visualize dynamic LysM:mGFP^+^ cells moving rapidly through the vasculature, LysM:mGFP^+^ cells moving on or near the vasculature, and slightly more elongated LysM:mGFP^+^ cells situated atop nerves that were largely static but surveille the nerves via their pseudopodia (**Video S1**). To characterize *Siglec1* or CD169^+^ NAMs more closely, we performed confocal, electron, and *in vivo* two-photon microscopy using CD169^Cre^:Rosa26^mT/mG^ dual reporter mice (referred to here as CD169:mTmG). Whole mount immunofluorescence (IF) imaging of the VAT from adult CD169:mTmG females stained with the pan-neuronal marker TUBB3 showed elongated CD169^+^ cells co-localizing on the surface of TUBB3^+^ nerve bundles and blood vessels (**Fig. 3D, Video S2**). Confocal imaging also showed elongated CD169^+^ NAMs embedded deep within TUBB3^+^ nerve bundles (**Fig. 3D, Video S2**). Notably, we frequently observed CD169^+^ NAMs with extended pseudopodia contacting thin and large nerve fibers and blood vessels (**Fig. 3D**). Higher resolution imaging revealed elongated CD169^+^ NAMs with pseudopodia wrapped around nerves while simultaneously contacting blood vessels (**Fig. 3D**). Moreover, serial high resolution imaging of VAT from CD169:mTmG reporter mice stained with DAPI showed multiple elongated CD169^+^ cells surrounding TUBB3^+^ nerves, suggesting NAMs may be involved in adipose nerve maintenance and/or support their survival (**Fig. 4A, Video S3**). Consistent with the imaging, our scRNAseq data showed NAMs (Cluster 5) were enriched for genes required for podosome formation, cell adhesion, and cell polarity/contractility (*Vcam1, Cd38, Sh3pxd2b, Myo5a, Mylk, Tmod1*) and the neurotropic factors (*Serpinf1, Sema6d*) (**Fig. 3C**). To examine these interactions in more detail, we performed transmission electron microscopy (TEM) of isolated nerves from the inguinal WAT (iWAT). Unlike the VAT, the large nerves from inguinal WAT (iWAT) can be isolated using a dissection microscope. The electron microscopy (EM) of isolated iWAT nerves showed classic nerve fascicles with endoneurium and perineurium, myelinated and unmyelinated axons, Schwann cells, and blood vessels (**Fig. 4B**). Magnification of the endoneurium showed elongated macrophages with extended pseudopodia surrounded by myelinated and unmyelinated nerve axons, suggesting these cells survey the nerve microenvironment (**Fig. 4C**). Intriguingly, EM of the iWAT nerves also showed macrophages with internalized electron dense material, likely myelin, suggesting NAMs may play a role in myelin clearance (**Fig. 4D**). Consistent with this hypothesis, our scRNAseq analysis showed NAMs were enriched for the gene *Pmp22*, which is a major component of myelin in the peripheral nervous system (PNS) (**Fig. 3C**). To investigate these findings further, we conducted whole mount confocal imaging of iWAT and VAT stained with TUBB3 and the myelin markers MBP or CNPase, which are major constituents of the myelin sheath in Schwann cells, the glial cells that form the myelin sheath in the PNS. The high resolution imaging confirmed the presence of CD169^+^ NAMs on myelinated nerve bundles and deeply embedded between myelinated nerve fibers in iWAT (**Fig. 4E**). Furthermore, in the VAT we observed co-localization of CD169^+^ cells with MBP^+^ particles adjacent to myelinated nerves, supporting a role for CD169^+^ NAMs in myelin uptake/clearance and/or in the maintenance of myelinating Schwann cells (**Fig. 4F, Video S4 & S5**).

**Figure 4.**
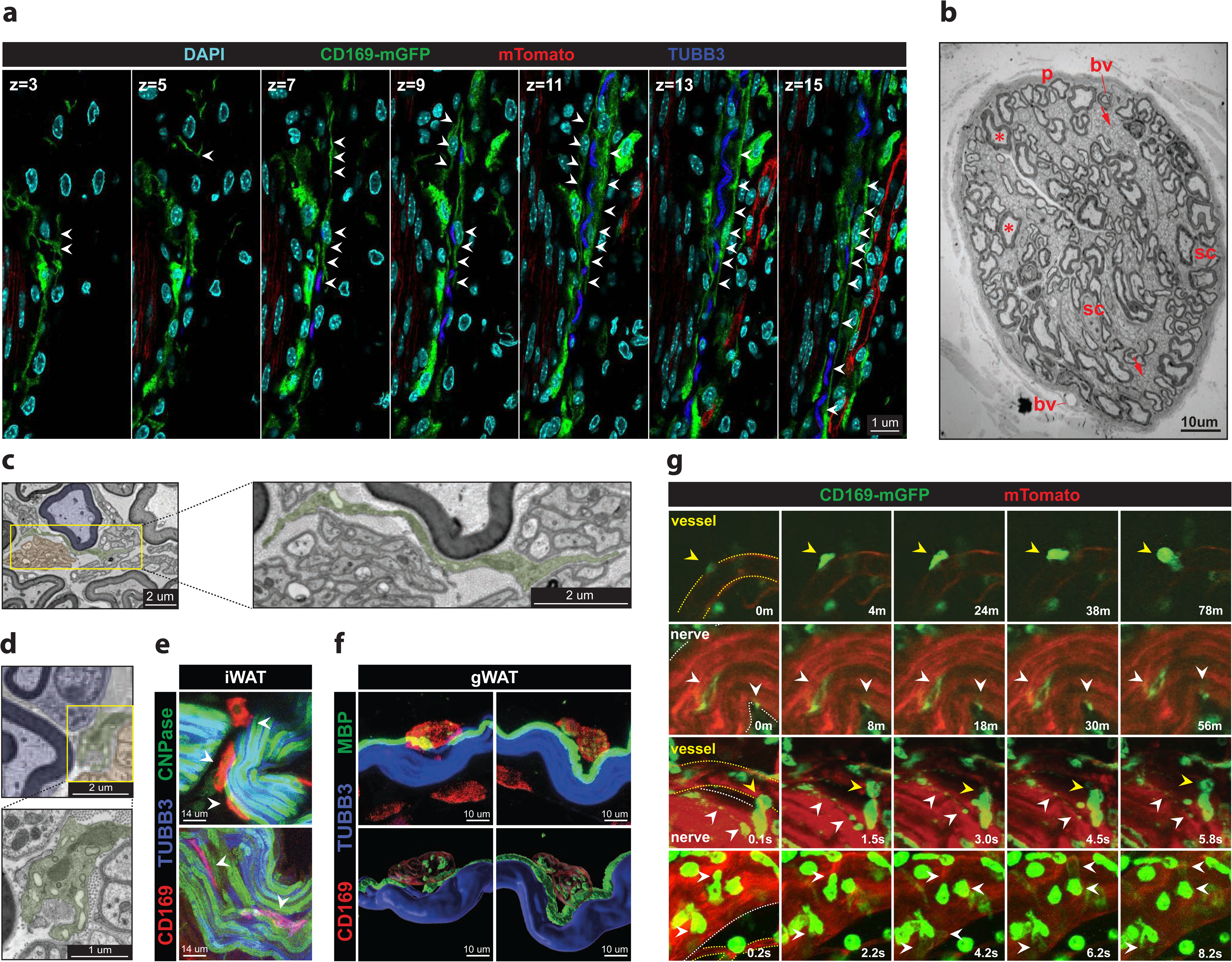
CD169+ nerve-associated macrophages (NAMs) engulf myelin and can surveil the adipose nerve microenvironment. (A) Whole mount immunofluorescence of VAT from adult CD169:mTmG dual reporter mice labeling the nucleus (*cyan*), the pan-neuronal marker TUBB3 (*blue*), CD169+ macrophages with membrane EGFP (mGFP+) (*green*), and non-recombined structures such as adipocytes, nerves, and blood vessels with membrane Tomato (mTomato+) (*red*). Non-recombined structures such as adipocytes, nerves, and blood vessels can be distinguished by their morphology with adipocytes being round, nerves appearing dense and curved/sinusoidal, and blood vessels appearing straight and hollow. Each panel represents a single plane on the z-axis (z). White arrowheads highlight elongated CD169+ cells. Data is representative of n = 4 independent experiments using 3-4 month-old females. (B) Transmission electron microscopy (TEM) of a nerve bundle isolated from the inguinal white adipose tissue (iWAT). The following structures are labeled: perineurium (*p*), Schwann cells (*sc*), myelinated axons (*\**), unmyelinated axons (*black arrow*), blood vessels (*bv*). Data is representative of n = 2 independent experiments using 3-month-old wildtype males. (C) Transmission electron microscopy (TEM) of a nerve bundle isolated from the inguinal white adipose tissue (iWAT) depicting an elongated nerve-associated macrophage (NAM) (*green*) located between myelinated (*blue*) and unmyelinated (*orange*) axons. Data is representative of n = 2 independent experiments using 3-month-old wildtype males. (D) Transmission electron microscopy (TEM) of a nerve bundle isolated from the inguinal white adipose tissue (iWAT) depicting a nerve-associated macrophage (NAM) (*green*) with pseudopodia and intracellular electron dense material located between myelinated (*blue*) and unmyelinated (*orange*) axons. Data is representative of n = 2 independent experiments using 3-month-old wildtype males. (E) Whole mount immunofluorescence of iWAT labeled with antibodies against the NAM marker (*red*, *white arrowheads*), the pan-neuronal marker TUBB3 (*blue*), and the myelin marker CNPase (2′,3′-Cyclic-nucleotide 3’-phosphodiesterase) (*green*). Data is representative of n = 1 independent experiment using 3-month-old wildtype males (n = 2). (F) Whole mount immunofluorescence of VAT from adult wildtype males labeled with antibodies against the NAM marker CD169 (*red*, *white arrowheads*), the pan-neuronal marker TUBB3 (*blue*), and the myelin marker MBP (myelin basic protein) (*green*). Data is representative of n = 1 independent experiment using 3-month-old wildtype males (n = 2). (G) *In vivo* two-photon imaging of iWAT from adult CD169:mTmG dual reporter mice labeling CD169^+^ macrophages with membrane EGFP (mGFP^+^) (*green*), and non-recombined structures such as adipocytes, nerves, and blood vessels with membrane Tomato (mTomato^+^) (*red*). Non-recombined structures such as adipocytes, nerves (*white outline*), and blood vessels (*yellow outline*) and can be distinguished by their morphology with adipocytes being round, nerves appearing dense and curved/sinusoidal, and blood vessels appearing straight and hollow. Yellow arrowheads highlight CD169^+^ cells interacting with blood vessels and/or nerves, white arrowheads highlight CD169^+^ cells on large nerve bundles. Data is representative of n = 3 independent experiment using 3-month-old males

Next, we performed *in vivo* two-photon microscopy to assess the dynamics of CD169^+^ NAMs in iWAT using CD169:mTmG dual reporter mice. We imaged large nerve bundles, as small thin fibers that normally run along blood vessels are not strongly labeled by mTomato (**Fig. 3D**). Visualization of CD169^+^ NAMs in the iWAT revealed motile CD169^+^ cells on blood vessels, and non-motile CD169^+^ cells on large nerve bundles (**Fig. 4G, Video S6-S7**). Despite being fairly static, live imaging revealed that CD169^+^ NAMs extend their dendrites, suggesting they are actively surveilling the surrounding nerve microenvironment (**Fig. 4G, Video S6**). Moreover, CD169^+^ cells on nerves could be seen forming balloon-like protrusions on the surface of the nerve, which may represent active phagocytosis of nerve cellular components (**Fig. 4G, Video S7**).

In summary, these data demonstrate that CD169^+^ NAMs have diverse morphology and can be observed as elongated cells that contact nerves and blood vessels via their extended pseudopodia, and support a role for adipose CD169^+^ NAMs in nerve surveillance and/or maintenance of adipose tissue nerve homeostasis.

### CD169^+^ nerve-associated macrophages (NAMs) are necessary for preventing catecholamine resistance and adipose tissue dysfunction

We next sought to understand the functional role of CD169^+^ NAMs in the adipose tissue *in vivo*. To do this we utilized CD169 diphtheria toxin receptor (CD169-DTR) mice to deplete CD169^+^ macrophages in the VAT of 3-month-old wildtype (WT) or CD169-DTR female mice treated with diphtheria toxin (DT) for 12-days (**Fig. 5A**) ^34^. Depletion of NAMs in VAT was confirmed via flow cytometry on day 13 (D13) (**Fig. 5B**). Depletion of NAMs led to a 5% reduction in body weight by day 3 that gradually increased and was equivalent to control mice after day 6 (**Fig. S6A**). Consistent with the latter observation, VAT weight was not significantly different between groups at D13, but there was an increase in the number of total stromal vascular fraction (SVF) cells from the VAT of CD169-DTR mice (**Fig. 5C**). Flow cytometry analysis of SVF from VAT showed an increase in the proportion of live CD45^+^ cells and decrease in live CD45^−^ cells (**Fig. 5D, S5**). There were no significant changes in the proportion of B-cells, T-cells, neutrophils, eosinophils, and MHCII^+^CD11c^+^ cell populations, however there was a 2-fold increase in the proportion of Ly6C^+^ monocytes in the adipose tissue, most likely in response to a reduction in tissue resident NAMs (**Fig. 5D, S5**). Unlike the VAT, in the spleen, the tissue weight and total number of splenocytes did not change significantly following depletion of CD169^+^ cells, and the increase in Ly6C^+^ monocytes was negligible (**Fig. S6B-E**).

**Figure 5.**
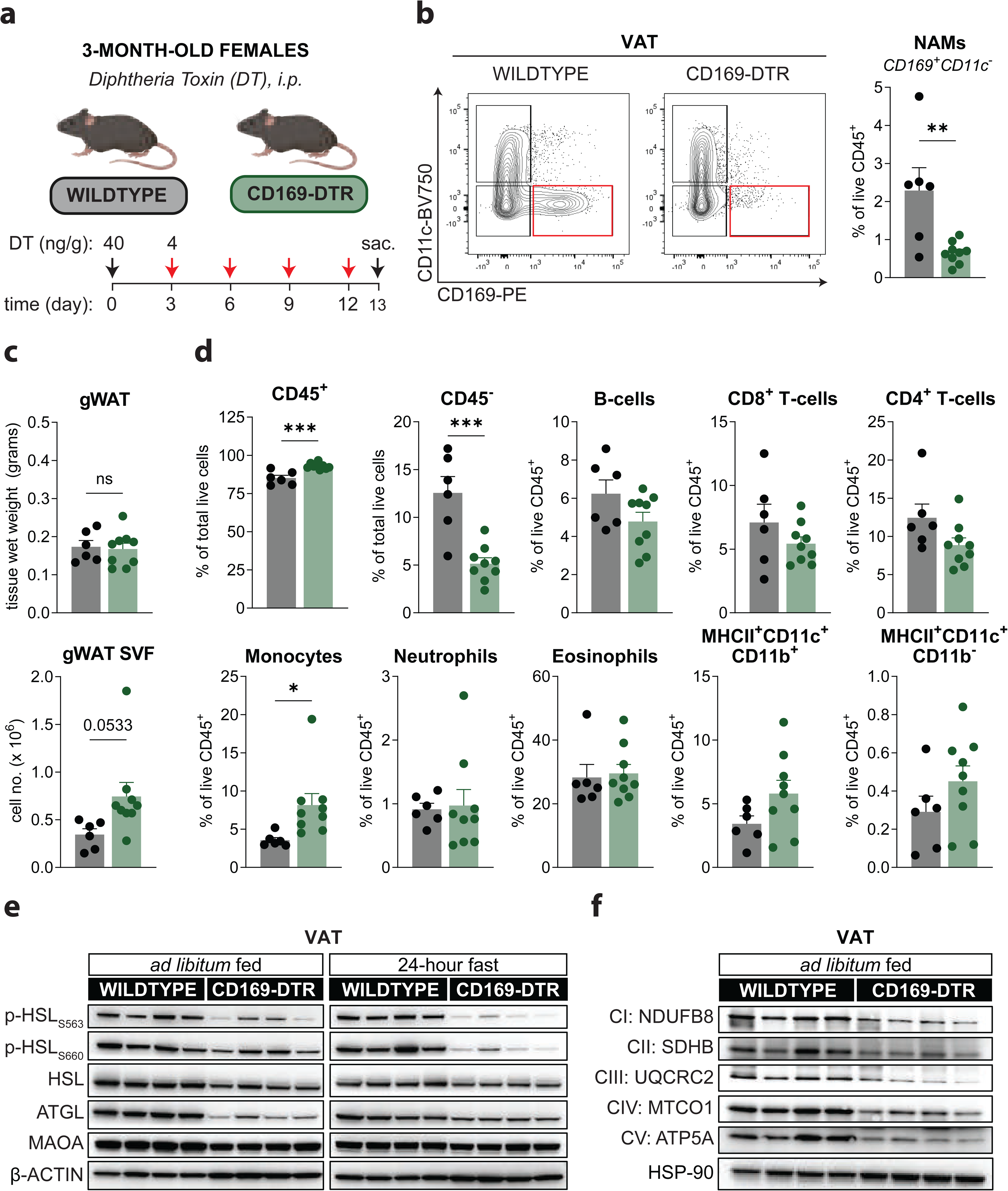
CD169^+^ nerve-associated macrophages (NAMs) are necessary for preventing catecholamine resistance and adipose tissue dysfunction. (A) Schematic representation of the experimental protocol used to deplete CD169-macrophages using CD169 diphtheria toxin receptor (CD169-DTR) mice. 3-month-old wildtype (WT) (n=6) and CD169-DTR (n=9) female mice were injected intra-peritoneal (i.p) with 40ng/gram diphtheria toxin (DT) on day 0 (D0) followed by 4ng/gram DT on D3, D6, D9, and D12, and sacrificed on D13. (B) Representative flow cytometry plots (*left*) and bar graph (*right*) quantifying nerve-associated macrophages in the stromal vascular fraction (SVF) of VAT from 3-month-old WT and CD169-DTR females treated with DT and sacrificed on D13. Data is represented as a mean +/− SEM. Statistical significance was determined via Student’s two-tailed t-test with α=0.05 and p < 0.05. (C) Tissue wet weight (*top*) and stromal vascular fraction (SVF) cell number (*bottom*) from the VAT of from 3-month-old WT and CD169-DTR females treated with DT and sacrificed on D13. Data is represented as a mean +/− SEM. Statistical significance was determined via Student’s two-tailed t-test with α=0.05 and p < 0.05. (D) Flow cytometry from the VAT of 3-month-old WT and CD169-DTR females treated with DT and sacrificed on D13. Bar plots represent either the proportion of CD45+ and CD45-cells (as a fraction of total live cells) and B-cells, CD8^+^ T-cells, CD4^+^ T-cells, Monocytes, Neutrophils, Eosinophils, and MHCII+CD11c+ ATMs or dendritic cells (DCs) (as a fraction of live CD45+). Populations were defined using the cell surface markers and gating strategy outlined in Supplementary Figure 5. Data is represented as a mean +/− SEM. Statistical significance was determined via Student’s two-tailed t-test with α=0.05 and p < 0.05. (E) Total expression and phosphorylation levels of lipolysis proteins from the VAT of *ad libitum* fed or 24-hour *fasted* 9-month-old WT and CD169-DTR females treated with DT and sacrificed on D8 (n=4 per genotype). (F) Total expression of electron transport chain proteins from the VAT of *ad libitum* fed or 24-hour *fasted* 9-month-old WT and CD169-DTR females treated with DT and sacrificed on D8 (n=4 per genotype).

Previously, we and others showed that NAMs can regulate norepinephrine bioavailability and may have pro-lipolytic and anti-obesogenic effects ^3,12^. To investigate the role for CD169^+^ NAMs in lipolysis we studied two enzymes that regulate lipolysis, HSL and ATGL, in the VAT of adult 9-month-old WT and CD169-DTR females sacrificed on D8 (**Fig. S7A-B**). Immunoblot analysis of VAT from *ad libitum* fed and 24-hour *fasted* mice showed depletion of CD169^+^ NAMs led to a reduction in monoamine oxidase A (MAOA) protein levels (**Fig. 5E, S7C**). Interestingly, depletion of CD169^+^ NAMs also resulted in a reduction of total ATGL and HSL in the fed and fasted conditions, but levels of phosphorylated HSL (p-HSL) at both serine 563 (S563) and serine 660 (S660) were significantly reduced in the 24-hour fasted condition (**Fig. 5E, S7C**). Furthermore, expression levels of the β3-adrenergic receptor (*Adrb3*), which acts upstream of ATGL and HSL during catecholamine-induced lipolysis, was also reduced in CD169-DTR mice (**Fig. S6F**). Together, these data suggested that a reduction in adipose NAMs and consequential lowered MAOA may promote catecholamine resistance by increasing catecholamine availability. The increase in catecholamines likely activates a negative feedback mechanism to downregulate the expression of lipolytic proteins. Consistent with the above, we also observed decreased expression of the master regulator of adipocyte differentiation, *Pparγ*, suggesting there is an overall dysregulation of lipid metabolism following NAM depletion (**Fig. S6F**). Lastly, we also observed a reduction in the expression of oxidative phosphorylation proteins in the VAT of CD169-DTR mice, suggesting loss of NAMs may also alter mitochondrial function (**Fig. 5F, S7D**). Together, our results confirm that NAMs are critical for regulating catecholamine signaling and suggest that NAMs may be required for preventing catecholamine resistance and are required for maintenance of adipose tissue function.

### Depletion of CD169^+^ NAMs during aging impair adipose tissue metabolism and increase inflammation

Next, to determine the functional role of CD169^+^ NAMs through lifespan, we aged the CD169-DTR mice up to 24-months and evaluated the animals 24-days post NAM depletion (**Fig. 6A**). Consistent with our previous data, 6m CD169-DTR females lost approximately 5% of their body weight by day 3 but regained it by day 12 and maintained their body weight until day 24 (**Fig. 6B**). On the contrary, 24mo CD169-DTR females lost approximately 15% of their bodyweight by day 24 and had smaller fat pads and more SVF cells compared to controls (**Fig. 6C, S8A**). To investigate whether the reduction in adipose tissue weight was due to changes in lipolysis, we measured serum free fatty acid (FFA) levels on D24 and found that young 6m-old CD169-DTR females had similar serum FFA levels, but 24m-old females had lower FFA levels relative to wildtype controls (**Fig. 6D**). In addition, immunoblot analysis of VAT from 24m WT and CD169-DTR females showed that ablation of CD169^+^ NAMs also reduced total MAOA, ATGL, and phosphorylated HSL (S563 and S660) protein levels, but not total HSL (**Fig. 6E, S8B**). Moreover, GDF3 (*Gdf3*) which can inhibit catecholamine-induced lipolysis, was increased 20-fold in the VAT following depletion of NAMs in 24m but not 6m CD169-DTR females ^3,35,36^ (**Fig. 6F**). Expression of *Adrb3* on the other hand was lower between genotypes but did not reach statistical significance (**Fig. 6F**). Consistent with our data in young females, loss of CD169^+^ NAMs in aging also led to reduction in the expression of *Pparγ* in the VAT (**Fig. 6F, S8B**). Together, these data confirm that CD169^+^ NAMs are required for proper regulation of the lipolytic program and suggest that the weight loss observed in aged animals may be dependent on GDF3.

**Figure 6.**
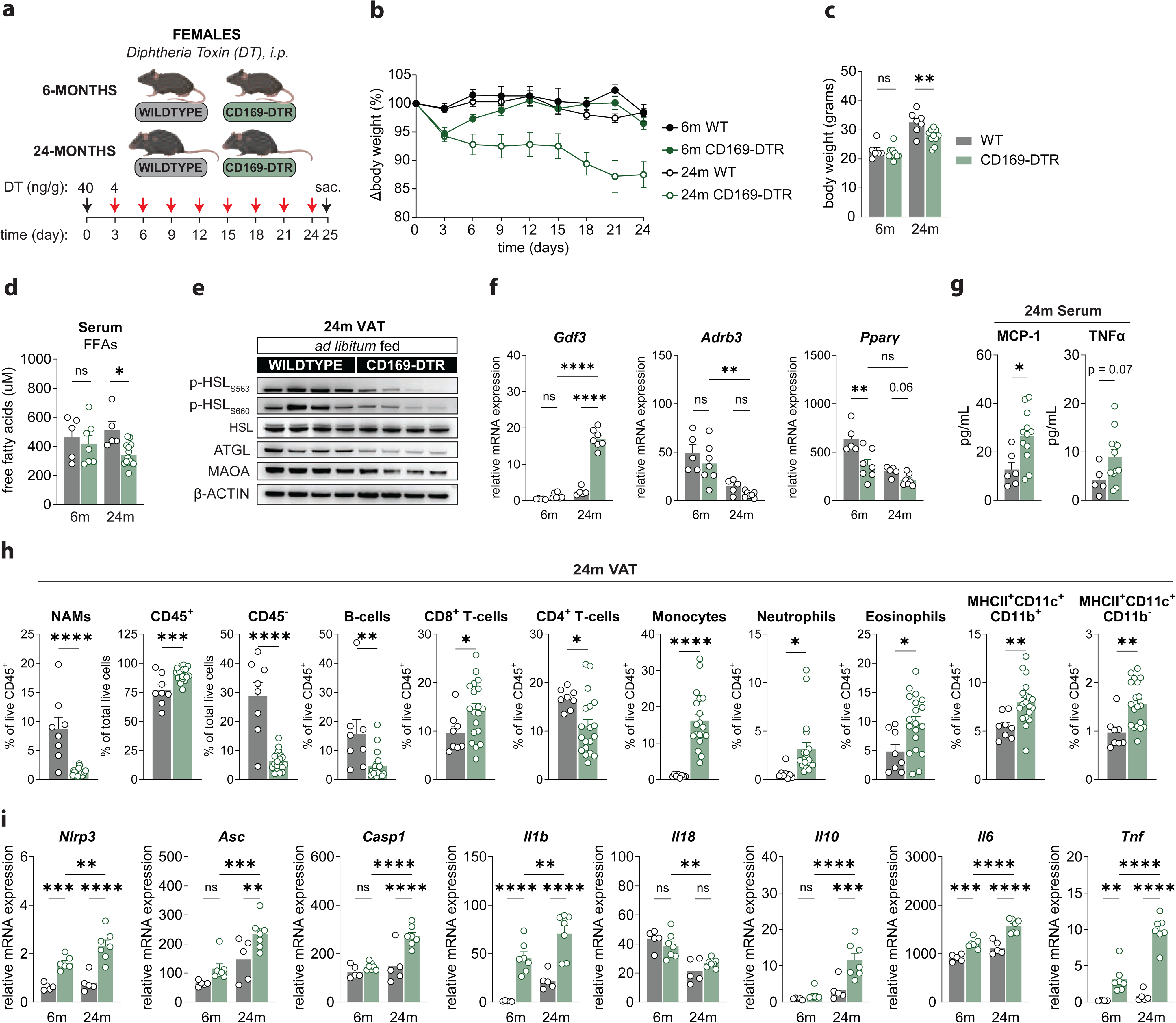
CD169^+^ nerve-associated macrophages (NAMs) restrain age-induced inflammation in the adipose tissue. (A) Schematic representation of the experimental protocol used to deplete CD169-macrophages using CD169 diphtheria toxin receptor (CD169-DTR) mice. 6-month and 24-month-old wildtype (WT) and CD169-DTR female mice were injected intra-peritoneal (i.p) with 40ng/gram diphtheria toxin (DT) on day 0 (D0) followed by 4ng/gram DT on D3, D6, D9, and D12, D15, D18, D21, D24, and sacrificed on D25. (B) Change in body weight curves for 6-month and 24-month-old WT and CD169-DTR females treated with DT and sacrificed on D24. (C) Bar graph of body weight for 6-month and 24-month-old WT and CD169-DTR females (n=6-11 per genotype) treated with DT and sacrificed on D24. Data is represented as a mean +/− SEM. Statistical significance was determined via two-way ANOVA with α=0.05 and p < 0.05. (D) Quantification of free fatty acids (FFA) in the serum of 6-month and 24-month-old WT and CD169-DTR females (n=5-15 per genotype) treated with DT and sacrificed on D24. Data is represented as a mean +/− SEM. Statistical significance was determined via two-way ANOVA with α=0.05 and p < 0.05. (E) Total expression and phosphorylation levels of lipolysis proteins from the VAT of *ad libitum* 24-month-old WT and CD169-DTR females treated with DT and sacrificed on D25 (n=4 per genotype). (F) Relative mRNA expression of *Gdf3*, *Adrb3*, and *Pparγ* genes in the VAT of *ad libitum* fed 6- and 24-month-old WT and CD169-DTR females treated with DT and sacrificed on D25 (n=5-7 per genotype). Data is represented as a mean +/− SEM. Statistical significance was determined via two-way ANOVA with α=0.05 and p < 0.05. (G) Monocyte chemoattractant protein-1 (MCP-1) and TNFα levels in the serum of *ad libitum* fed 24-month-old WT and CD169-DTR females treated with DT and sacrificed on D25 (n=5-7 per genotype). Data is represented as a mean +/− SEM. Statistical significance was determined via Student’s two-tailed t-test with α=0.05 and p < 0.05. (H) Flow cytometry from the VAT of 24-month-old WT and CD169-DTR females (n=8-19 per genotype) treated with DT and sacrificed on D25. Bar plots represent either the proportion of CD45^+^ and CD45^−^ cells (as a fraction of total live cells) and B-cells, CD8^+^ T-cells, CD4^+^ T-cells, Monocytes, Neutrophils, Eosinophils, and MHCII^+^CD11c^+^ ATMs or dendritic cells (DCs) (as a fraction of live CD45^+^). Populations were defined using the cell surface markers and gating strategy outlined in Supplementary Figure 5. Data is represented as a mean +/− SEM. Statistical significance was determined via Student’s two-tailed t-test with α=0.05 and p < 0.05. Data is represented as a mean +/− SEM. Statistical significance was determined via Student’s two-tailed t-test with α=0.05 and p < 0.05. (I) Relative mRNA expression of inflammasome and pro-inflammatory cytokine genes in the VAT of *ad libitum* fed 6- and 24-month-old WT and CD169-DTR females treated with DT and sacrificed on D25 (n=5-6 per genotype). Data is represented as a mean +/− SEM. Statistical significance was determined via two-way ANOVA with α=0.05 and p < 0.05.

Since chronic inflammation is considered a hallmark feature of the aging process and is directly linked to a decline in tissue function, we next assessed if loss of CD169^+^ NAMs alters age-induced inflammation. The analysis of several cytokines in the serum of 24m females showed there was an elevation in monocyte chemoattractant protein-1 (MCP-1) and TNFα levels in CD169-DTR mice, but no changes in systemic levels of IL-1β or IL-6 (**Fig. 6G, S8C**). Multi-parameter flow cytometry analyses revealed that unlike in adult VAT, depletion of NAMs led to a decrease in B-cells in aged VAT (**Fig. 6H**). Assessment of T-cell subsets also showed there was an overall reduction in CD4^+^ T-cells and increase in CD8^+^ T-cells (**Fig. 6H**). Moreover, NAM depletion in aged VAT was coupled with a 16-fold increase in monocytes, 5-fold increase in neutrophils, 2-fold increase in eosinophils, and an increase in MHCII^+^CD11c^+^ ATMs/DCs (**Fig. 6H**). We observed similar changes in 24m VAT from mice treated with DT for 34-days (**Fig. S8D**). Assessment of specific CD4^+^ T-cell subsets showed regulatory T-cells (T-regs), gamma delta T-cells (gd T-cells), and Type 2 helper T-cells (T_H2_ cells) were reduced following NAM depletion, suggesting a decrease in CD169^+^-macrophages could alter self-tolerance mechanisms or the ability to clear extracellular pathogens (**Fig. S8E**). In comparison, we did not observe changes in tissue weight or the B-cell and T-cell subsets in the spleen, but there was a 2-fold increase in monocytes, neutrophils, and eosinophils (**Fig. S9A-C**). Together, these data demonstrated that a reduction in NAMs during aging drastically alters the immune cell landscape in VAT, but not spleen, and suggests that a reduction of NAMs over time may potentiate age-induced inflammation.

Consistent with the latter hypothesis, elimination of CD169^+^ NAMs in aged mice significantly increased expression of pro-inflammatory cytokines *Il1b*, *Il6*, and *Tnf* (**Fig. 6I**). Interestingly, *Il10*, which is generally considered an anti-inflammatory, was elevated upon NAM depletion in aged mice without affecting *Il18 (***Fig. 6I**). Given NLRP3 inflammasome activation induces age-related inflammation and impairs metabolic function in aging^2^, we also assessed the expression of adaptor protein, ASC (*Asc*), and the main effector enzyme, Caspase-1 (*Casp1*). Interestingly, we found that both *Asc* and *Casp1* were significantly increased only in the VAT of aged CD169-DTR mice (**Fig. 6I**). Together, these data show that CD169^+^ NAMs are anti-inflammatory and are necessary for restraining age-induced inflammation in the adipose tissue.

## Discussion

In this study, we utilized an iv-labeling approach to exclude circulating and enrich for non-circulating F4/80^+^CD11b^+^ cells, and performed unbiased scRNAseq to generate unprecedented insight into the transcriptional diversity of ATMs in the visceral adipose tissue with age. Our study validated the transcriptional signatures associated with two ATM sub-clusters: VAMs (vessel-associated macrophages) and LAMs (lipid-associated macrophages). Importantly, our study uncovered previously unknown transcriptional signatures of two new macrophage subtype: AAMs (age-associated macrophages) and NAMs (nerve-associated macrophages). Using our scRNAseq dataset, we identified that the NAM sub-cluster in VAT is enriched for *Siglec1* (CD169) as well as genes previously associated with in cellular senescence. Furthermore, we show that CD169^+^ NAMs decrease with age in the VAT and provide imaging evidence showing this ATM-subset may be important for the regulation of myelin uptake and/or processing. Finally, using CD169-DTR mice to deplete NAMs, we demonstrate that CD169^+^ NAMs are functionally important for regulating lipolysis, restraining age-induced inflammation, and preventing adipose tissue dysfunction in the VAT at homeostasis and during aging.

Our study supports a model whereby a reduction in adipose CD169^+^ NAMs may drive or partially explain the reduction in lipolysis and increase in visceral adiposity observed during aging. This hypothesis is supported by our data showing that CD169^+^ NAMs decrease with age in the VAT and that a reduction in NAMs in both young and aged CD169-DTR mice blunts lipolysis protein expression. However, it remains unclear if a similar mechanism (i.e. reduction in NAMs) might also play a role in diet-induced obesity. For example, one study demonstrated that ablation of CD169^+^ macrophages using CD169-DTR mice fed a chow or high-fat diet (HFD) resulted in adipocyte hypertrophy in VAT, a result that would be consistent with our current hypothesis ^37^. A separate study that did not use CD169 showed that F4/80^+^CD45^+^ NAMs accumulate on sympathetic nerves from the iWAT of HFD-fed mice, but it is unknown if similar changes also occur in the VAT ^12^. The latter study also noted an accumulation of CD11c^+^ macrophages on sympathetic adipose nerves from obese mice, suggesting that the increase may also be due to infiltrating CD169^−^CD11c^+^ cells that express F4/80 on their surface. Thus, it remains unclear what role NAMs play under distinct energy states (i.e., lean and obese) or in distinct fat depots (i.e., VAT and iWAT).

Here we discovered that CD169^+^ macrophage subsets preferentially associate with nerves in the adipose tissue, consistent with another study showing CD169 (i.e. Siglec-1 or Sn) can be used to identify NAMs in the lung ^13^. We also provide evidence that CD169^+^ NAMs can simultaneously contact both nerves and blood vessels suggesting they likely participate in the maintenance of both structures. In line with these observations, our scRNAseq indicated NAMs (Cluster 5) are enriched for *Vcam1* expression, which mediates the adhesion of cells to the vascular endothelium. However, the role of CD169^+^ cells in the peripheral nervous system (PNS) remains less well-defined. Our whole mount confocal imaging of adipose tissue found that CD169^+^ NAMs can engulf myelin, supporting a role for NAMs in myelin uptake and/or processing. Our analysis also showed that Cluster 5 NAMs are enriched for genes involved in gliogenesis, the process of generating glial cells, such as myelinating Schwann cells found in the PNS. Intriguingly, two studies using genetic models of peripheral and central nervous system demyelination have shown that CD169-deficient mice exhibited reduced demyelination and infiltration of CD8^+^ T-cells and macrophages ^38,39^. Hence, future studies are needed to delineate if CD169^+^ NAMs have defined roles in Schwann cell generation, maintenance, and/or nerve demyelination. Lastly, it’s important to note that use of CD169-DTR mice results in global depletion of CD169^+^ cells. Thus, the development of new conditional knockout mouse models to target NAMs will be critical for understanding their role in the PNS.

Notably, our scRNAseq study identified a population of macrophages in the VAT, which we termed AAMs, that was virtually absent in the young condition. Using flow cytometry we confirmed that AAMs increase with age in the VAT. We found that AAMs were enriched for expression of genes encoding several secreted proteins that have been associated with the senescence-associated secretory phenotype (SASP), such as: C3, CD55, CCL8 and MMP9. Our data suggests that accumulation of AAMs may promote age-related dysfunction. In support of this hypothesis, we showed that AAMs can be defined by expression of the cell surface marker CD38 – a marker previously linked to a subset of macrophages that reduces NAD^+^ levels and drives functional decline in aging ^29,30^. Furthermore, AAMs were also enriched for *Cxcl13* expression, which is involved in the formation of fat associated lymphoid clusters (FALCs) that accumulate in the adipose tissue during aging ^40^. These data support a role for CD38^+^ macrophages in aging, however, future studies will be required to characterize whether this ATM subpopulation displays other canonical markers and features associated with senescent cells.

In summary, our results transcriptionally identify CD169^+^ macrophages as an ATM subpopulation in the VAT, NAMs, and identify a role for these ATMs in catecholamine-induced lipolysis and age-induced inflammation. Furthermore, we identify a novel ATM subpopulation, AAMs, which may play a role in driving age-related dysfunction.

## Methods

### Mice

2- and 20-month-old C57BL/6J wildtype males and females were obtained from the National Institute of Aging (NIA) Aged Rodent Colony and housed in specific pathogen-free facilities at Yale University. CD169^Cre^:Rosa26^mTmG^ (CD169:mTmG) and LysM^Cre^:Rosa26^mTmG^ (LysM:mTmG) reporter mice were generated in our facility by crossing CD169^Cre^ mice (kindly provided by Dr K.M. Khanna, NYU) or LysM^Cre^ mice to Rosa26^mTmG^ mice (The Jackson Laboratory). Siglec1^DTR/+^ (CD169-DTR) mice were generated by Dr K.M. Khanna in NYU Langone Health. Experiments using CD169-DTR mice were conducted in adult females (3-, 6-, and 9-months-old) or aged females (24-months-old). Animal experiments and animal use was conducted in compliance with the National Institute of Health Guidelines for the Care and Use of Laboratory Animals and was approved by the Institutional Care and Use Committee at Yale University.

### Diphtheria Toxin Administration

Diphtheria toxin (DT, Sigma Aldrich, St. Louis, MO, USA) was prepared in endotoxin free PBS and administered intraperitoneal (i.p.) on day 1 (40 ng/g body weight) and then every 2-3 days (4 ng/g body weight) until mice were sacrificed.

### Adipose Tissue Digestion and Flow Cytometry

Visceral adipose tissue (VAT) was digested in 0.8mg/mL collagenase II (Worthington Biochemicals), 3% BSA (Sigma), 1.2mM Calcium Chloride, 1.0mM Magnesium Chloride, 0.8mM Zinc Chloride, and 1x HBSS (Life Technologies 14185-052) for 40-45 minutes in a 37°C water bath with vigorous shaking. The Stromal Vascular Fraction (SVF) was filtered on a 100μm cell strainer. Red blood cell lysis was performed using ACK lysis buffer, and cells were filtered a second time with a 70μm cell strainer. One million SVF cells were used for flow cytometry staining. Cells were stained with live/dead Aqua viability dye (Invitrogen) or Viakrome 808 viability dye (Beckman Coulter) for 30 minutes, followed by Fc receptor blocking (Invitrogen) for 10 minutes and surface staining for 30 minutes with the following antibodies: CD45-PE-Cy7 (Clone 30-F11, catalog no. 103114), CD11b-APC (Clone M1/70 catalog no. 101212), FOLR2-PE (Clone 10/FR2 catalog no. 153304), CD9-FITC (Clone MZ3 catalog no. 124808), CD226-BV421 (Clone TX42.1 catalog no. 133615), CD11c-BV750 (Clone N418 catalog no. 117357), B220-BV711 (Clone RA3-6B2 catalog no. 103255), Ly6G-FITC (Clone 1A8 catalog no. 127606), Ly6C-AF700 (Clone HK1.4 catalog no. 128024) (**all from BioLegend**); CD45.2-FITC (Clone 104 catalog no. 11-0454-85), F4/80-PE-Texas Red (Clone BM8 catalog no. MF48017), CD163-SB702 (Clone TNKUPJ catalog no. 67-1631-82), CD169-APC-eFluor780 (Clone SER-4 catalog no. 47-5755-82), CD45-APC (Clone 30-F11, catalog no. 17-0451-83), CD169-PE (Clone SER-4 catalog no. 12-5755-80), FOXP3-PE-Cy5.5 (Clone FJK-16s catalog no. 35-5773-82), LYVE-1-Biotin (Clone ALY7 catalog no. 13-0443-82) (**all from Thermo Fisher Scientific**); CD38-BUV737 (Clone 90/CD38 catalog no. 741748), Siglec-F-BV605 (Clone E50-2440 catalog no. 740388), CD11b-BUV395 (Clone M1/70 catalog no. 563553), CD3-BV480 (Clone 17A2 catalog no. 565642), CD4-BUV563 (Clone GK1.5 catalog no. 612923), CD8a-BUV615 (Clone 53-6.7 catalog no. 613004), I-A/I-E-BUV805 (Clone M5/114.15.2 catalog no. 748844), CD25-PE-CF594 (Clone PC61 catalog no. 562694) (**all from BD Biosciences**); and F4/80-StarBrightViolet570 (Clone CI:A3-1 catalog no. MCA497) (from Bio-Rad). Flow cytometry analysis was performed on a Becton Dickinson Symphony instrument, and data were analyzed using FlowJo software (Treestar).

### RNA Extraction, cDNA Generation, and RT-qPCR

Total RNA was isolated using Trizol followed by the RNeasy Kit (Qiagen) according to manufacturer’s instructions. Synthesis of cDNA was performed using the iScript cDNA Synthesis Kit (BioRad) according to manufacturer’s instructions. 1-2ug of cDNA was diluted to 5ng/mL and was amplified by specific primers in a 20uL reaction using *Power* SYBR Green PCR Master Mix (Applied Biosystems). Analysis of gene expression was carried out in a LightCycler 480 II instrument (Roche). For each gene, mRNA expression was calculated as 2^deltaCT relative to *Actb* and *Hprt1* expression. Primer sequences used are provided in **Table S5**.

### Western Blotting

The VAT was collected and immediately snap frozen in liquid nitrogen. The VAT was pulverized using a mortar and pestle chilled with liquid nitrogen and homogenized with 300-500uL of RIPA buffer (Sigma) containing protease inhibitor (Sigma) and phosphatase inhibitors 2 and 3 (Sigma). Samples were left on ice for 30 min with vortexing every 5 min to disrupt membranes and then centrifuged at 13,200 rpm at 4°C for 15 min. Supernatant was collected and protein concentration was quantified using the DC protein assay (Bio-Rad). 30ug of protein lysate containing 1x NuPAGE Sample Reducing Agent (Invitrogen) and 1x NuPAGE LDS Sample Buffer (Life Technologies) were heated at 95 °C for 10 min and then separated by SDS-PAGE using a NuPAGE 4-12% Bis-Tris Gel (Invitrogen). Proteins were transferred onto nitrocellulose membranes using a semi-dry transfer system (BioRad). Membranes were then blocked in 5% BSA in 1x TBST (BioRad) and incubated in primary antibodies overnight. The blots were probed with the following antibodies: Phosphorylated HSL S536 (Cell Signaling; 4139S), Phosphorylated HSL S660 (Cell Signaling; 45804S), Total HSL (Cell Signaling; 4107S), ATGL (Cell Signaling; 2439S), MAOA (Abcam; ab126751), β-Actin (Cell Signaling; 4967S), HSP-90 (Cell Signaling; 4874S), and Ndufb8, Sdhb, Uqcrc2, Atp5a (MitoProfile total OXPHOS, Abcam; 110413). HRP-conjugated anti-mouse (Invitrogen) or anti-rabbit (Invitrogen) secondary antibodies were used appropriately. Images were collected using a ChemiDoc MP Imaging System (BioRad) and analyzed using Image Lab software (BioRad).

### Cytokine Measurements

Blood was collected and incubated at room temperature for 30 min then centrifuged at 3,000 rpm at 4°C for 20 min to isolate the serum. The concentrations of serum IL-1β, IL-6, GROα, TNFα, MCP-1, and Eotaxin were determined by 6-plex magnetic bead panel (Invitrogen) following the manufacturer’s instructions with appropriate dilution. Analysis of cytokine levels was carried out in a Luminex 200 instrument (Thermo Fisher Scientific).

### Free Fatty Acid Measurements

Blood was collected and incubated at room temperature for 30 min then centrifuged at 3,000 rpm at 4°C for 20 min to isolate the serum. Serum free fatty acid levels were measured using the Free Fatty Acid Assay Kit (Sigma MAK466) per manufacturer instructions.

### Whole Mount Immunofluorescence and Confocal Microscopy

Adipose tissues were fixed in 4% paraformaldehyde (PFA) overnight at 4°C, washed 3 times with PBS for 30 minutes, followed by permeabilization with 5% BSA and 0.2% Triton X-100 overnight at 4°C or permeabilization with 5% BSA and 0.5% Triton X-100 for 48-hours at 4°C. Tissues were stained with primary antibodies for 48 hours at 4°C. The following primary antibodies were used: TUBB3 (TUJ1, 1:200), CD169 (SER4, 1:100), MBP (P82H9, 1:100), and CNP (SMI 91, 1:100). Tissues were washed and mounted on a glass slide and coverslip in mounting media (DAKO S3023). For imaging of large nerve fibers, we isolated nerves from iWAT but kept surrounding adipocytes to preserve nerve integrity and processed as described above. Samples were imaged by confocal microscopy using Leica Stellaris 5 or 8 and analyzed using ImageJ or Leica Application Suite X.

### Transmission Electron Microscopy (TEM)

Mice were perfused with 4% PFA. iWAT was collected and maintained in 4% PFA. 1-2mm^2^ pieces of tissues containing nerve fibers were dissected out and fixed in 2.5% glutaraldehyde and 2% paraformaldehyde in 0.1M cacodylate buffer (pH 7.4) for 2 hours at room temperature and overnight at 4°C. They were then rinsed and post-fixed in 1% OsO4 at room temperature for 1 hour. After staining *en bloc* with 2% aqueous uranyl acetate for 30 min, the tissue was dehydrated in a graded series of ethanol to 100% propylene oxide and finally embedded in EMbed 812 resin. For TEM, sample blocks were then polymerized in 60°C oven for 24 hr. Thin sections (60 nm) were cut by a Leica ultramicrotome and post-stained with 2% uranyl acetate and lead citrate. Sample were examined with a FEI Tecnai transmission electron microscope at 80 kV of accelerating voltage, digital images were recorded with an Olympus Morada CCD camera and iTEM imaging software.

### Intravital Two-photon Microscopy

LysM^Cre^:Rosa26^mTmG^ (LysM:mTmG) or CD169^Cre^:Rosa26^mTmG^ (CD169:mTmG) mice were anesthetized with Ketamine/ Xylazine and placed in dorsal recumbency position on a heating pad. For VAT, a 1cm abdominal incision was made to open the skin and peritoneal cavity. The VAT was gently pulled out through the incision and placed on a metal support arm. For the iWAT, a 4-5 cm curved abdominal incision was made to create a lateral skin flap exposing the iWAT lying on the inside of the skin. Skin and iWAT were placed over a metal support arm resting next to the abdomen. A coverslip was placed over the tissue on the support arm using a micromanipulator. Sterile gauze and a warm saline infusion were utilized to ensure moistening of exposed tissue and abdominal cavity during imaging. The animals were restrained for approximately 4 hours and kept under isoflurane anesthetic by a precision vaporizer. After imaging was completed, the animals were euthanized. Inguinal adipose tissues were visualized by two-photon microscopy using LaVision Biotec TriMScope (Miltenyi Biotec, Germany) and analyzed using Imaris (Oxford Instruments) software.

### Bulk RNA-sequencing Analysis

mRNA from VAT and BAT was purified from approximately 1 ng of total RNA with oligo-dT beads and sheared by incubation at 94 °C. RNA integrity was determined by running an Agilent Bioanalyzer gel, which measures the ratio of the ribosomal peaks. Following first-strand synthesis with random primers, second-strand synthesis is performed with dUTP for generating strand-specific sequencing libraries. The cDNA library was then end-repaired, and A-tailed, adapters are ligated and second-strand digestion is performed by uricil-DNA-glycosylase. Indexed libraries that meet appropriate cut-offs for both are quantified by qRT–PCR using a commercially available kit (KAPA Biosystems). Insert size distribution was determined with the LabChip GX or Agilent Bioanalyzer. Samples with a yield of ≥0.5 ng μl−1 were used for sequencing. Raw reads were quality-assessed with FastQC ^41^. They were mapped to the GENCODE vM9 mouse reference genome ^42^ with STAR ^43^ using options: outFilterMultimapNmax 15 -- outFilterMismatchNmax 6 --outSAMstrandField All --outSAMtype BAM SortedByCoordinate --quantMode TranscriptomeSAM. Gene expression was quantified with RSEM ^44^. PCA was performed in R after removing a donor effect with the ComBat function from the sva R-package (version 3.28.0)^45^. PCA was based on the rlog-transformed FPKM values of the top 5,000 genes with the highest average expression.

### Single Cell RNA-sequencing Analysis

Intra-vascular labeling was performed as previously described ^15^. Briefly, 2- and 22-month-old mice received 2.5ug of murine CD45-APC antibody i.v. by retro-orbital injection to label circulating cells (5 mice per group). Mice were euthanized 3 minutes post antibody administration, and the visceral adipose tissue (VAT) was digested as previously described (refer to ‘Adipose Tissue Digestion and Flow Cytometry’ section). Following adipose tissue digestion, stromal vascular fraction (SVF) cells isolated from VAT were then stained with viability dye, CD45.2, CD45, CD3, CD19, SiglecF, F4/80, CD11b, and were were sorted using a BD FACSAria II and analyzed using Chromium Next GEM Automated Single Cell 3’ cDNA Kit v3.1 (10X genomics). Experiments on 2- and 22-month-old female mice were performed the next day using the same protocol (5 mice per group). The Cell Ranger Single-Cell Software 3.0.2 available at 10x website was used to perform sample demultiplexing and resulting fastq files were aligned on mm10 genome reference (10x website, https://cf.10xgenomics.com/supp/cell-exp/refdata-gex-mm10-2020-A.tar.gz) with cellranger count tool. Data analysis was performed using Seurat v4.3.0 R package, including quality control, dimension reduction, clustering, cell-type identification and comparative analyses between conditions ^46–50^. In the step of quality control, poor-quality cells with the number of expressed genes < 363 (363 corresponding to the first mode in the bimodal distribution of number of expressed genes) were filtered out. We also excluded cells if their mitochondrial gene percentages were over 10%. After combining cells from all samples, we first normalized the raw count matrix using Log-Normalization with scale.factor 10,000 and then defined top variable genes using “mean.var.plot” methods. We then applied principal component analysis (PCA) for dimensionality reduction and retained 20 leading principal components for cell clustering. The Shared Nearest Neighbor (SNN) graph was constructed using default hyperparameters, which was then used for cell clustering with the Louvain algorithm at a resolution of 0.5. Cluster-specific genes were found by Wilcoxon Rank Sum test (min.pct = 0.1, logfc.threshold = 0.1). After identifying cluster-specific genes, we annotated cell types based on canonical marker genes. Clusters 9, 12, and 14 did not express significant levels of *Ptprc* (CD45), *Itgam* (CD11b), and *Adgre1* (F4/80), suggesting these clusters were likely contaminating cells and were therefore excluded from future analyses. Differentially expressed genes for condition were identified by performing Wilcoxon Rank Sum test on each cluster for condition Old and Young (min.cell = 25, min.pct = 0.01, logfc.threshold = log(2)).

### Enrichment Analysis

Enrichment analysis was performed using ClusterProfiler V4.11.0 R package on DEGs found in scRNAseq between clusters and conditions ^51^. R package msigdbr V7.5.1 was used to retrieve terms of interest from The Molecular Signatures Database ^52^. Specifically, Gene Ontology (GO) terms were retrieved from species *Mus musculus* and category C5 (subcategory GO:BP, GO:CC, GO:MF), and Hallmark Gene Sets were retrieved from species *Mus musculus* and category H. Benjamini-Hochberg method was used for adjusting p values. The threshold cutoff for p value and q value were set to be 0.05. Single cell gene set enrichment score (GSES) was calculated for each cell using the “enrichIt” function from R package “escape” V1.10.0, with method set to UCell ^53,54^. Single cell GSES was calculated for the following gene sets: SenMayo (SAUL_SEN_MAYO, GSEA: MM16098) and NAMs (LEVEAU_NERVE_ASSOCIATED_MACROPHAGES, **Table S3**).

### Trajectory Analysis

The Velocyto tool was employed to generate two count matrices for each sample: one for pre-mature (unspliced) and another for mature (spliced) RNA abundances ^55^. These matrices serve as the input for subsequent RNA velocity inference. Specifically, the data from four samples were combined and processed using scVelo (V0.2.5) ^56^. Preprocessing steps include gene filtering (removing genes with a minimum shared count of less than 20), normalization, extracting top 5000 highly variable genes, and Log1p transform. RNA velocities were inferred based on the steady-state model. Velocities were projected onto the UMAP cell embedding for visualization, which facilitates inference of cell state transition dynamics.

## Acknowledgements

We thank Xinran Liu MD, PhD from the Yale Center for Cellular and Molecular Imaging (CCMI) Electron Microscopy facility for TEM and FIB-SEM imaging. We thank Diane Trotta from the Yale Flow Cytometry Facility for cell sorting on BD FACSAria II.

## Author Contributions

EGH and CL performed experiments, analyzed data, and wrote the manuscript. KL and RQ analyzed the scRNAseq data. MM performed the adipose tissue digestion and flow cytometry for quantification of nerve associated macrophages in young and aged females in VAT. EG performed and analyzed the bulk RNAseq of resident versus circulating macrophages in VAT and BAT. SS analyzed bulk RNAseq data of resident versus circulating macrophages in VAT and BAT. STY and CK administered diphtheria toxin, monitored body weight, and collected tissue samples from CD169-DTR mice. TMS performed and analyzed RT-qPCR of VAT from 9-month CD169-DTR mice. DG performed the two-photon *in vivo* imaging in CD169^Cre^:Rosa^mTmG^ mice (CD169:mTmG). CC analyzed the two-photon videos in LysM^Cre^:Rosa26^mTmG^ mice. MNA provided critical advising for the bioinformatic analysis. KMK provided critical reviews and supervised the depletion experiments with CD169-DTR mice. YK provided critical reviews and supervised the bioinformatic analysis and interpretation of the scRNAseq data. VDD conceived the project, helped with data analysis, interpretation, and preparation of the manuscript.

## Data Material and Availability

Sequencing data will be deposited on Gene Expression Omnibus (GEO) and are available upon request.

## Supplemental Figures

**Supplemental Figure 1.**
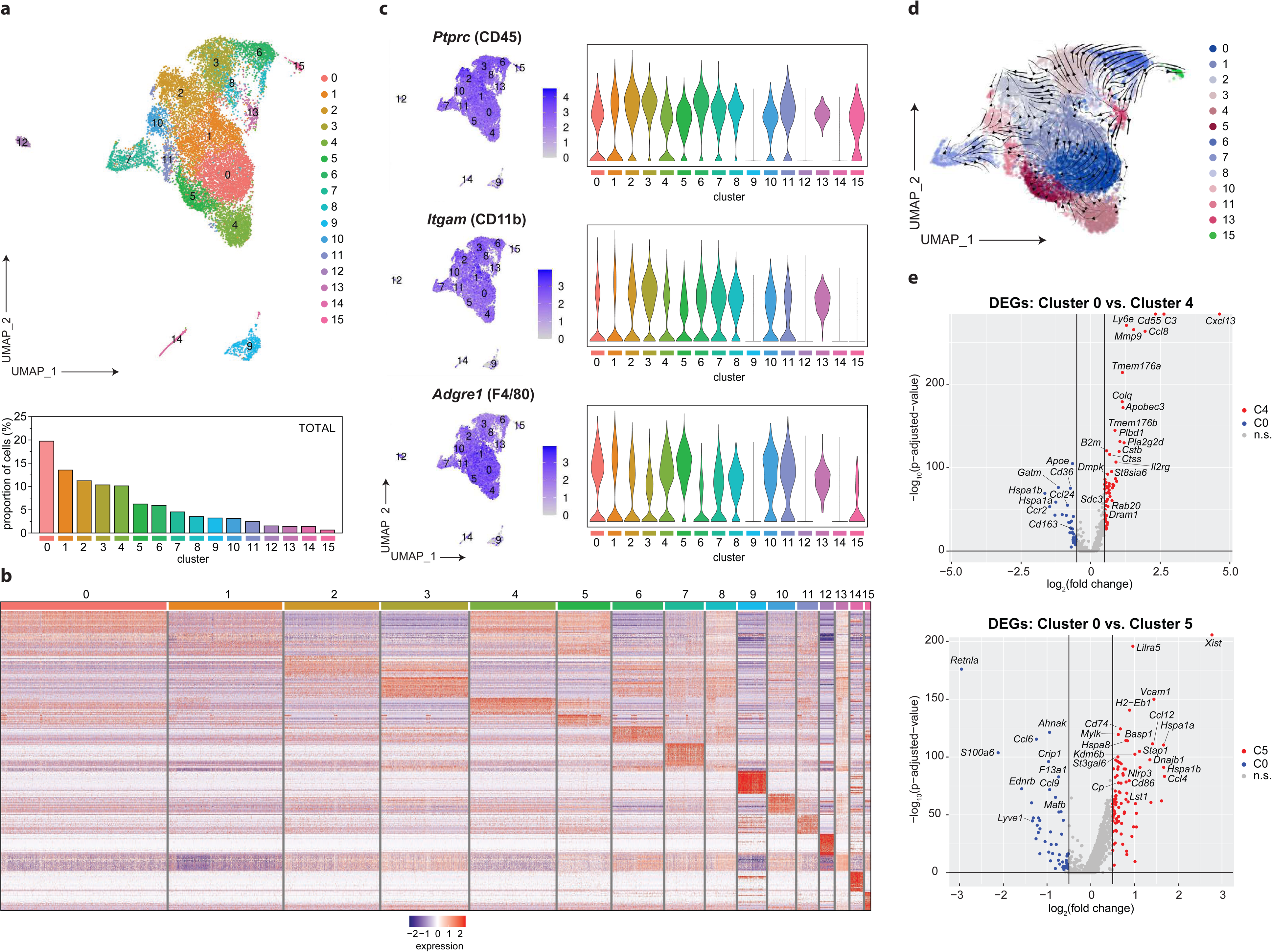
(A) Uniform Manifold Approximation and Projection (UMAP) representation of unsupervised clustering of 15,709 resident F4/80^+^CD11b^+^ (CD45^+^CD45.2iv^−^CD3^−^CD19^−^) cells sorted from VAT of young and aged males (13,134 cells) and females (2,575 cells) (*top*) and repartition of cells in each cluster as a fraction of total cells by cluster (*bottom*). (B) Heatmap depicting the top 25 differentially expressed genes (DEGs) per cluster. (C) Uniform Manifold Approximation and Projection (UMAP) (left) and violin plots (right) showing the expression of *Ptprc* (CD45), *Itgam* (CD11b), and *Adgre1* (F4/80) genes by cluster. (D) RNA velocity projected onto the Uniform Manifold Approximation and Projection (UMAP) plot. Velocities are based on the steady-state model. (E) Volcano plots depicting the top differentially expressed genes (DEGs) between Cluster 0 versus Cluster 4 (*top*) and Cluster 0 versus Cluster 5 (*bottom*). Genes with an adjusted p-value > 0.05 and log_2_ fold-change > 0.5 were deemed significant.

**Supplemental Figure 2.**
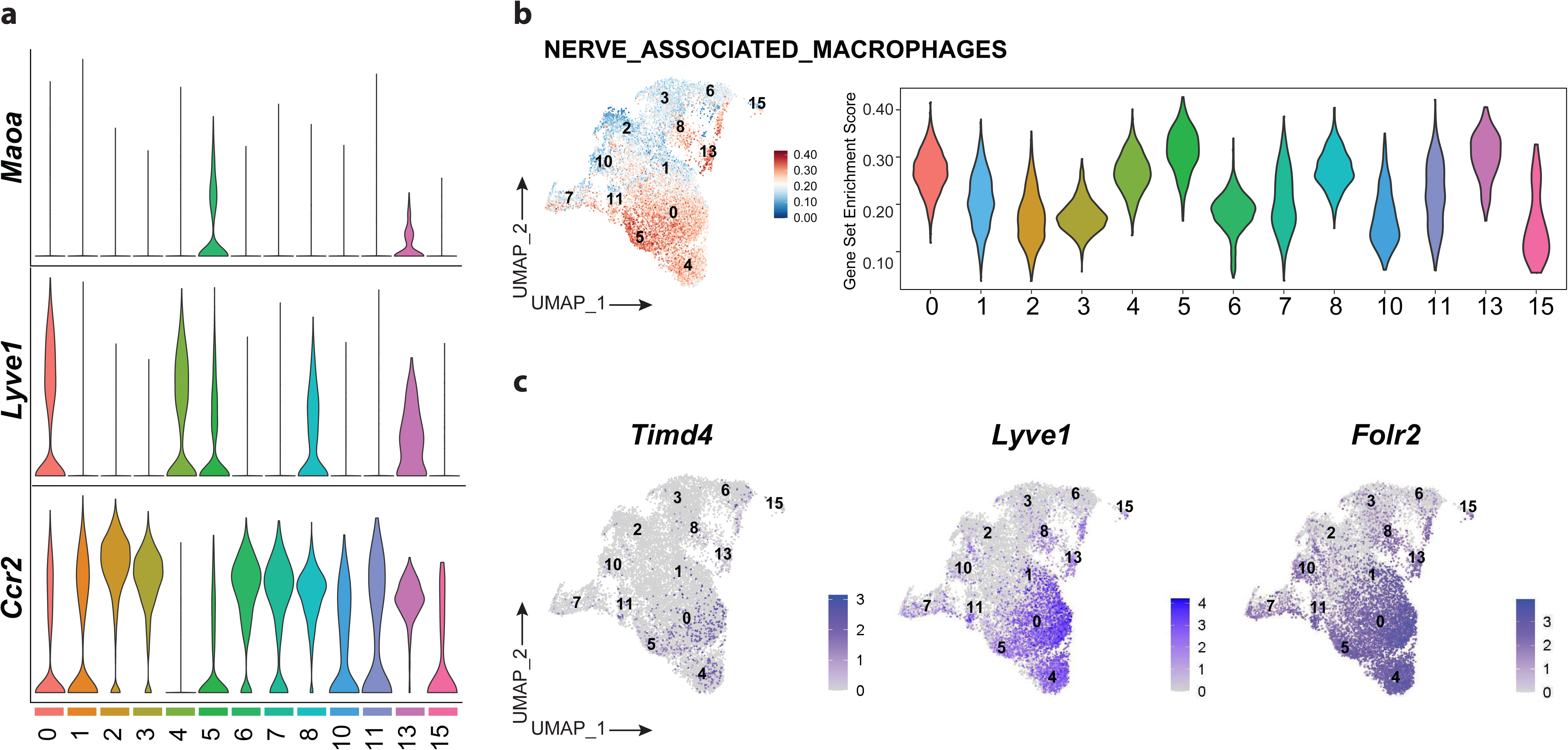
(A) Violin plot depicting the expression of *Maoa, Lyve1, and Ccr2* per cluster. (B) Uniform Manifold Approximation and Projection (UMAP) (*left*) and violin plot (*right*) showing enrichment of nerve-associated macrophage (NAM) genes and gene set enrichment score (GSES) per cell. The NAM gene set (LEVEAU_NERVE_ASSOCIAED_MACROPHAGES, **Table S3**) was constructed from bulk RNA-sequencing and microarray datasets of NAM-enriched populations in different tissues (adipose tissue, lung, gut, skin, and peripheral nerves); a gene was considered NAM-enriched and included in the NAM gene set if it appeared in > 5 of the published datasets assessed. (C) Uniform Manifold Approximation and Projection (UMAP) (*right*) depicting the expression of *Timd4*, *Lyve1*, and *Folr2* genes.

**Supplemental Figure 3.**
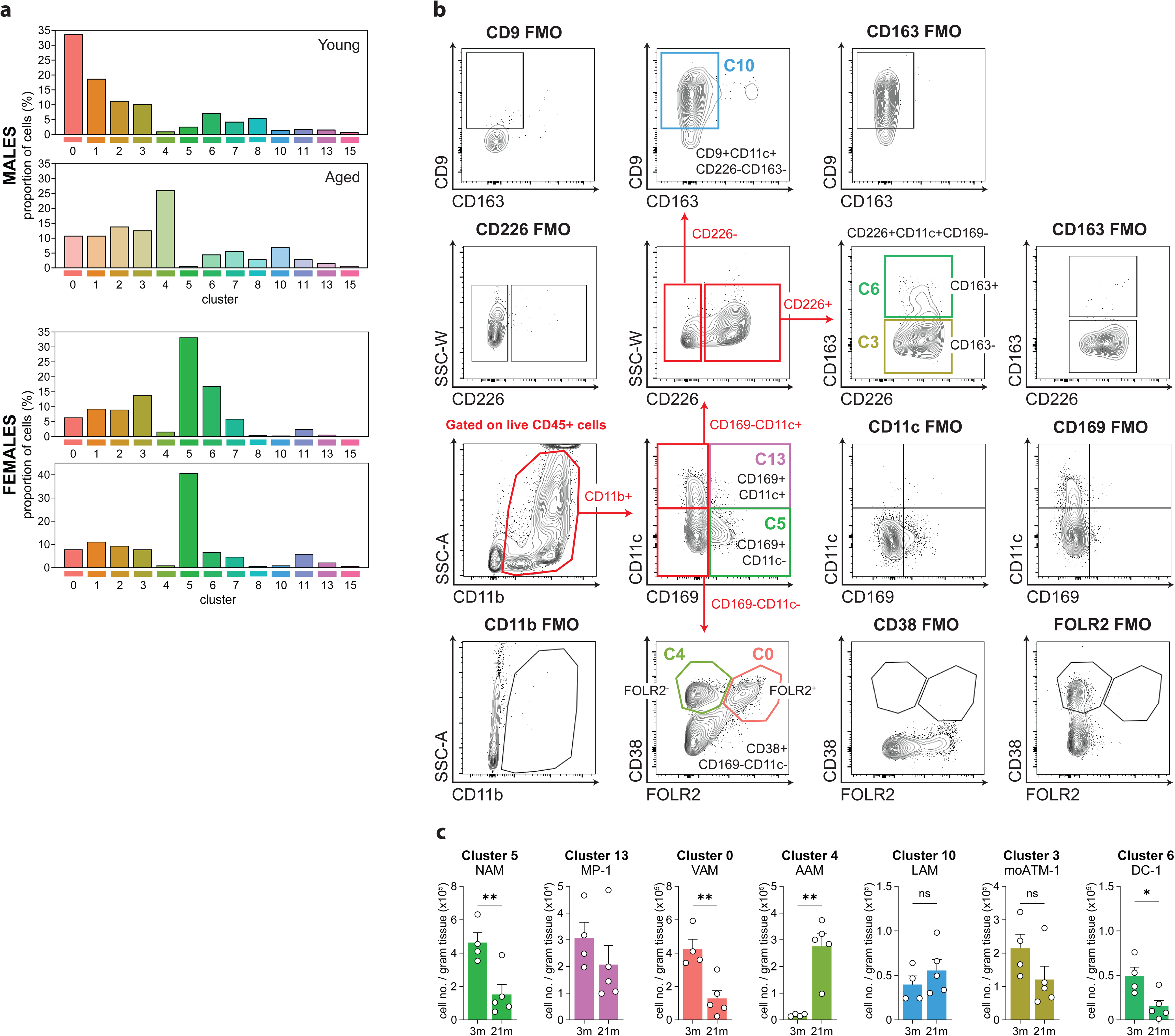
(A) Repartition of cells in each cluster as a fraction of total cells from young or aged VAT from males (*top*) and females (*bottom*). (B) Schematic depicting the flow cytometry gating strategy used to define clusters 0, 3, 4, 5, 6, 10, and 13 identified via single cell RNA-sequencing of resident F4/80^+^CD11b^+^ (CD45^+^CD45.2iv^−^CD3^−^CD19^−^) cells from the visceral white adipose tissue (VAT). Fluorescence minus one (FMO) control(s) were included for the following cell surface markers: CD11b, CD169, CD11c, CD226, CD9, CD163, FOLR2, and CD38. Abbreviations for each population and defining cell surface markers are as follows: Nerve-associated-macrophages (NAM, C5): CD169^+^CD11c^−^, Myeloid Precursors-1 (MP-1, C13): CD169^+^CD11c^+^, Vessel-associated Macrophages (VAM, C0): CD38^+^FOLR2^+^CD11c^−^ CD38^+^, Age-associated Macrophages (AAM, C4): CD38^+^FOLR2^−^CD169^−^CD11c^−^, Lipid-associated Macrophages (LAM, C10): CD9^+^CD11c^+^CD226^−^CD163^−^CD169^−^, Adipose Tissue Macrophages-1 (ATM-1, C3): CD226^+^CD11c^+^CD163^−^CD169^−^, Dendritic Cells-1 (DC-1, C6): CD163^+^CD226^+^CD11c^+^CD163^+^CD169^−^. (C) Quantification of cells from clusters 5, 13, 0, 4, 10, 3, and 6 as a fraction of live CD45+ cells per gram of visceral white adipose tissue (VAT). Data is displayed as the mean +/− SEM and is representative of one independent experiment from 3-month (n=4) and 21-month-old (n=5) females. Statistical significance was determined via Student’s two-tailed t-test with α=0.05 and *, p < 0.05; **, p < 0.01, n.s. = not significant.

**Supplemental Figure 4.**
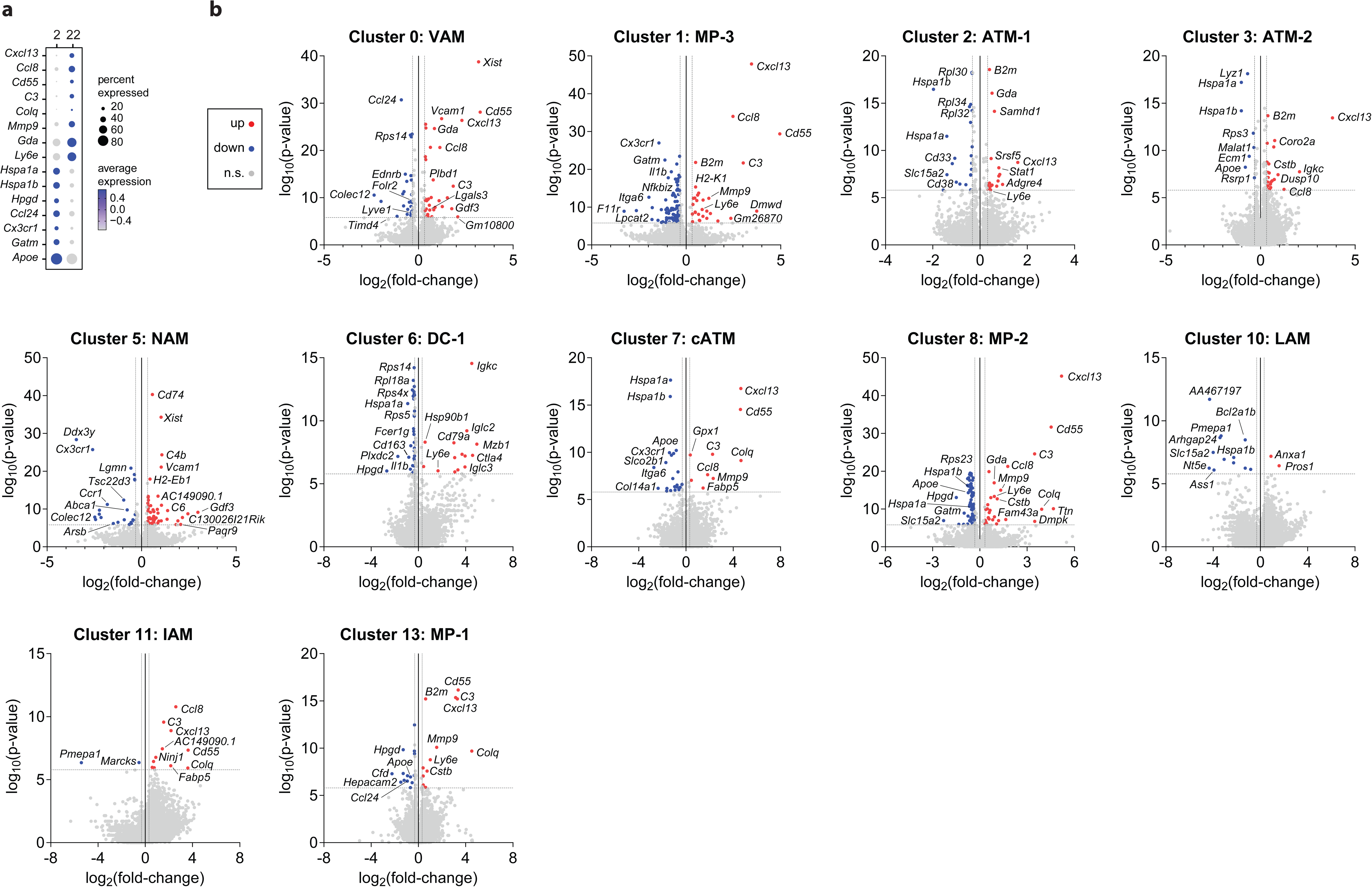
(A) Dot plot depicting the average gene expression of genes upregulated (*blue*) or downregulated (*gray*) by age in four or more clusters. Dot size represents the total percentage of cells expressing the corresponding gene. (B) Volcano plots depicting upregulated (red) and downregulated (blue) genes with age (aged versus young) for each cluster. The genes regulated by age were considered significant if the adjusted p-value > 0.05 and fold-change > 1.25.

**Supplemental Figure 5.**
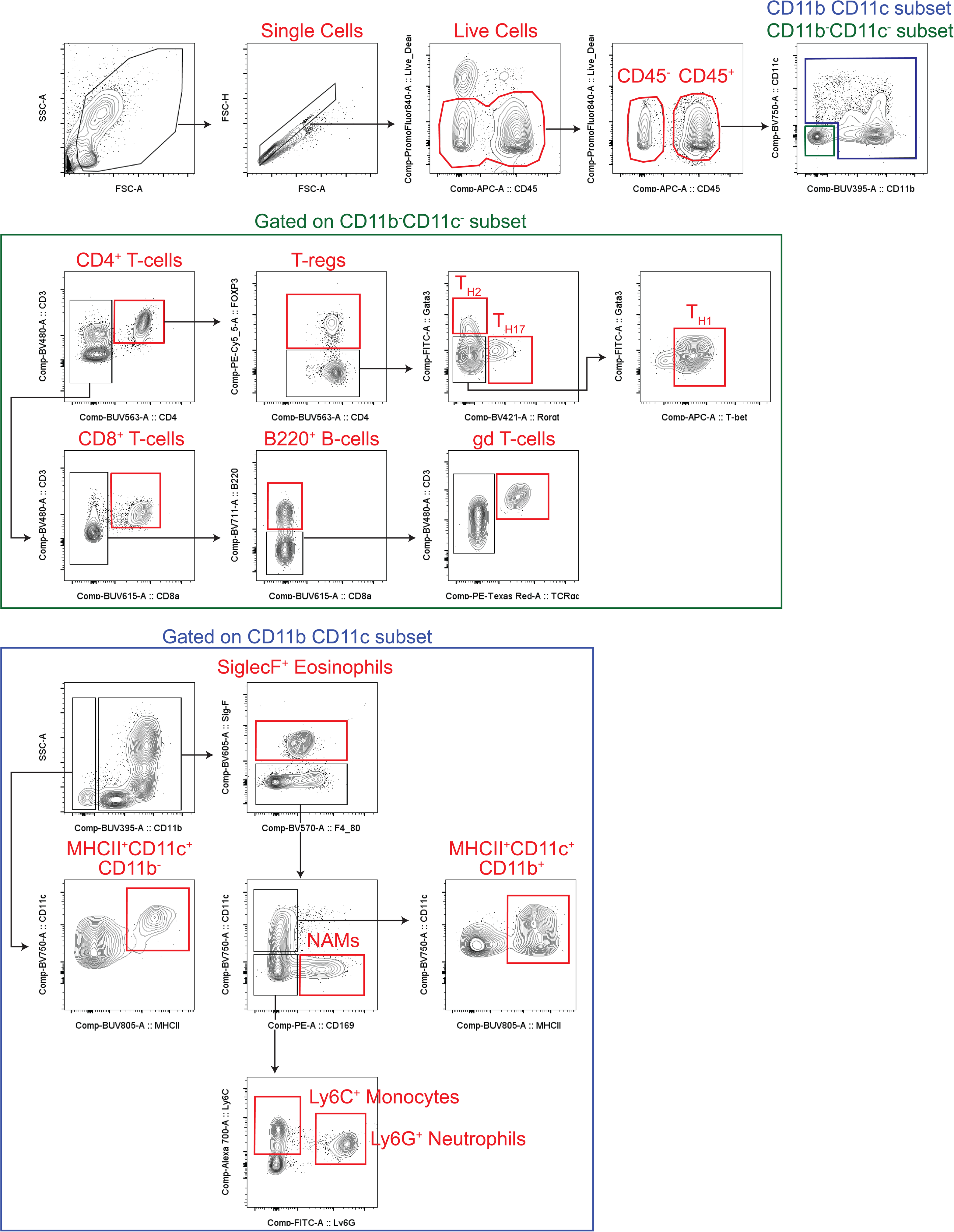
(A) Representative schematic depicting the flow cytometry gating strategy used to quantify the depletion efficiency of nerve-associated macrophages (NAMs) and other immune cell subsets in the visceral white adipose tissue (VAT) and spleen of wildtype (WT) and CD169-DTR mice following intraperitoneal diphtheria toxin administration in 3-, 6-, 9-, and 24-month-old females and at the following endpoints: day 8, day 13, day 25, and day 34.

**Supplemental Figure 6.**
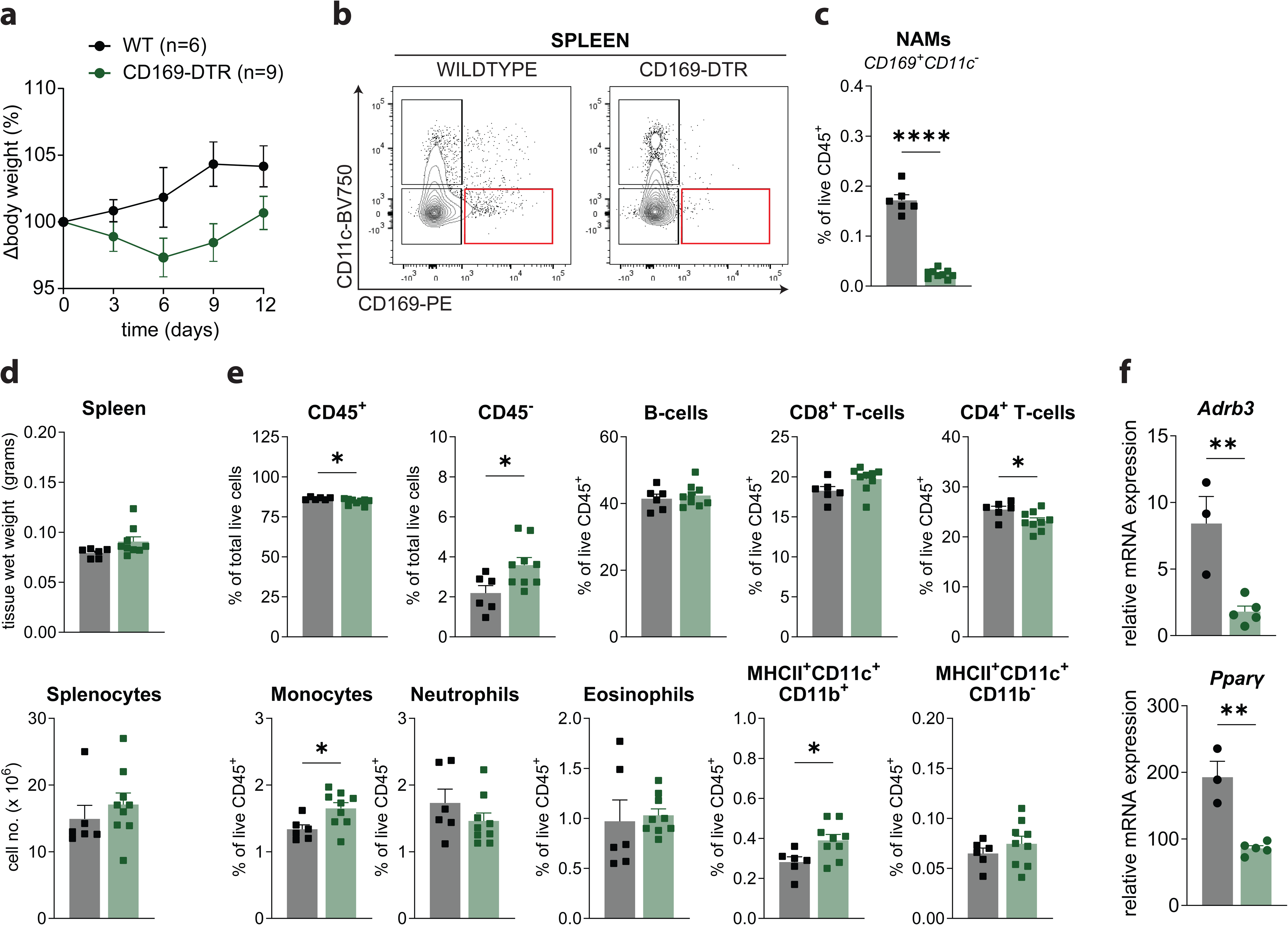
(A) Change in body weight curves for 3-month-old WT and CD169-DTR females (n=6-9 per genotype) treated with DT and sacrificed on D13. (B) Representative flow cytometry plots quantifying nerve-associated macrophages (NAMs) in the spleen from 3-month-old WT and CD169-DTR females treated with DT and sacrificed on D13 (n=6-9 per genotype). (C) Bar plot quantifying the proportion of nerve-associated macrophages in the spleen from 3-month-old WT and CD169-DTR females treated with DT and sacrificed on D13. Data is represented as a mean +/− SEM. Statistical significance was determined via Student’s two-tailed t-test with α=0.05 and p < 0.05. (D) Spleen tissue wet weight (*top*) and splenocyte cell number (*bottom*) of from 3-month-old WT and CD169-DTR females treated with DT and sacrificed on D13. Data is represented as a mean +/− SEM. Statistical significance was determined via Student’s two-tailed t-test with α=0.05 and p < 0.05. (E) Flow cytometry of the spleen from 3-month-old WT and CD169-DTR females treated with DT and sacrificed on D13. Bar plots represent either the proportion of CD45^+^ and CD45^−^ cells (as a fraction of total live cells) and B-cells, CD8^+^ T-cells, CD4^+^ T-cells, Monocytes, Neutrophils, Eosinophils, and MHCII^+^CD11c^+^ ATMs or dendritic cells (DCs) (as a fraction of live CD45^+^). Populations were defined using the cell surface markers and gating strategy outlined in Supplementary Figure 5. Data is represented as a mean +/− SEM. Statistical significance was determined via Student’s two-tailed t-test with α=0.05 and p < 0.05. (F) Relative mRNA expression of *Adrb3* (*top*) and *Pparγ* (*bottom*) in the VAT of *ad libitum* fed 9-month-old WT and CD169-DTR females treated with DT and sacrificed on D8 (n=3-5 per genotype). Data is represented as a mean +/− SEM. Statistical significance was determined via via Student’s two-tailed t-test with α=0.05 and p < 0.05.

**Supplemental Figure 7.**
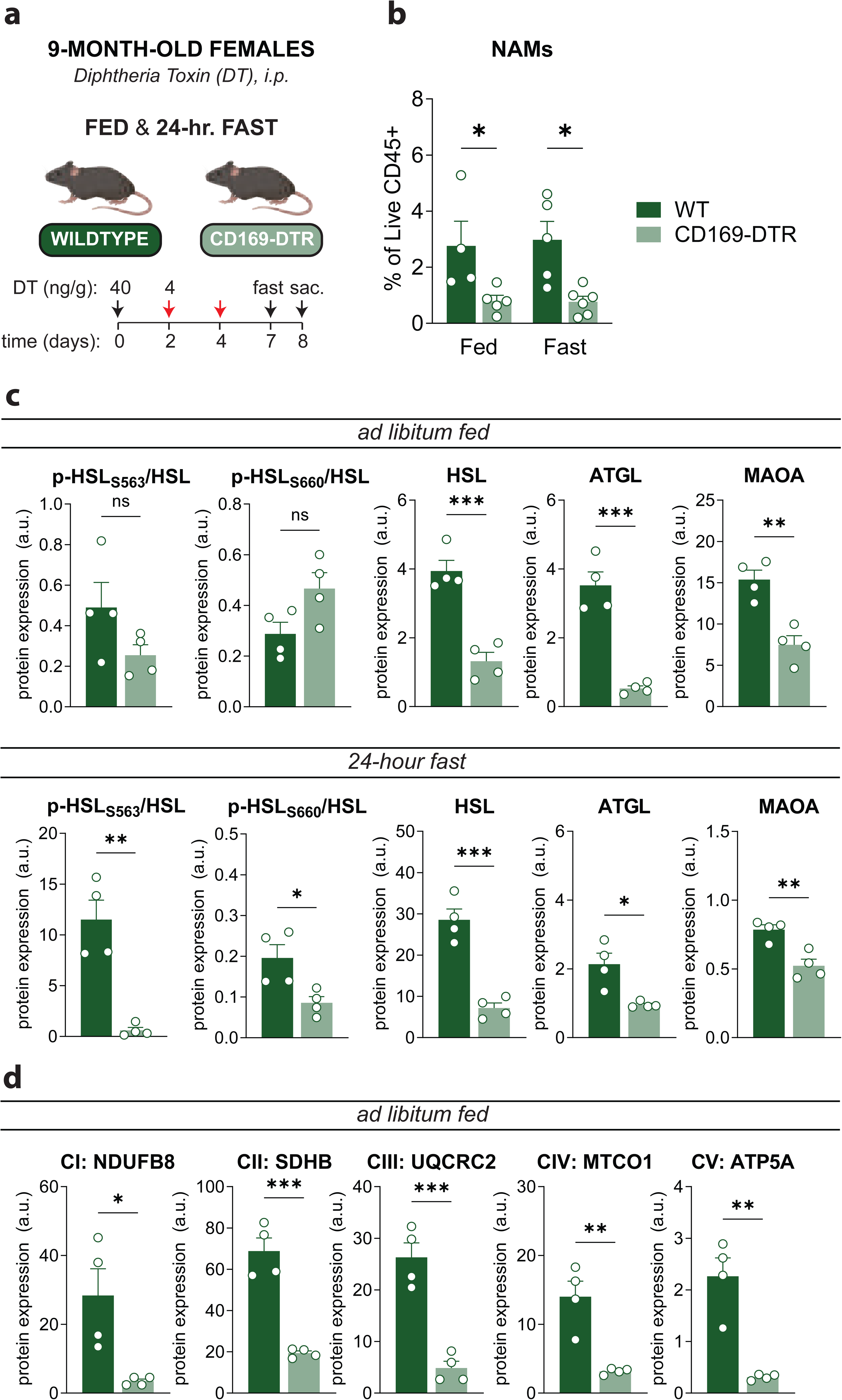
(A) Schematic representation of the experimental protocol used to deplete CD169-macrophages using CD169 diphtheria toxin receptor (CD169-DTR) mice. 9-month-old Wildtype (WT) and CD169-DTR (n=9-11) female mice were injected intra-peritoneal (i.p) with 40ng/gram diphtheria toxin (DT) on day 0 (D0) followed by 4ng/gram DT on D2 and D4. On day 7 (D7) mice either remained *ad libitum* fed (n=4-5 per genotype) or were fasted for 24-hours (n=5-6 per genotype) and then scarified on D8. (B) Quantification of nerve-associated macrophages (NAMs) in the stromal vascular fraction (SVF) of VAT from 9-month-old Wildtype (WT) and CD169-DTR females treated with DT that were either *ad libitum* fed or 24-hour *fasted* and then sacrificed on D8. Data is represented as a mean +/− SEM. Statistical significance was determined via Student’s two-tailed t-test with α=0.05 and p < 0.05. (C) Relative protein quantification of the following proteins: Hormone-sensitive Lipase (HSL), HSL phosphorylated at Ser563 (p-HSL_S563_), HSL phosphorylated at Ser660 (p-HSL_S660_), Adipose triglyceride lipase (ATGL), and Monoamine-oxidase A (MAOA). Protein lysate is from the VAT of *ad libitum* fed (*top panel*) or 24-hour fasted (*bottom panel*) 9-month-old WT and CD169-DTR females treated with DT and sacrificed on D8 (n=4 per genotype). Data is represented as a mean +/− SEM. Statistical significance was determined via Student’s two-tailed t-test with α=0.05 and p < 0.05. (D) Relative protein quantification of oxidative phosphorylation proteins: Complex I (CI): NADH:ubiquinone oxidoreductase subunit B8 (NDUFB8), Complex II (CII): succinate dehydrogenase complex, subunit B, iron sulfur (Ip) (SDHB), CIII: ubiquinol cytochrome c reductase core protein 2 (UQCRC2), Complex IV (CIV): mitochondrially encoded cytochrome c oxidase I (MTCO1), Complex V (CV): ATP synthase F1 subunit alpha (ATP5A). Protein lysate is from the VAT of *ad libitum* fed 9-month-old WT and CD169-DTR females treated with DT and sacrificed on D8 (n=4 per genotype). Data is represented as a mean +/− SEM. Statistical significance was determined via Student’s two-tailed t-test with α=0.05 and p < 0.05.

**Supplemental Figure 8.**
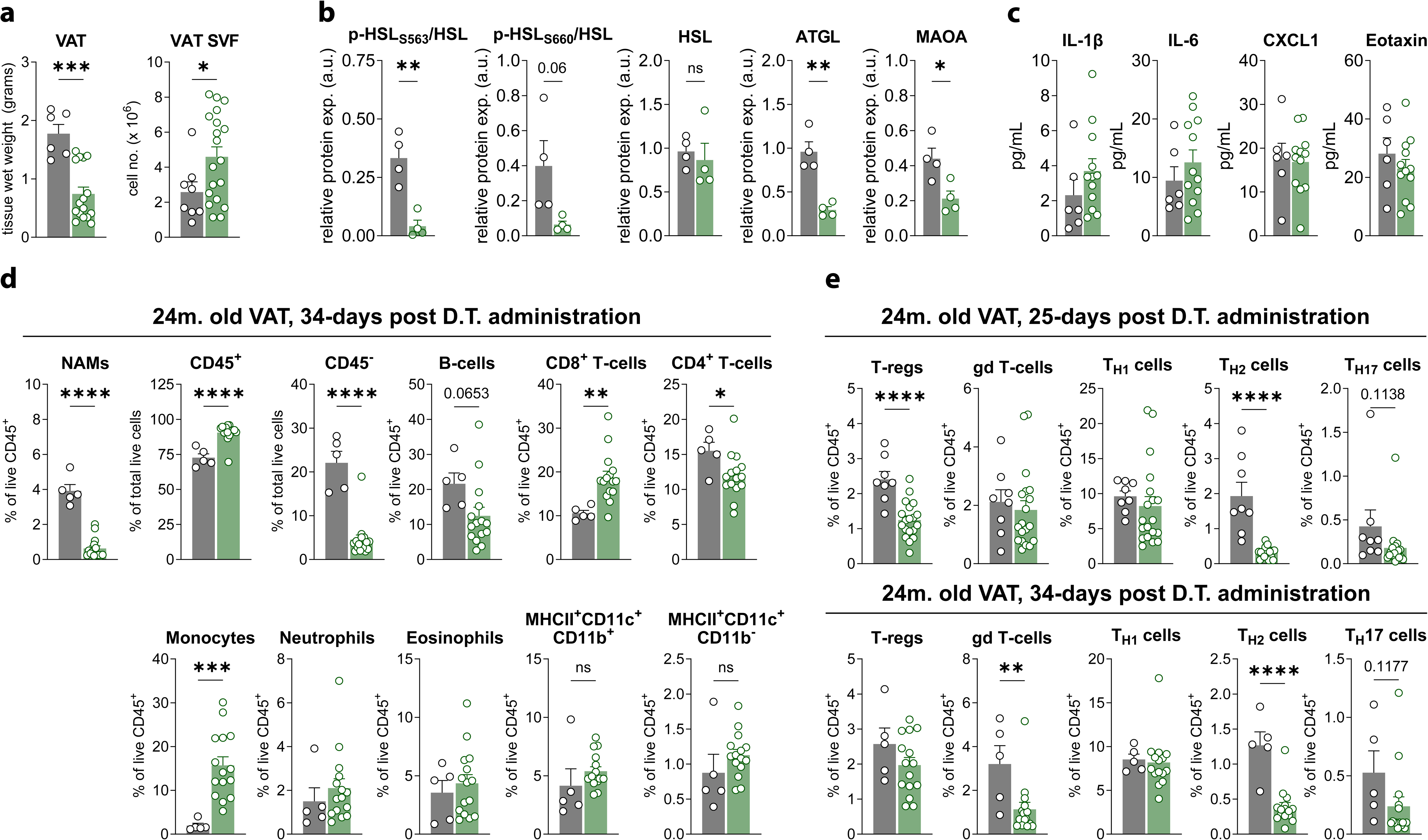
(A) Visceral white adipose tissue (VAT) wet weight (*right*) and stromal vascular fraction (SVF) cell number (*left*) from 24-month-old Wildtype (WT) and CD169-DTR females treated with DT and sacrificed on D25 (n=6-13 per genotype). Data is represented as a mean +/− SEM. Statistical significance was determined via Student’s two-tailed t-test with α=0.05 and p < 0.05. (B) Relative protein quantification of the following proteins: Hormone-sensitive Lipase (HSL), HSL phosphorylated at Ser563 (p-HSL_S563_), HSL phosphorylated at Ser660 (p-HSL_S660_), Adipose triglyceride lipase (ATGL), and Monoamine-oxidase A (MAOA). Protein lysate is from the VAT of *ad libitum* fed 24-month-old WT and CD169-DTR females treated with DT and sacrificed on D25 (n=4 per genotype). Data is represented as a mean +/− SEM. Statistical significance was determined via Student’s two-tailed t-test with α=0.05 and p < 0.05. (C) IL-1β, IL-6, CXCL1, and Eotaxin levels in the serum of *ad libitum* fed 24-month-old Wildtype (WT) and CD169-DTR females treated with diphtheria toxin (DT) and sacrificed on D25 (n=5-11 per genotype). Data is represented as a mean +/− SEM. Statistical significance was determined via Student’s two-tailed t-test with α=0.05 and p < 0.05. (D) Flow cytometry from the VAT of 24-month-old WT and CD169-DTR females treated with DT and sacrificed on D34. Bar plots represent either the proportion of CD45^+^ and CD45^−^ cells (as a fraction of total live cells) and B-cells, CD8^+^ T-cells, CD4^+^ T-cells, Monocytes, Neutrophils, Eosinophils, and MHCII^+^CD11c^+^ ATMs or dendritic cells (DCs) (as a fraction of live CD45^+^). Populations were defined using the cell surface markers and gating strategy outlined in Supplementary Figure 5. Data is represented as a mean +/− SEM. Statistical significance was determined via Student’s two-tailed t-test with α=0.05 and p < 0.05. (E) Flow cytometry from the VAT of 24-month-old WT and CD169-DTR females treated with DT and sacrificed on D25 (*top*) or D34 (*bottom*). Bar plots represent either the proportion of regulatory T-cells, gd T-cells, TH1 cells, TH2 cells, and TH17 cells. Populations were defined using the cell markers and gating strategy outlined in Supplementary Figure 5. Data is represented as a mean +/− SEM. Statistical significance was determined via Student’s two-tailed t-test with α=0.05 and p < 0.05.

**Supplemental Figure 9.**
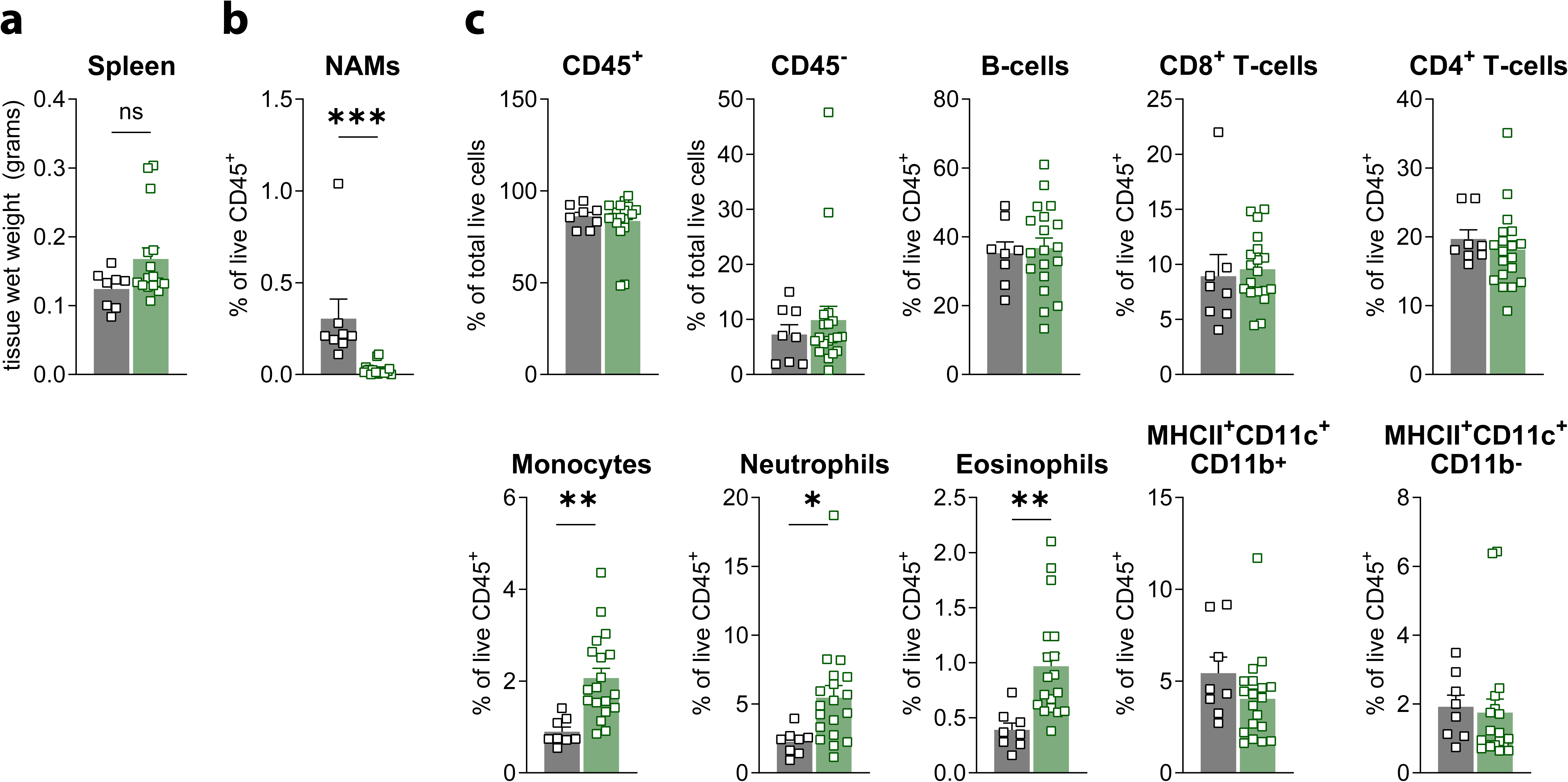
(A) Tissue wet weight of the spleen from 24-month-old Wildtype (WT) and CD169-DTR females treated with DT and sacrificed on D25 (n=8-19 per genotype). Data is represented as a mean +/− SEM. Statistical significance was determined via Student’s two-tailed t-test with α=0.05 and p < 0.05. (B) Bar plot quantifying nerve-associated macrophages in the spleen from 24-month-old WT and CD169-DTR females treated with DT and sacrificed on D13. Data is represented as a mean +/− SEM. Statistical significance was determined via Student’s two-tailed t-test with α=0.05 and p < 0.05. (C) Flow cytometry from the spleen of 24-month-old WT and CD169-DTR females treated with DT and sacrificed on D25. Bar plots represent either the proportion of CD45^+^ and CD45^−^ cells (as a fraction of total live cells) and B-cells, CD8^+^ T-cells, CD4^+^ T-cells, Monocytes, Neutrophils, Eosinophils, and MHCII^+^CD11c^+^ ATMs or dendritic cells (DCs) (as a fraction of live CD45^+^). Populations were defined using the cell surface markers and gating strategy outlined in Supplementary Figure 5. Data is represented as a mean +/− SEM. Statistical significance was determined via Student’s two-tailed t-test with α=0.05 and p < 0.05.

## Supplemental Tables

**Table S1**. Quality control (QC) metrics for single cell RNA-sequencing samples from resident F4/80^+^CD11b^+^ cells sorted from the visceral white adipose tissue (VAT) of 2- and 22-mont-old male and female mice. Resident cells were defined as: CD45.1^+^CD45.2iv^−^CD3^−^CD19^−^.

**Table S2**. List of differentially expressed genes (DEGs) for each cluster.

**Table S3**. Nerve-associated macrophage (NAM) gene set list.

**Table S4**. List of differentially expressed genes (DEGs) in the aged compared to young condition for each cluster.

**Table S5**.List of primer sequences used for real time quantitative PCR (RT-qPCR).

## Supplemental Videos

**Video S1**. *In vivo* two-photon imaging of visceral white adipose tissue (VAT) from a 24-month old male LysM:mTmG reporter mouse which labels LysM+ cells with membrane EGFP (mGFP+) (*green*) and non-LysM (non-recombined) cells/tissues such as adipocytes, nerves, and blood vessels with membrane tdTomato (mTomato+) (*red*). Scale bar: 100µm. Data is representative of n = 1 independent experiment.

**Video S2**. Serial sections from whole mount immunofluorescence imaging of VAT from CD169:mTmG reporter mice which labels CD169^+^ macrophages with membrane EGFP (mGFP) (*green*) and non-CD169 (non-recombined) cells/tissues such as adipocytes, nerves, and blood vessels with membrane tdTomato (mTomato+) (*red*). VAT was stained with the pan-neuronal marker, TUBB3. Scale bar: 20µm.

**Video S3**. Serial sections from whole mount immunofluorescence imaging of VAT from CD169:mTmG reporter mice which labels CD169^+^ macrophages with membrane EGFP (mGFP) and non-CD169 (non-recombined) cells/tissues such as adipocytes, nerves, and blood vessels with membrane tdTomato (mTomato). VAT was stained with DAPI and the pan-neuronal marker, TUBB3. Scale bar: 1µm.

**Video S4**. 3D surface rendering reconstruction of a CD169^+^ nerve-associated macrophage (NAM) (*red*) associated with an MBP^+^ myelinated nerve fiber (*green*) in the VAT from wildtype mice. Tissue was stained with antibodies against CD169 (red), MBP (green), and TUBB3 (blue). Data is representative of n = 1 independent experiment using 3-month-old wildtype males (n = 2). Scale bar: 10µm.

**Video S5**. 3D surface rendering reconstruction of a CD169^+^ nerve-associated macrophage (NAM) (*red*) associated with an MBP^+^ myelinated nerve fiber (*green*) in VAT from wildtype mice. Tissue was stained with antibodies against CD169 (red), MBP (green), and TUBB3 (blue). Data is representative of n = 1 independent experiment using 3-month-old wildtype males (n = 2). Scale bar: 10µm.

**Video S6**. *In vivo* two-photon imaging of inguinal white adipose tissue (iWAT) from adult CD169:mTmG reporter mice which labels CD169^+^ macrophages with membrane EGFP (mGFP) (*green*) and non-CD169 (non-recombined) cells/tissues such as adipocytes, nerves, and blood vessels with membrane tdTomato (mTomato+) (*red*). Data is representative of n = 2 independent experiments using 3-month-old males.

**Video S7**. *In vivo* two-photon imaging of iWAT from adult CD169:mTmG reporter mice which labels CD169^+^ macrophages with membrane EGFP (mGFP+) (*green*) and non-CD169 (non-recombined) cells/tissues such as adipocytes, nerves, and blood vessels with membrane tdTomato (mTomato+) (*red*). Data is representative of n = 2 independent experiments using 3-month-old males.

