## Supplementary material for "Nerve-associated macrophages control adipose homeostasis across lifespan and restrain age-related inflammation": Table S1

| Sample Condition | Cells Before Filtering |
| --- | --- |
| Old Females | 2607 |
| Old Males | 7082 |
| Young Females | 1233 |
| Young Males | 7878 |

| Filtering Criterial | Cells After Filtering |
| --- | --- |
| percent.mito <= 0.10 & nFeature_RNA >= 363 | 1749 |
| percent.mito <= 0.10 & nFeature_RNA >= 363 | 6079 |
| percent.mito <= 0.10 & nFeature_RNA >= 363 | 826 |
| percent.mito <= 0.10 & nFeature_RNA >= 363 | 7055 |
