## Supplementary material for "Nerve-associated macrophages control adipose homeostasis across lifespan and restrain age-related inflammation": Table S2

| p_val | avg_log2FC | pct.1 | pct.2 | p_val_adj | cluster | gene | pct.1-pct.2 |
|---|---|---|---|---|---|---|---|
| 0 | 2.3528095 | 0.758 | 0.086 | 0 | 5 | Xist | 0.672 |
| 0 | 1.1144675 | 0.495 | 0.045 | 0 | 5 | Lilra5 | 0.45 |
| 9.24E-302 | 0.9213565 | 0.598 | 0.135 | 2.87E-297 | 5 | St3gal6 | 0.463 |
| 2.46E-289 | 1.7475139 | 0.733 | 0.236 | 7.65E-285 | 5 | Vcam1 | 0.497 |
| 1.01E-271 | 0.9907469 | 0.752 | 0.233 | 3.13E-267 | 5 | Slco2b1 | 0.519 |
| 1.35E-266 | 2.2356845 | 0.821 | 0.375 | 4.19E-262 | 5 | Hspa1a | 0.446 |
| 7.26E-264 | 0.8012212 | 0.464 | 0.093 | 2.25E-259 | 5 | Serpinf1 | 0.371 |
| 1.53E-261 | 1.0574118 | 1 | 0.716 | 4.75E-257 | 5 | C1qc | 0.284 |
| 5.84E-260 | 0.9476956 | 0.646 | 0.178 | 1.81E-255 | 5 | Hpgds | 0.468 |
| 2.35E-258 | 0.6288856 | 0.234 | 0.023 | 7.31E-254 | 5 | Slc7a13 | 0.211 |
| 7.96E-238 | 1.1300244 | 0.726 | 0.237 | 2.47E-233 | 5 | Pla2g2d | 0.489 |
| 1.04E-230 | 0.9929462 | 0.991 | 0.675 | 3.23E-226 | 5 | Csf1r | 0.316 |
| 3.52E-230 | 2.3013021 | 0.779 | 0.358 | 1.09E-225 | 5 | Hspa1b | 0.421 |
| 3.69E-230 | 0.6173129 | 0.341 | 0.057 | 1.15E-225 | 5 | Mylk | 0.284 |
| 2.98E-229 | 1.6813962 | 0.628 | 0.195 | 9.24E-225 | 5 | Ccl12 | 0.433 |
| 1.68E-228 | 0.7952936 | 0.597 | 0.166 | 5.21E-224 | 5 | Tanc2 | 0.431 |
| 7.31E-228 | 0.9270164 | 0.588 | 0.163 | 2.27E-223 | 5 | Cp | 0.425 |
| 9.67E-224 | 1.0869208 | 0.356 | 0.065 | 3.00E-219 | 5 | Slc40a1 | 0.291 |
| 8.67E-222 | 0.9403971 | 0.685 | 0.222 | 2.69E-217 | 5 | Hpgd | 0.463 |
| 1.40E-215 | 1.0142419 | 0.936 | 0.44 | 4.35E-211 | 5 | Cbr2 | 0.496 |
| 1.42E-215 | 1.1271327 | 0.9 | 0.412 | 4.42E-211 | 5 | Stab1 | 0.488 |
| 2.39E-208 | 1.1003691 | 0.848 | 0.372 | 7.41E-204 | 5 | Cd163 | 0.476 |
| 4.65E-208 | 0.946942 | 0.807 | 0.318 | 1.44E-203 | 5 | Ms4a7 | 0.489 |
| 5.21E-208 | 0.6171967 | 0.314 | 0.053 | 1.62E-203 | 5 | Dnase1l3 | 0.261 |
| 3.41E-204 | 0.7580588 | 0.399 | 0.087 | 1.06E-199 | 5 | Slc15a2 | 0.312 |
| 9.55E-203 | 0.777986 | 0.562 | 0.166 | 2.96E-198 | 5 | Bank1 | 0.396 |
| 5.70E-201 | 1.006064 | 0.963 | 0.592 | 1.77E-196 | 5 | Trf | 0.371 |
| 1.84E-200 | 0.929914 | 0.999 | 0.699 | 5.71E-196 | 5 | C1qb | 0.3 |
| 4.79E-200 | 1.0438408 | 0.773 | 0.305 | 1.49E-195 | 5 | Mmp9 | 0.468 |
| 1.72E-198 | 1.0659664 | 0.998 | 0.812 | 5.34E-194 | 5 | Apoe | 0.186 |
| 1.48E-196 | 1.2700961 | 0.716 | 0.282 | 4.59E-192 | 5 | C4b | 0.434 |
| 4.74E-196 | 0.9460266 | 0.954 | 0.561 | 1.47E-191 | 5 | Cd81 | 0.393 |
| 1.49E-192 | 0.9781641 | 0.926 | 0.491 | 4.62E-188 | 5 | C5ar1 | 0.435 |
| 2.16E-192 | 0.967769 | 0.992 | 0.698 | 6.72E-188 | 5 | Dab2 | 0.294 |
| 1.01E-184 | 0.8080833 | 0.999 | 0.834 | 3.15E-180 | 5 | Mrc1 | 0.165 |
| 1.96E-178 | 0.572791 | 0.311 | 0.06 | 6.09E-174 | 5 | Siglec1 | 0.251 |
| 4.01E-177 | 0.4966905 | 0.294 | 0.054 | 1.25E-172 | 5 | Myh10 | 0.24 |
| 6.21E-177 | 1.1413887 | 0.914 | 0.557 | 1.93E-172 | 5 | Mgl2 | 0.357 |
| 2.96E-176 | 1.0056542 | 0.743 | 0.307 | 9.18E-172 | 5 | Pou2f2 | 0.436 |
| 4.91E-175 | 0.6651058 | 1 | 0.943 | 1.52E-170 | 5 | Cst3 | 0.057 |
| 1.39E-172 | 0.8248525 | 1 | 0.746 | 4.32E-168 | 5 | C1qa | 0.254 |
| 1.93E-172 | 0.4937676 | 0.348 | 0.076 | 5.98E-168 | 5 | Gpx3 | 0.272 |
| 1.30E-169 | 0.8595115 | 0.975 | 0.889 | 4.05E-165 | 5 | H2-Eb1 | 0.086 |
| 3.10E-169 | 1.5524552 | 0.529 | 0.186 | 9.61E-165 | 5 | Dnajb1 | 0.343 |
| 4.11E-169 | 0.7979769 | 0.611 | 0.209 | 1.28E-164 | 5 | Cd38 | 0.402 |
| 4.23E-163 | 0.8621404 | 0.998 | 0.673 | 1.31E-158 | 5 | Pf4 | 0.325 |
| 1.06E-162 | 0.628893 | 0.425 | 0.116 | 3.30E-158 | 5 | Aoah | 0.309 |
| 4.47E-160 | 0.5185526 | 0.344 | 0.079 | 1.39E-155 | 5 | Smpdl3b | 0.265 |
| 4.85E-160 | 0.7147849 | 0.999 | 0.866 | 1.51E-155 | 5 | Serinc3 | 0.133 |
| 5.87E-160 | 0.9098634 | 0.694 | 0.296 | 1.82E-155 | 5 | Cd86 | 0.398 |
| 1.46E-159 | 0.7063046 | 0.999 | 0.938 | 4.55E-155 | 5 | Ftl1 | 0.061 |
| 5.31E-159 | 0.8006411 | 0.827 | 0.377 | 1.65E-154 | 5 | Igf1 | 0.45 |
| 3.24E-155 | 0.8126484 | 0.863 | 0.438 | 1.01E-150 | 5 | C3ar1 | 0.425 |
| 3.85E-151 | 0.6982294 | 0.992 | 0.934 | 1.20E-146 | 5 | Cd74 | 0.058 |
| 4.70E-148 | 0.8085604 | 0.944 | 0.588 | 1.46E-143 | 5 | Adgre1 | 0.356 |

| | | | | | | | |
|---|---|---|---|---|---|---|---|
| 9.37E-145 | 0.6743093 | 0.623 | 0.239 | 2.91E-140 | 5 | Slc9a9 | 0.384 |
| 9.89E-143 | 0.8990591 | 0.939 | 0.613 | 3.07E-138 | 5 | Marcksl1 | 0.326 |
| 4.55E-142 | 0.6564985 | 0.739 | 0.32 | 1.41E-137 | 5 | Tgfbr2 | 0.419 |
| 1.41E-141 | 0.5278161 | 0.509 | 0.166 | 4.37E-137 | 5 | Cxcl12 | 0.343 |
| 5.56E-141 | 0.7429415 | 0.764 | 0.359 | 1.73E-136 | 5 | Cd300ld | 0.405 |
| 3.61E-140 | 0.8002299 | 0.981 | 0.644 | 1.12E-135 | 5 | Maf | 0.337 |
| 6.88E-140 | 0.6489572 | 0.608 | 0.233 | 2.14E-135 | 5 | P2ry6 | 0.375 |
| 1.30E-139 | 0.7891986 | 0.872 | 0.494 | 4.03E-135 | 5 | Clec4a1 | 0.378 |
| 1.70E-135 | 0.7502623 | 0.881 | 0.51 | 5.28E-131 | 5 | Fcgr3 | 0.371 |
| 9.41E-135 | 0.6663633 | 0.942 | 0.523 | 2.92E-130 | 5 | Wwp1 | 0.419 |
| 1.80E-132 | 0.6791819 | 0.947 | 0.507 | 5.59E-128 | 5 | Gas6 | 0.44 |
| 2.33E-132 | 0.6091606 | 0.648 | 0.262 | 7.24E-128 | 5 | Itsn1 | 0.386 |
| 4.39E-132 | 0.641577 | 0.998 | 0.89 | 1.36E-127 | 5 | Ctsb | 0.108 |
| 7.45E-132 | 0.7097936 | 1 | 0.806 | 2.31E-127 | 5 | Selenop | 0.194 |
| 1.15E-131 | 0.5643701 | 0.434 | 0.135 | 3.57E-127 | 5 | C5ar2 | 0.299 |
| 1.70E-131 | 0.6318263 | 0.519 | 0.188 | 5.29E-127 | 5 | Adap2 | 0.331 |
| 3.51E-131 | 0.6891963 | 0.773 | 0.377 | 1.09E-126 | 5 | Man1a | 0.396 |
| 8.43E-131 | 0.5360245 | 0.403 | 0.121 | 2.62E-126 | 5 | Siglece | 0.282 |
| 1.81E-130 | 0.6826404 | 0.422 | 0.132 | 5.63E-126 | 5 | Creb5 | 0.29 |
| 2.31E-129 | 0.6736015 | 0.68 | 0.297 | 7.16E-125 | 5 | Blvrb | 0.383 |
| 4.89E-129 | 1.0307217 | 0.854 | 0.492 | 1.52E-124 | 5 | Sdc4 | 0.362 |
| 9.83E-129 | 0.4630804 | 0.373 | 0.105 | 3.05E-124 | 5 | Maoa | 0.268 |
| 6.22E-128 | 0.6402639 | 0.989 | 0.925 | 1.93E-123 | 5 | H2-Aa | 0.064 |
| 8.48E-128 | 0.4499036 | 0.286 | 0.067 | 2.63E-123 | 5 | Samd4 | 0.219 |
| 1.39E-127 | 0.7071007 | 0.916 | 0.542 | 4.31E-123 | 5 | Mef2c | 0.374 |
| 4.43E-126 | 0.6434588 | 0.999 | 0.833 | 1.38E-121 | 5 | Ctsc | 0.166 |
| 1.62E-125 | 0.4200387 | 0.249 | 0.053 | 5.04E-121 | 5 | Tmod1 | 0.196 |
| 2.46E-123 | 1.1054566 | 0.824 | 0.507 | 7.64E-119 | 5 | Ier3 | 0.317 |
| 3.48E-121 | 0.6700585 | 0.718 | 0.333 | 1.08E-116 | 5 | Cxcl16 | 0.385 |
| 5.24E-120 | 1.3754077 | 0.823 | 0.521 | 1.63E-115 | 5 | Ccl2 | 0.302 |
| 9.68E-119 | 0.5581823 | 1 | 0.933 | 3.01E-114 | 5 | Itm2b | 0.067 |
| 6.85E-118 | 0.4880481 | 0.55 | 0.204 | 2.13E-113 | 5 | Reps2 | 0.346 |
| 9.37E-117 | 0.6327219 | 0.756 | 0.363 | 2.91E-112 | 5 | Lst1 | 0.393 |
| 1.21E-116 | 0.6016343 | 0.592 | 0.24 | 3.74E-112 | 5 | Lair1 | 0.352 |
| 6.40E-115 | 0.5439401 | 0.598 | 0.245 | 1.99E-110 | 5 | Psd3 | 0.353 |
| 7.48E-115 | 0.5960689 | 0.989 | 0.803 | 2.32E-110 | 5 | Cltc | 0.186 |
| 1.34E-114 | 0.5201591 | 0.466 | 0.162 | 4.15E-110 | 5 | Gpr34 | 0.304 |
| 2.37E-114 | 0.5310545 | 0.417 | 0.142 | 7.36E-110 | 5 | Mknk1 | 0.275 |
| 3.28E-114 | 0.5239198 | 0.71 | 0.314 | 1.02E-109 | 5 | Cd93 | 0.396 |
| 7.53E-114 | 0.432231 | 0.258 | 0.061 | 2.34E-109 | 5 | Adam33 | 0.197 |
| 1.22E-113 | 0.7205553 | 0.744 | 0.383 | 3.78E-109 | 5 | Cd68 | 0.361 |
| 3.25E-111 | 0.5465882 | 0.481 | 0.176 | 1.01E-106 | 5 | Fcgr1 | 0.305 |
| 4.17E-111 | 0.4686466 | 0.507 | 0.189 | 1.30E-106 | 5 | Nfxl1 | 0.318 |
| 6.19E-111 | 0.6751722 | 0.992 | 0.909 | 1.92E-106 | 5 | Hspa8 | 0.083 |
| 4.02E-110 | 0.5792664 | 0.8 | 0.404 | 1.25E-105 | 5 | Pmp22 | 0.396 |
| 2.70E-109 | 0.6686653 | 0.908 | 0.584 | 8.39E-105 | 5 | Ehd4 | 0.324 |
| 1.36E-108 | 1.0205861 | 0.556 | 0.244 | 4.23E-104 | 5 | Nlrp3 | 0.312 |
| 4.97E-108 | 0.5430444 | 0.498 | 0.198 | 1.54E-103 | 5 | Ftl1-ps1 | 0.3 |
| 9.59E-108 | 0.4897736 | 0.387 | 0.126 | 2.98E-103 | 5 | Tlr7 | 0.261 |
| 1.72E-107 | 0.6079227 | 0.903 | 0.503 | 5.33E-103 | 5 | Igfbp4 | 0.4 |
| 6.41E-107 | 0.5023901 | 0.998 | 0.93 | 1.99E-102 | 5 | B2m | 0.068 |
| 3.27E-106 | 0.6170932 | 0.586 | 0.262 | 1.01E-101 | 5 | Lacc1 | 0.324 |
| 8.14E-106 | 0.5528475 | 0.714 | 0.345 | 2.53E-101 | 5 | Abhd12 | 0.369 |
| 2.70E-105 | 1.3777142 | 0.356 | 0.119 | 8.40E-101 | 5 | Ccl3 | 0.237 |
| 1.10E-103 | 0.4351597 | 0.596 | 0.256 | 3.41E-99 | 5 | Basp1 | 0.34 |
| 2.26E-103 | 0.4732805 | 0.586 | 0.243 | 7.03E-99 | 5 | Ms4a6b | 0.343 |

| | | | | | | | |
|---|---|---|---|---|---|---|---|
| 2.60E-102 | 0.4924498 | 0.288 | 0.081 | 8.08E-98 | 5 | Fcgr4 | 0.207 |
| 4.20E-102 | 0.9577857 | 0.931 | 0.791 | 1.30E-97 | 5 | Cd83 | 0.14 |
| 5.53E-102 | 0.394864 | 0.372 | 0.121 | 1.72E-97 | 5 | Smagp | 0.251 |
| 6.69E-102 | 0.4284458 | 0.29 | 0.081 | 2.08E-97 | 5 | Cd72 | 0.209 |
| 1.14E-101 | 0.4602392 | 0.672 | 0.307 | 3.56E-97 | 5 | Gatm | 0.365 |
| 2.06E-100 | 0.5277672 | 0.697 | 0.339 | 6.40E-96 | 5 | Arrb2 | 0.358 |
| 4.89E-100 | 0.4731093 | 0.553 | 0.23 | 1.52E-95 | 5 | Tcn2 | 0.323 |
| 8.90E-100 | 0.425499 | 0.35 | 0.11 | 2.76E-95 | 5 | Sema6d | 0.24 |
| 1.83E-99 | 1.1387033 | 0.7 | 0.378 | 5.68E-95 | 5 | Ccl7 | 0.322 |
| 1.83E-99 | 0.5064008 | 0.767 | 0.387 | 5.68E-95 | 5 | Dse | 0.38 |
| 2.22E-99 | 0.6703503 | 0.735 | 0.385 | 6.89E-95 | 5 | AC149090.1 | 0.35 |
| 4.64E-98 | 0.674524 | 0.641 | 0.306 | 1.44E-93 | 5 | Ccnd1 | 0.335 |
| 9.01E-98 | 0.6062488 | 0.268 | 0.073 | 2.80E-93 | 5 | Siglech | 0.195 |
| 8.40E-97 | 0.3860672 | 0.325 | 0.101 | 2.61E-92 | 5 | Ctsf | 0.224 |
| 1.68E-96 | 0.3135925 | 0.165 | 0.031 | 5.23E-92 | 5 | Tpbgl | 0.134 |
| 3.26E-96 | 0.6371831 | 0.813 | 0.471 | 1.01E-91 | 5 | Clec2d | 0.342 |
| 7.66E-96 | 1.4182872 | 0.493 | 0.215 | 2.38E-91 | 5 | Ccl4 | 0.278 |
| 9.55E-95 | 0.4834388 | 0.668 | 0.313 | 2.97E-90 | 5 | Irf8 | 0.355 |
| 1.19E-94 | 0.435653 | 0.447 | 0.171 | 3.68E-90 | 5 | Clcn5 | 0.276 |
| 1.55E-94 | 0.4108983 | 0.304 | 0.092 | 4.81E-90 | 5 | P3h2 | 0.212 |
| 1.43E-93 | 0.3480943 | 0.337 | 0.108 | 4.45E-89 | 5 | Trpv4 | 0.229 |
| 2.80E-93 | 0.3793743 | 0.25 | 0.067 | 8.70E-89 | 5 | Dusp8 | 0.183 |
| 3.65E-93 | 0.5883781 | 0.722 | 0.388 | 1.13E-88 | 5 | Runx1 | 0.334 |
| 6.50E-93 | 0.4666172 | 0.671 | 0.327 | 2.02E-88 | 5 | Il10rb | 0.344 |
| 9.03E-93 | 0.58417 | 0.367 | 0.132 | 2.80E-88 | 5 | Jdp2 | 0.235 |
| 5.53E-92 | 0.5701216 | 0.846 | 0.457 | 1.72E-87 | 5 | Fcrls | 0.389 |
| 9.61E-92 | 0.4076797 | 1 | 0.996 | 2.98E-87 | 5 | mt-Atp6 | 0.004 |
| 2.77E-91 | 0.2940329 | 0.369 | 0.125 | 8.60E-87 | 5 | Dst | 0.244 |
| 4.05E-91 | 0.763186 | 0.641 | 0.322 | 1.26E-86 | 5 | Cited2 | 0.319 |
| 2.10E-90 | 0.4858198 | 0.828 | 0.471 | 6.52E-86 | 5 | Eps15 | 0.357 |
| 6.18E-90 | 0.3006847 | 0.194 | 0.044 | 1.92E-85 | 5 | Ppp1r9a | 0.15 |
| 2.90E-89 | 0.4826749 | 0.909 | 0.506 | 9.02E-85 | 5 | Cd63 | 0.403 |
| 4.64E-89 | 0.4242959 | 0.396 | 0.146 | 1.44E-84 | 5 | Etv5 | 0.25 |
| 6.11E-89 | 0.5296009 | 0.762 | 0.41 | 1.90E-84 | 5 | Clec12a | 0.352 |
| 7.14E-89 | 0.7108834 | 0.836 | 0.556 | 2.22E-84 | 5 | Vmp1 | 0.28 |
| 8.80E-89 | 0.3670999 | 0.232 | 0.061 | 2.73E-84 | 5 | Cdr2 | 0.171 |
| 1.64E-88 | 0.4552753 | 0.427 | 0.167 | 5.08E-84 | 5 | Ebi3 | 0.26 |
| 1.73E-88 | 0.4318678 | 0.413 | 0.157 | 5.36E-84 | 5 | P2rx7 | 0.256 |
| 1.79E-88 | 0.4949641 | 0.573 | 0.262 | 5.55E-84 | 5 | Rab3il1 | 0.311 |
| 2.01E-88 | 0.5132083 | 0.775 | 0.429 | 6.26E-84 | 5 | Asah1 | 0.346 |
| 3.01E-88 | 0.4094038 | 0.39 | 0.142 | 9.33E-84 | 5 | Itgb5 | 0.248 |
| 4.55E-88 | 0.5422275 | 0.899 | 0.592 | 1.41E-83 | 5 | Ctsa | 0.307 |
| 5.93E-87 | 0.3415362 | 0.267 | 0.078 | 1.84E-82 | 5 | Eef2k | 0.189 |
| 6.41E-87 | 0.5779203 | 0.944 | 0.637 | 1.99E-82 | 5 | Clec10a | 0.307 |
| 2.17E-86 | 0.5715827 | 0.963 | 0.69 | 6.75E-82 | 5 | Tmem176b | 0.273 |
| 3.03E-86 | 0.8636564 | 0.967 | 0.819 | 9.41E-82 | 5 | Ubc | 0.148 |
| 1.41E-85 | 0.4842541 | 0.522 | 0.232 | 4.38E-81 | 5 | Slc11a1 | 0.29 |
| 3.03E-85 | 0.6513827 | 0.977 | 0.919 | 9.41E-81 | 5 | Zfp36 | 0.058 |
| 4.05E-85 | 0.3024578 | 0.225 | 0.059 | 1.26E-80 | 5 | B3galnt1 | 0.166 |
| 4.89E-85 | 0.5015345 | 0.805 | 0.445 | 1.52E-80 | 5 | Ifi207 | 0.36 |
| 5.83E-84 | 0.5973603 | 0.748 | 0.424 | 1.81E-79 | 5 | Lilr4b | 0.324 |
| 6.06E-84 | 0.362788 | 0.433 | 0.167 | 1.88E-79 | 5 | Cmah | 0.266 |
| 9.69E-84 | 0.4862049 | 0.689 | 0.358 | 3.01E-79 | 5 | Bst2 | 0.331 |
| 1.16E-83 | 0.4078572 | 0.456 | 0.185 | 3.59E-79 | 5 | Adam19 | 0.271 |
| 1.18E-83 | 0.4349969 | 0.621 | 0.304 | 3.67E-79 | 5 | Tsc22d1 | 0.317 |
| 6.16E-83 | 0.6314415 | 0.973 | 0.772 | 1.91E-78 | 5 | Zfp36l1 | 0.201 |

| | | | | | | | |
|---|---|---|---|---|---|---|---|
| 1.01E-82 | 0.2499674 | 0.149 | 0.029 | 3.14E-78 | 5 | Epor | 0.12 |
| 1.95E-82 | 0.4439834 | 0.999 | 0.978 | 6.07E-78 | 5 | mt-Cytb | 0.021 |
| 2.17E-82 | 0.3121568 | 0.155 | 0.032 | 6.74E-78 | 5 | Rad51b | 0.123 |
| 1.47E-81 | 0.3966367 | 0.31 | 0.105 | 4.55E-77 | 5 | Zfp704 | 0.205 |
| 8.02E-81 | 0.4000534 | 0.65 | 0.319 | 2.49E-76 | 5 | Sec14l1 | 0.331 |
| 8.63E-81 | 0.3591209 | 0.297 | 0.098 | 2.68E-76 | 5 | Cyp27a1 | 0.199 |
| 3.21E-80 | 0.335634 | 0.178 | 0.041 | 9.97E-76 | 5 | Spic | 0.137 |
| 1.17E-79 | 0.4261969 | 0.997 | 0.905 | 3.63E-75 | 5 | Marcks | 0.092 |
| 1.18E-79 | 0.4862741 | 0.836 | 0.485 | 3.67E-75 | 5 | Cfh | 0.351 |
| 1.93E-79 | 0.4748937 | 0.758 | 0.424 | 5.99E-75 | 5 | Zfhx3 | 0.334 |
| 1.23E-78 | 0.6225509 | 0.889 | 0.573 | 3.81E-74 | 5 | Ly6e | 0.316 |
| 1.88E-78 | 0.3501511 | 0.325 | 0.114 | 5.84E-74 | 5 | Scamp5 | 0.211 |
| 6.91E-78 | 0.2557773 | 0.195 | 0.05 | 2.15E-73 | 5 | Wtip | 0.145 |
| 9.27E-78 | 0.5597943 | 0.778 | 0.433 | 2.88E-73 | 5 | Folr2 | 0.345 |
| 1.21E-77 | 0.7241609 | 0.774 | 0.523 | 3.77E-73 | 5 | Kdm6b | 0.251 |
| 1.25E-77 | 0.525445 | 0.973 | 0.777 | 3.87E-73 | 5 | Lcp1 | 0.196 |
| 2.91E-77 | 0.45416 | 0.726 | 0.395 | 9.02E-73 | 5 | Myo5a | 0.331 |
| 3.12E-77 | 0.41967 | 0.734 | 0.39 | 9.69E-73 | 5 | Rgs10 | 0.344 |
| 4.54E-77 | 0.3296967 | 0.273 | 0.087 | 1.41E-72 | 5 | Car3 | 0.186 |
| 5.25E-77 | 0.4231303 | 0.56 | 0.256 | 1.63E-72 | 5 | Cd33 | 0.304 |
| 5.76E-77 | 0.3167636 | 0.383 | 0.144 | 1.79E-72 | 5 | Prune2 | 0.239 |
| 1.18E-76 | 0.6458469 | 0.281 | 0.093 | 3.66E-72 | 5 | Cd209b | 0.188 |
| 1.49E-76 | 0.4408826 | 0.682 | 0.36 | 4.62E-72 | 5 | Aldh2 | 0.322 |
| 3.23E-76 | 0.3145657 | 0.284 | 0.092 | 1.00E-71 | 5 | Gprc5c | 0.192 |
| 3.25E-76 | 0.3912425 | 0.336 | 0.122 | 1.01E-71 | 5 | Tmem8 | 0.214 |
| 5.18E-76 | 0.4823561 | 0.667 | 0.338 | 1.61E-71 | 5 | Rnase4 | 0.329 |
| 1.65E-75 | 0.4034902 | 0.55 | 0.259 | 5.13E-71 | 5 | Sbf2 | 0.291 |
| 2.53E-75 | 0.608544 | 0.897 | 0.646 | 7.85E-71 | 5 | Ier5 | 0.251 |
| 6.79E-75 | 0.5071215 | 0.488 | 0.224 | 2.11E-70 | 5 | Slamf7 | 0.264 |
| 7.48E-75 | 0.4063145 | 0.435 | 0.187 | 2.32E-70 | 5 | Mob3c | 0.248 |
| 2.08E-74 | 0.3765825 | 0.431 | 0.178 | 6.46E-70 | 5 | Ms4a6d | 0.253 |
| 7.64E-74 | 0.3921696 | 0.454 | 0.201 | 2.37E-69 | 5 | Sh3pxd2b | 0.253 |
| 4.62E-73 | 0.4308922 | 0.444 | 0.193 | 1.43E-68 | 5 | Lgals3bp | 0.251 |
| 5.03E-73 | 0.4502809 | 0.968 | 0.747 | 1.56E-68 | 5 | Unc93b1 | 0.221 |
| 8.42E-73 | 0.3270754 | 0.308 | 0.109 | 2.61E-68 | 5 | Frmd6 | 0.199 |
| 1.17E-72 | 0.4656258 | 0.276 | 0.092 | 3.62E-68 | 5 | Clec4n | 0.184 |
| 2.37E-72 | 0.372639 | 1 | 0.974 | 7.36E-68 | 5 | mt-Nd4 | 0.026 |
| 4.35E-72 | 0.4490938 | 0.914 | 0.588 | 1.35E-67 | 5 | Glul | 0.326 |
| 7.87E-72 | 0.2985643 | 0.357 | 0.135 | 2.44E-67 | 5 | Ophn1 | 0.222 |
| 9.43E-72 | 0.40204 | 0.464 | 0.208 | 2.93E-67 | 5 | Dclre1c | 0.256 |
| 1.26E-71 | 0.3875956 | 0.263 | 0.086 | 3.92E-67 | 5 | Nxpe5 | 0.177 |
| 1.29E-71 | 0.4383364 | 0.681 | 0.359 | 4.01E-67 | 5 | Rgl1 | 0.322 |
| 2.68E-71 | 0.4936422 | 0.761 | 0.444 | 8.32E-67 | 5 | Rab7b | 0.317 |
| 2.78E-71 | 0.3646986 | 0.426 | 0.18 | 8.64E-67 | 5 | Rasa4 | 0.246 |
| 2.93E-71 | 0.380746 | 0.351 | 0.138 | 9.11E-67 | 5 | Lpin2 | 0.213 |
| 3.14E-71 | 1.1148667 | 0.773 | 0.567 | 9.75E-67 | 5 | Cxcl2 | 0.206 |
| 4.98E-71 | 0.7302767 | 0.886 | 0.602 | 1.55E-66 | 5 | Cd14 | 0.284 |
| 5.21E-71 | 0.2963611 | 0.38 | 0.15 | 1.62E-66 | 5 | Slc12a7 | 0.23 |
| 6.86E-71 | 0.3217578 | 0.394 | 0.16 | 2.13E-66 | 5 | Zbtb4 | 0.234 |
| 8.37E-71 | 0.4157749 | 0.396 | 0.168 | 2.60E-66 | 5 | Limd2 | 0.228 |
| 9.58E-71 | 0.6116377 | 0.937 | 0.782 | 2.98E-66 | 5 | Nfkbiz | 0.155 |
| 1.08E-70 | 0.35286 | 0.534 | 0.251 | 3.35E-66 | 5 | Cmklr1 | 0.283 |
| 1.11E-70 | 0.454048 | 0.405 | 0.171 | 3.45E-66 | 5 | Cdk6 | 0.234 |
| 2.35E-70 | 0.4045776 | 0.553 | 0.269 | 7.29E-66 | 5 | Rab11fip5 | 0.284 |
| 2.51E-70 | 0.2676779 | 0.288 | 0.1 | 7.80E-66 | 5 | Retreg1 | 0.188 |
| 4.66E-70 | 0.2676926 | 0.23 | 0.07 | 1.45E-65 | 5 | Csf3r | 0.16 |

| | | | | | | | |
|---|---|---|---|---|---|---|---|
| 8.58E-70 | 0.4664882 | 0.744 | 0.423 | 2.66E-65 | 5 | Fyb | 0.321 |
| 2.23E-69 | 0.2994843 | 0.327 | 0.121 | 6.92E-65 | 5 | Ralgps2 | 0.206 |
| 2.42E-69 | 0.7020265 | 0.331 | 0.128 | 7.52E-65 | 5 | Bmp2 | 0.203 |
| 2.82E-69 | 0.3822276 | 0.598 | 0.303 | 8.77E-65 | 5 | Gna12 | 0.295 |
| 3.92E-69 | 0.3816513 | 0.406 | 0.17 | 1.22E-64 | 5 | Snx29 | 0.236 |
| 5.73E-69 | 0.1733953 | 0.953 | 0.666 | 1.78E-64 | 5 | Cd36 | 0.287 |
| 1.04E-68 | 0.3548506 | 0.451 | 0.195 | 3.22E-64 | 5 | Msr1 | 0.256 |
| 1.15E-68 | 0.4661351 | 0.578 | 0.298 | 3.57E-64 | 5 | Peli1 | 0.28 |
| 1.26E-68 | 0.3459091 | 0.535 | 0.254 | 3.91E-64 | 5 | Prkcb | 0.281 |
| 1.54E-68 | 0.3338754 | 0.538 | 0.26 | 4.78E-64 | 5 | Swap70 | 0.278 |
| 2.67E-68 | 0.3978836 | 0.534 | 0.261 | 8.29E-64 | 5 | Sft2d2 | 0.273 |
| 4.85E-68 | 0.5334237 | 0.929 | 0.708 | 1.51E-63 | 5 | Egr1 | 0.221 |
| 5.94E-68 | 0.3896649 | 0.556 | 0.276 | 1.84E-63 | 5 | Man1c1 | 0.28 |
| 1.27E-67 | 0.3845972 | 1 | 0.961 | 3.94E-63 | 5 | mt-Nd2 | 0.039 |
| 2.88E-67 | 0.2728178 | 0.221 | 0.067 | 8.93E-63 | 5 | Jade2 | 0.154 |
| 3.41E-67 | 0.31011 | 0.437 | 0.19 | 1.06E-62 | 5 | Il13ra1 | 0.247 |
| 1.28E-66 | 0.4264947 | 0.788 | 0.492 | 3.96E-62 | 5 | Pea15a | 0.296 |
| 2.34E-66 | 0.2222474 | 0.152 | 0.036 | 7.28E-62 | 5 | Cables1 | 0.116 |
| 3.32E-66 | 0.3216951 | 0.731 | 0.39 | 1.03E-61 | 5 | Frmd4b | 0.341 |
| 3.51E-66 | 0.4381423 | 0.406 | 0.176 | 1.09E-61 | 5 | Adrb2 | 0.23 |
| 4.38E-66 | 0.3265985 | 0.73 | 0.403 | 1.36E-61 | 5 | Ap1b1 | 0.327 |
| 1.26E-65 | 0.3678948 | 0.58 | 0.29 | 3.91E-61 | 5 | Rnf150 | 0.29 |
| 1.52E-65 | 0.3862264 | 0.604 | 0.32 | 4.72E-61 | 5 | Lmo2 | 0.284 |
| 2.47E-65 | 0.2631288 | 0.133 | 0.029 | 7.68E-61 | 5 | Sgce | 0.104 |
| 2.96E-65 | 0.4546498 | 0.981 | 0.917 | 9.19E-61 | 5 | H2-Ab1 | 0.064 |
| 3.66E-65 | 0.292751 | 0.231 | 0.073 | 1.14E-60 | 5 | Oas2 | 0.158 |
| 3.74E-65 | 0.3623524 | 0.886 | 0.551 | 1.16E-60 | 5 | Fcgrt | 0.335 |
| 1.39E-64 | 0.279238 | 0.212 | 0.065 | 4.32E-60 | 5 | Rnaset2a | 0.147 |
| 2.57E-64 | 0.3821758 | 0.3 | 0.115 | 7.99E-60 | 5 | Pdcd2l | 0.185 |
| 1.05E-63 | 0.2865196 | 0.262 | 0.091 | 3.26E-59 | 5 | Gaa | 0.171 |
| 1.19E-63 | 0.3973579 | 0.258 | 0.09 | 3.71E-59 | 5 | Plau | 0.168 |
| 1.35E-63 | 0.1987184 | 0.159 | 0.04 | 4.18E-59 | 5 | Rac3 | 0.119 |
| 2.49E-63 | 1.0550434 | 0.245 | 0.085 | 7.73E-59 | 5 | Gdf15 | 0.16 |
| 2.64E-63 | 0.2784687 | 0.261 | 0.09 | 8.21E-59 | 5 | Npl | 0.171 |
| 4.16E-63 | 0.3821512 | 0.712 | 0.405 | 1.29E-58 | 5 | Tpp1 | 0.307 |
| 9.35E-63 | 0.3114219 | 0.277 | 0.1 | 2.90E-58 | 5 | Arhgap19 | 0.177 |
| 9.76E-63 | 0.2442804 | 0.15 | 0.037 | 3.03E-58 | 5 | Asgr2 | 0.113 |
| 1.08E-62 | 0.2431099 | 0.171 | 0.047 | 3.37E-58 | 5 | Rnaset2b | 0.124 |
| 1.33E-62 | 0.1800914 | 0.122 | 0.026 | 4.12E-58 | 5 | Unc5a | 0.096 |
| 1.33E-62 | 0.3316644 | 0.409 | 0.179 | 4.12E-58 | 5 | Mertk | 0.23 |
| 1.45E-62 | 0.3864232 | 0.661 | 0.36 | 4.49E-58 | 5 | Hacd4 | 0.301 |
| 1.91E-62 | 0.4048445 | 0.369 | 0.159 | 5.92E-58 | 5 | Tnf | 0.21 |
| 2.09E-62 | 0.3855309 | 0.272 | 0.098 | 6.50E-58 | 5 | Gpr183 | 0.174 |
| 5.63E-62 | 0.3833138 | 0.547 | 0.282 | 1.75E-57 | 5 | Cd302 | 0.265 |
| 7.98E-62 | 0.3137457 | 0.575 | 0.291 | 2.48E-57 | 5 | Acp2 | 0.284 |
| 1.17E-61 | 0.263359 | 0.179 | 0.05 | 3.63E-57 | 5 | Cnrip1 | 0.129 |
| 1.71E-61 | 0.4533477 | 0.977 | 0.864 | 5.30E-57 | 5 | mt-Nd3 | 0.113 |
| 1.72E-61 | 0.3382696 | 0.662 | 0.368 | 5.34E-57 | 5 | ;10507B11R | 0.294 |
| 2.04E-61 | 0.3635107 | 0.262 | 0.093 | 6.35E-57 | 5 | Ang | 0.169 |
| 3.60E-61 | 0.2851126 | 0.276 | 0.102 | 1.12E-56 | 5 | Selenon | 0.174 |
| 1.07E-60 | 0.3136812 | 0.395 | 0.173 | 3.33E-56 | 5 | Cttnbp2nl | 0.222 |
| 1.35E-60 | 0.3060424 | 0.281 | 0.105 | 4.20E-56 | 5 | Adgre4 | 0.176 |
| 1.40E-60 | 0.2307849 | 0.557 | 0.277 | 4.35E-56 | 5 | Kitl | 0.28 |
| 1.49E-60 | 0.6557312 | 0.581 | 0.313 | 4.64E-56 | 5 | Cd209f | 0.268 |
| 1.59E-60 | 0.3459021 | 0.645 | 0.353 | 4.93E-56 | 5 | Spred1 | 0.292 |
| 2.39E-60 | 0.382044 | 0.747 | 0.445 | 7.41E-56 | 5 | Tmem106a | 0.302 |

| | | | | | | | |
|---|---|---|---|---|---|---|---|
| 3.49E-60 | 0.1881878 | 0.378 | 0.159 | 1.08E-55 | 5 | Gstm1 | 0.219 |
| 4.17E-60 | 0.3741785 | 0.211 | 0.068 | 1.30E-55 | 5 | Gm48065 | 0.143 |
| 4.21E-60 | 0.3747187 | 0.982 | 0.819 | 1.31E-55 | 5 | Grn | 0.163 |
| 4.43E-60 | 0.3103406 | 0.308 | 0.119 | 1.37E-55 | 5 | Itga9 | 0.189 |
| 6.21E-60 | 0.4447655 | 0.195 | 0.06 | 1.93E-55 | 5 | Hbegf | 0.135 |
| 1.06E-59 | 0.3299481 | 0.44 | 0.21 | 3.30E-55 | 5 | Coro1b | 0.23 |
| 1.43E-59 | 0.3918017 | 0.843 | 0.506 | 4.43E-55 | 5 | Aif1 | 0.337 |
| 1.58E-59 | 0.2752701 | 0.31 | 0.121 | 4.91E-55 | 5 | Rhobtb1 | 0.189 |
| 2.28E-59 | 0.3798212 | 0.603 | 0.329 | 7.09E-55 | 5 | Tfe3 | 0.274 |
| 3.40E-59 | 0.278822 | 0.549 | 0.281 | 1.05E-54 | 5 | Grina | 0.268 |
| 3.48E-59 | 0.3017186 | 0.325 | 0.133 | 1.08E-54 | 5 | Ly96 | 0.192 |
| 4.40E-59 | 0.8166489 | 0.573 | 0.31 | 1.36E-54 | 5 | Ccl8 | 0.263 |
| 7.04E-59 | 0.4886598 | 0.507 | 0.252 | 2.19E-54 | 5 | Cd209g | 0.255 |
| 7.82E-59 | 0.3168546 | 0.465 | 0.226 | 2.43E-54 | 5 | Cat | 0.239 |
| 9.06E-59 | 0.344872 | 0.484 | 0.24 | 2.81E-54 | 5 | Neu1 | 0.244 |
| 1.03E-58 | 0.2800976 | 0.2 | 0.063 | 3.21E-54 | 5 | Utp14b | 0.137 |
| 1.11E-58 | 0.4209898 | 0.468 | 0.233 | 3.45E-54 | 5 | Ppp1r10 | 0.235 |
| 1.33E-58 | 0.3115945 | 0.744 | 0.431 | 4.12E-54 | 5 | Nrp1 | 0.313 |
| 1.43E-58 | 0.497206 | 0.537 | 0.286 | 4.45E-54 | 5 | AC160336.1 | 0.251 |
| 2.27E-58 | 0.3092406 | 0.283 | 0.109 | 7.04E-54 | 5 | Il16 | 0.174 |
| 2.64E-58 | 0.3604134 | 0.438 | 0.21 | 8.20E-54 | 5 | Antxr2 | 0.228 |
| 2.70E-58 | 0.4137248 | 0.777 | 0.467 | 8.39E-54 | 5 | Man2b1 | 0.31 |
| 5.52E-58 | 0.2949221 | 0.218 | 0.072 | 1.72E-53 | 5 | Kansl1l | 0.146 |
| 6.31E-58 | 0.2572713 | 0.327 | 0.131 | 1.96E-53 | 5 | Ggt5 | 0.196 |
| 6.49E-58 | 0.2635238 | 0.364 | 0.153 | 2.02E-53 | 5 | Etv1 | 0.211 |
| 9.07E-58 | 0.6458943 | 0.734 | 0.484 | 2.82E-53 | 5 | Gadd45b | 0.25 |
| 1.25E-57 | 0.405227 | 0.854 | 0.56 | 3.90E-53 | 5 | Tm6sf1 | 0.294 |
| 1.26E-57 | 0.1765458 | 0.115 | 0.025 | 3.91E-53 | 5 | Tnfrsf17 | 0.09 |
| 1.57E-57 | 0.2078495 | 0.179 | 0.052 | 4.86E-53 | 5 | Scd1 | 0.127 |
| 2.84E-57 | 0.4432699 | 0.507 | 0.266 | 8.81E-53 | 5 | Fgl2 | 0.241 |
| 3.60E-57 | 0.2686118 | 0.254 | 0.092 | 1.12E-52 | 5 | BC035044 | 0.162 |
| 3.90E-57 | 0.3093503 | 0.739 | 0.429 | 1.21E-52 | 5 | Laptm4a | 0.31 |
| 4.70E-57 | 0.3022646 | 0.396 | 0.18 | 1.46E-52 | 5 | Pik3cg | 0.216 |
| 4.92E-57 | 0.2547709 | 0.627 | 0.324 | 1.53E-52 | 5 | Ifi204 | 0.303 |
| 4.97E-57 | 0.4122792 | 0.691 | 0.406 | 1.54E-52 | 5 | Spi1 | 0.285 |
| 1.04E-56 | 0.2433643 | 0.157 | 0.043 | 3.22E-52 | 5 | Scimp | 0.114 |
| 1.23E-56 | 0.2332439 | 0.226 | 0.077 | 3.81E-52 | 5 | Snx24 | 0.149 |
| 1.39E-56 | 0.3290239 | 0.736 | 0.429 | 4.30E-52 | 5 | Nr3c1 | 0.307 |
| 1.77E-56 | 0.2299303 | 0.533 | 0.266 | 5.49E-52 | 5 | Sult1a1 | 0.267 |
| 2.77E-56 | 0.3463668 | 0.728 | 0.429 | 8.61E-52 | 5 | Nisch | 0.299 |
| 5.52E-56 | 0.3187929 | 0.482 | 0.236 | 1.71E-51 | 5 | Pla2g15 | 0.246 |
| 6.43E-56 | 0.3771341 | 0.632 | 0.353 | 2.00E-51 | 5 | Pla2g7 | 0.279 |
| 7.25E-56 | 0.2479299 | 0.259 | 0.096 | 2.25E-51 | 5 | Agpat3 | 0.163 |
| 1.39E-55 | 0.3295359 | 0.436 | 0.211 | 4.32E-51 | 5 | Ncoa4 | 0.225 |
| 1.82E-55 | 0.3116401 | 0.521 | 0.27 | 5.65E-51 | 5 | Dgkd | 0.251 |
| 3.91E-55 | 0.3520985 | 0.459 | 0.222 | 1.21E-50 | 5 | Tfrc | 0.237 |
| 4.57E-55 | 0.3428727 | 0.504 | 0.264 | 1.42E-50 | 5 | Lrrc25 | 0.24 |
| 6.91E-55 | 0.2201546 | 0.248 | 0.089 | 2.14E-50 | 5 | Sh3pxd2a | 0.159 |
| 1.11E-54 | 0.4139967 | 0.715 | 0.443 | 3.44E-50 | 5 | Slc3a2 | 0.272 |
| 3.69E-54 | 0.3241526 | 0.568 | 0.299 | 1.15E-49 | 5 | Axl | 0.269 |
| 7.41E-54 | 0.3504545 | 0.848 | 0.548 | 2.30E-49 | 5 | Snx5 | 0.3 |
| 7.52E-54 | 0.2143446 | 0.352 | 0.149 | 2.34E-49 | 5 | Chp2 | 0.203 |
| 7.99E-54 | 0.3795356 | 0.922 | 0.644 | 2.48E-49 | 5 | Tmem176a | 0.278 |
| 8.43E-54 | 0.3357757 | 0.356 | 0.159 | 2.62E-49 | 5 | Bcl6 | 0.197 |
| 1.41E-53 | 0.3119994 | 0.327 | 0.139 | 4.36E-49 | 5 | Trim47 | 0.188 |
| 1.90E-53 | 0.2842847 | 0.496 | 0.251 | 5.91E-49 | 5 | Wdfy3 | 0.245 |

| | | | | | | | |
|---|---|---|---|---|---|---|---|
| 2.43E-53 | 0.289425 | 0.437 | 0.214 | 7.53E-49 | 5 | Ppt1 | 0.223 |
| 2.89E-53 | 0.3481707 | 0.661 | 0.381 | 8.97E-49 | 5 | Lipa | 0.28 |
| 3.92E-53 | 0.2741821 | 0.443 | 0.218 | 1.22E-48 | 5 | Trim41 | 0.225 |
| 4.53E-53 | 0.3163791 | 0.514 | 0.271 | 1.41E-48 | 5 | Fgd2 | 0.243 |
| 5.04E-53 | 0.3388591 | 0.742 | 0.448 | 1.56E-48 | 5 | Gns | 0.294 |
| 6.52E-53 | 0.2068556 | 0.291 | 0.114 | 2.02E-48 | 5 | A4galt | 0.177 |
| 9.50E-53 | 0.2616976 | 0.463 | 0.228 | 2.95E-48 | 5 | Tceal9 | 0.235 |
| 1.01E-52 | 0.2831304 | 0.233 | 0.084 | 3.15E-48 | 5 | Slc14a1 | 0.149 |
| 1.40E-52 | 0.251977 | 0.666 | 0.368 | 4.35E-48 | 5 | Stard8 | 0.298 |
| 1.50E-52 | 0.3245394 | 0.734 | 0.421 | 4.66E-48 | 5 | Ninj1 | 0.313 |
| 1.54E-52 | 0.8606745 | 0.846 | 0.619 | 4.79E-48 | 5 | Hsp90aa1 | 0.227 |
| 1.74E-52 | 0.2934654 | 0.467 | 0.237 | 5.41E-48 | 5 | Rela | 0.23 |
| 1.74E-52 | 0.2942306 | 0.432 | 0.207 | 5.41E-48 | 5 | Fchsd2 | 0.225 |
| 2.31E-52 | 0.333436 | 0.777 | 0.483 | 7.16E-48 | 5 | Ptprj | 0.294 |
| 2.50E-52 | 0.3731423 | 0.989 | 0.886 | 7.76E-48 | 5 | H2-D1 | 0.103 |
| 2.64E-52 | 0.2303688 | 0.124 | 0.031 | 8.20E-48 | 5 | Slc25a33 | 0.093 |
| 2.98E-52 | 0.3674116 | 0.605 | 0.338 | 9.27E-48 | 5 | Rnasel | 0.267 |
| 4.29E-52 | 0.2859956 | 0.627 | 0.35 | 1.33E-47 | 5 | Epsti1 | 0.277 |
| 6.69E-52 | 0.2065005 | 0.846 | 0.564 | 2.08E-47 | 5 | Ubb | 0.282 |
| 8.01E-52 | 0.3677479 | 0.721 | 0.437 | 2.49E-47 | 5 | Lifr | 0.284 |
| 8.84E-52 | 0.1952943 | 0.121 | 0.029 | 2.75E-47 | 5 | Ifi213 | 0.092 |
| 8.97E-52 | 0.4828035 | 0.483 | 0.259 | 2.78E-47 | 5 | Gpr65 | 0.224 |
| 1.41E-51 | 0.2677753 | 0.226 | 0.082 | 4.39E-47 | 5 | Bcl2 | 0.144 |
| 2.36E-51 | 0.344626 | 0.547 | 0.298 | 7.33E-47 | 5 | Fam129a | 0.249 |
| 3.29E-51 | 0.3536506 | 0.582 | 0.33 | 1.02E-46 | 5 | Cd48 | 0.252 |
| 5.06E-51 | 0.3185697 | 0.781 | 0.482 | 1.57E-46 | 5 | Fnbp1 | 0.299 |
| 5.22E-51 | 0.2643581 | 0.401 | 0.188 | 1.62E-46 | 5 | Stard9 | 0.213 |
| 5.44E-51 | 0.4391212 | 0.785 | 0.525 | 1.69E-46 | 5 | Plekho2 | 0.26 |
| 7.51E-51 | 0.3386775 | 0.914 | 0.633 | 2.33E-46 | 5 | Ly86 | 0.281 |
| 1.01E-50 | 0.2195814 | 0.24 | 0.088 | 3.13E-46 | 5 | C1qtnf1 | 0.152 |
| 1.19E-50 | 0.2772275 | 0.491 | 0.253 | 3.69E-46 | 5 | Camk1 | 0.238 |
| 2.40E-50 | 0.1937546 | 0.112 | 0.026 | 7.44E-46 | 5 | Zmynd15 | 0.086 |
| 3.34E-50 | 0.692709 | 0.921 | 0.739 | 1.04E-45 | 5 | Jun | 0.182 |
| 4.05E-50 | 0.2684098 | 0.527 | 0.278 | 1.26E-45 | 5 | Ctnnd1 | 0.249 |
| 5.26E-50 | 0.2696184 | 0.575 | 0.31 | 1.63E-45 | 5 | Mpp1 | 0.265 |
| 6.98E-50 | 0.2378449 | 0.427 | 0.208 | 2.17E-45 | 5 | Zfp395 | 0.219 |
| 7.56E-50 | 0.4957747 | 0.779 | 0.528 | 2.35E-45 | 5 | Nfkbid | 0.251 |
| 7.64E-50 | 0.3624098 | 0.63 | 0.362 | 2.37E-45 | 5 | Tlr2 | 0.268 |
| 1.02E-49 | 0.2474351 | 0.527 | 0.28 | 3.16E-45 | 5 | Slc12a6 | 0.247 |
| 2.20E-49 | 0.3955436 | 0.569 | 0.325 | 6.84E-45 | 5 | Casp4 | 0.244 |
| 2.35E-49 | 0.2499271 | 0.413 | 0.197 | 7.30E-45 | 5 | Sesn1 | 0.216 |
| 2.74E-49 | 0.3519319 | 0.288 | 0.122 | 8.50E-45 | 5 | Slfn8 | 0.166 |
| 3.81E-49 | 0.245739 | 0.488 | 0.25 | 1.18E-44 | 5 | Slc29a1 | 0.238 |
| 4.42E-49 | 0.3703658 | 0.834 | 0.53 | 1.37E-44 | 5 | Ptpn18 | 0.304 |
| 5.08E-49 | 0.2526274 | 0.413 | 0.2 | 1.58E-44 | 5 | St6gal1 | 0.213 |
| 7.79E-49 | 0.361218 | 0.99 | 0.83 | 2.42E-44 | 5 | Fcer1g | 0.16 |
| 8.53E-49 | 0.2403434 | 0.505 | 0.265 | 2.65E-44 | 5 | Fam208a | 0.24 |
| 1.10E-48 | 0.1942002 | 0.514 | 0.263 | 3.43E-44 | 5 | Hmgn1 | 0.251 |
| 1.15E-48 | 0.2046416 | 0.735 | 0.424 | 3.58E-44 | 5 | Apobec3 | 0.311 |
| 1.23E-48 | 0.3425607 | 0.809 | 0.525 | 3.82E-44 | 5 | Rock2 | 0.284 |
| 1.37E-48 | 0.2660986 | 0.233 | 0.087 | 4.25E-44 | 5 | Paox | 0.146 |
| 1.81E-48 | 0.4908703 | 0.64 | 0.381 | 5.61E-44 | 5 | Ccl24 | 0.259 |
| 1.90E-48 | 0.2937422 | 0.364 | 0.173 | 5.91E-44 | 5 | Tbc1d5 | 0.191 |
| 2.19E-48 | 0.2665029 | 0.468 | 0.244 | 6.79E-44 | 5 | Saraf | 0.224 |
| 2.21E-48 | 0.1794212 | 0.101 | 0.023 | 6.87E-44 | 5 | Ms4a14 | 0.078 |
| 2.50E-48 | 0.2116621 | 0.151 | 0.044 | 7.77E-44 | 5 | Maml2 | 0.107 |

| | | | | | | | |
|---|---|---|---|---|---|---|---|
| 3.08E-48 | 0.2676517 | 0.355 | 0.164 | 9.58E-44 | 5 | Gpr160 | 0.191 |
| 3.99E-48 | 0.2977338 | 0.452 | 0.236 | 1.24E-43 | 5 | Glmp | 0.216 |
| 5.55E-48 | 0.244909 | 0.184 | 0.062 | 1.72E-43 | 5 | Pld2 | 0.122 |
| 1.47E-47 | 0.2158378 | 0.474 | 0.243 | 4.56E-43 | 5 | H2-M3 | 0.231 |
| 2.06E-47 | 0.2428621 | 0.471 | 0.245 | 6.39E-43 | 5 | Mcfd2 | 0.226 |
| 3.34E-47 | 0.3111144 | 0.369 | 0.178 | 1.04E-42 | 5 | Mid1ip1 | 0.191 |
| 5.24E-47 | 0.6366847 | 0.519 | 0.307 | 1.63E-42 | 5 | Stap1 | 0.212 |
| 5.86E-47 | 0.2238021 | 0.472 | 0.243 | 1.82E-42 | 5 | Ypel3 | 0.229 |
| 7.47E-47 | 0.2530988 | 0.273 | 0.114 | 2.32E-42 | 5 | Mink1 | 0.159 |
| 8.45E-47 | 0.2635007 | 0.252 | 0.101 | 2.62E-42 | 5 | Slc16a6 | 0.151 |
| 1.29E-46 | 0.3037708 | 0.761 | 0.478 | 4.00E-42 | 5 | Clec4a3 | 0.283 |
| 2.23E-46 | 0.2571502 | 0.306 | 0.135 | 6.93E-42 | 5 | Arhgap12 | 0.171 |
| 2.53E-46 | 0.2292193 | 0.207 | 0.076 | 7.85E-42 | 5 | Engase | 0.131 |
| 2.58E-46 | 0.2649323 | 0.516 | 0.274 | 8.01E-42 | 5 | Sash1 | 0.242 |
| 3.08E-46 | 0.2781699 | 0.448 | 0.234 | 9.57E-42 | 5 | Dynll2 | 0.214 |
| 3.47E-46 | 0.2430287 | 0.281 | 0.119 | 1.08E-41 | 5 | Fads1 | 0.162 |
| 7.14E-46 | 0.2505275 | 0.246 | 0.099 | 2.22E-41 | 5 | Klhl18 | 0.147 |
| 8.25E-46 | 0.362779 | 0.868 | 0.585 | 2.56E-41 | 5 | Fcgr2b | 0.283 |
| 8.78E-46 | 0.2572032 | 0.443 | 0.229 | 2.73E-41 | 5 | Snx8 | 0.214 |
| 9.15E-46 | 0.4106186 | 0.287 | 0.126 | 2.84E-41 | 5 | Prdm1 | 0.161 |
| 9.38E-46 | 0.3002415 | 0.729 | 0.432 | 2.91E-41 | 5 | Abca1 | 0.297 |
| 1.26E-45 | 0.2766413 | 0.378 | 0.185 | 3.91E-41 | 5 | Usp24 | 0.193 |
| 2.19E-45 | 0.3641971 | 0.702 | 0.433 | 6.79E-41 | 5 | Herpud1 | 0.269 |
| 2.32E-45 | 0.2452629 | 0.491 | 0.266 | 7.19E-41 | 5 | Ttc3 | 0.225 |
| 3.45E-45 | 0.4812664 | 0.979 | 0.86 | 1.07E-40 | 5 | Kctd12 | 0.119 |
| 4.37E-45 | 0.2080368 | 0.214 | 0.079 | 1.36E-40 | 5 | Tcf7l2 | 0.135 |
| 5.25E-45 | 0.1757794 | 0.864 | 0.578 | 1.63E-40 | 5 | Tcf4 | 0.286 |
| 6.65E-45 | 0.2006115 | 0.155 | 0.049 | 2.07E-40 | 5 | Nlrc4 | 0.106 |
| 6.76E-45 | 0.2886489 | 0.675 | 0.401 | 2.10E-40 | 5 | Clec4a2 | 0.274 |
| 1.38E-44 | 0.268952 | 0.446 | 0.233 | 4.27E-40 | 5 | Wdfy2 | 0.213 |
| 2.75E-44 | 0.8737796 | 0.296 | 0.138 | 8.53E-40 | 5 | Cxcl1 | 0.158 |
| 2.76E-44 | 0.3502271 | 0.838 | 0.563 | 8.58E-40 | 5 | Pld4 | 0.275 |
| 5.77E-44 | 0.3123741 | 0.972 | 0.826 | 1.79E-39 | 5 | Ddx5 | 0.146 |
| 6.84E-44 | 0.3482782 | 0.629 | 0.38 | 2.12E-39 | 5 | Neurl3 | 0.249 |
| 8.70E-44 | 0.2703105 | 0.29 | 0.129 | 2.70E-39 | 5 | Nhlrc3 | 0.161 |
| 1.32E-43 | 0.2925974 | 0.946 | 0.74 | 4.09E-39 | 5 | Rtn4 | 0.206 |
| 1.55E-43 | 0.366769 | 0.696 | 0.445 | 4.80E-39 | 5 | Mapkapk2 | 0.251 |
| 1.57E-43 | 0.184149 | 0.196 | 0.071 | 4.86E-39 | 5 | Cbx6 | 0.125 |
| 2.01E-43 | 0.215778 | 0.33 | 0.154 | 6.24E-39 | 5 | Nfatc1 | 0.176 |
| 2.13E-43 | 0.2506588 | 0.534 | 0.3 | 6.62E-39 | 5 | Dnase2a | 0.234 |
| 2.70E-43 | 0.2612433 | 0.367 | 0.182 | 8.38E-39 | 5 | Ulk2 | 0.185 |
| 3.02E-43 | 0.1905488 | 0.105 | 0.026 | 9.38E-39 | 5 | P2ry13 | 0.079 |
| 3.98E-43 | 0.2456074 | 0.409 | 0.21 | 1.24E-38 | 5 | Tnfsf12 | 0.199 |
| 5.33E-43 | 0.1881294 | 0.136 | 0.04 | 1.65E-38 | 5 | Tfec | 0.096 |
| 6.79E-43 | 0.228871 | 0.234 | 0.094 | 2.11E-38 | 5 | Foxred2 | 0.14 |
| 1.01E-42 | 0.2173033 | 0.581 | 0.326 | 3.15E-38 | 5 | Serpinb8 | 0.255 |
| 1.17E-42 | 0.3158975 | 0.39 | 0.199 | 3.63E-38 | 5 | Rab20 | 0.191 |
| 1.46E-42 | 0.2507091 | 0.246 | 0.103 | 4.53E-38 | 5 | Acsl1 | 0.143 |
| 1.51E-42 | 0.2058617 | 0.223 | 0.089 | 4.68E-38 | 5 | Pygb | 0.134 |
| 1.79E-42 | 0.232948 | 0.233 | 0.094 | 5.55E-38 | 5 | Prpsap2 | 0.139 |
| 3.21E-42 | 0.205026 | 0.177 | 0.062 | 9.95E-38 | 5 | Zcchc14 | 0.115 |
| 3.59E-42 | 0.1860366 | 0.122 | 0.034 | 1.12E-37 | 5 | Aspa | 0.088 |
| 3.88E-42 | 0.2096433 | 0.281 | 0.124 | 1.21E-37 | 5 | Exoc6b | 0.157 |
| 7.65E-42 | 0.3689513 | 0.511 | 0.297 | 2.37E-37 | 5 | Zfp703 | 0.214 |
| 9.08E-42 | 0.4608748 | 0.936 | 0.774 | 2.82E-37 | 5 | Btg2 | 0.162 |
| 1.08E-41 | 0.2853697 | 0.794 | 0.523 | 3.36E-37 | 5 | Tmcc1 | 0.271 |

| | | | | | | | |
|---|---|---|---|---|---|---|---|
| 2.85E-41 | 0.2503078 | 0.203 | 0.078 | 8.84E-37 | 5 | Kmo | 0.125 |
| 2.86E-41 | 0.2210774 | 0.49 | 0.27 | 8.88E-37 | 5 | Ugcg | 0.22 |
| 2.92E-41 | 0.2184109 | 0.238 | 0.099 | 9.08E-37 | 5 | Tbc1d9b | 0.139 |
| 3.38E-41 | 0.2347755 | 0.298 | 0.137 | 1.05E-36 | 5 | Epb41 | 0.161 |
| 3.43E-41 | 0.3922418 | 0.636 | 0.407 | 1.07E-36 | 5 | Ets2 | 0.229 |
| 3.46E-41 | 0.221022 | 0.203 | 0.078 | 1.08E-36 | 5 | Xylt2 | 0.125 |
| 3.71E-41 | 0.1655908 | 0.949 | 0.678 | 1.15E-36 | 5 | Timp2 | 0.271 |
| 4.28E-41 | 0.2379482 | 0.651 | 0.394 | 1.33E-36 | 5 | Fcho2 | 0.257 |
| 4.28E-41 | 0.2224804 | 0.276 | 0.122 | 1.33E-36 | 5 | Bcl2l1 | 0.154 |
| 4.71E-41 | 0.2814066 | 0.994 | 0.918 | 1.46E-36 | 5 | H2-K1 | 0.076 |
| 4.99E-41 | 0.2411351 | 0.599 | 0.352 | 1.55E-36 | 5 | Gapvd1 | 0.247 |
| 5.30E-41 | 0.2216281 | 0.167 | 0.058 | 1.65E-36 | 5 | Cmtm4 | 0.109 |
| 5.35E-41 | 0.1964142 | 0.574 | 0.329 | 1.66E-36 | 5 | Leng8 | 0.245 |
| 7.96E-41 | 0.2437242 | 0.524 | 0.298 | 2.47E-36 | 5 | Snx6 | 0.226 |
| 8.50E-41 | 0.3285313 | 0.938 | 0.699 | 2.64E-36 | 5 | Ctsh | 0.239 |
| 9.78E-41 | 0.2501263 | 0.235 | 0.096 | 3.04E-36 | 5 | P2ry12 | 0.139 |
| 1.19E-40 | 0.2092911 | 0.254 | 0.11 | 3.70E-36 | 5 | Mtr | 0.144 |
| 1.24E-40 | 0.2651637 | 0.476 | 0.26 | 3.84E-36 | 5 | Ptgs1 | 0.216 |
| 2.58E-40 | 0.5301426 | 0.807 | 0.588 | 8.02E-36 | 5 | Atf3 | 0.219 |
| 9.67E-40 | 0.2584766 | 0.313 | 0.15 | 3.00E-35 | 5 | Dusp16 | 0.163 |
| 1.18E-39 | 0.4117472 | 0.329 | 0.161 | 3.68E-35 | 5 | Errfi1 | 0.168 |
| 1.42E-39 | 0.2364179 | 1 | 0.997 | 4.42E-35 | 5 | mt-Co3 | 0.003 |
| 1.70E-39 | 0.2293122 | 0.331 | 0.161 | 5.29E-35 | 5 | Nfatc2 | 0.17 |
| 1.77E-39 | 0.3689746 | 0.705 | 0.462 | 5.49E-35 | 5 | Nfe2l2 | 0.243 |
| 2.01E-39 | 0.2526583 | 0.682 | 0.425 | 6.23E-35 | 5 | Arhgap18 | 0.257 |
| 2.15E-39 | 0.2014464 | 0.215 | 0.086 | 6.68E-35 | 5 | Fam219a | 0.129 |
| 2.40E-39 | 0.2344384 | 0.226 | 0.094 | 7.45E-35 | 5 | Zeb2os | 0.132 |
| 3.15E-39 | 0.3050385 | 0.172 | 0.061 | 9.78E-35 | 5 | C6 | 0.111 |
| 3.31E-39 | 0.2122305 | 0.474 | 0.262 | 1.03E-34 | 5 | Dnajc13 | 0.212 |
| 3.54E-39 | 0.4401577 | 0.707 | 0.484 | 1.10E-34 | 5 | Birc3 | 0.223 |
| 5.46E-39 | 0.1957548 | 0.451 | 0.247 | 1.70E-34 | 5 | Hmox2 | 0.204 |
| 6.41E-39 | 0.2312145 | 0.516 | 0.295 | 1.99E-34 | 5 | Washc4 | 0.221 |
| 7.57E-39 | 0.2099768 | 0.259 | 0.114 | 2.35E-34 | 5 | Rnf169 | 0.145 |
| 1.02E-38 | 0.2601223 | 0.648 | 0.393 | 3.18E-34 | 5 | Plxnb2 | 0.255 |
| 1.11E-38 | 0.2313493 | 0.824 | 0.548 | 3.46E-34 | 5 | Rnf130 | 0.276 |
| 1.19E-38 | 0.1983869 | 0.203 | 0.08 | 3.69E-34 | 5 | Zfp90 | 0.123 |
| 1.30E-38 | 0.1618033 | 0.119 | 0.035 | 4.04E-34 | 5 | Tlcd2 | 0.084 |
| 1.39E-38 | 0.1934769 | 0.181 | 0.067 | 4.33E-34 | 5 | Agmo | 0.114 |
| 3.29E-38 | 0.3173672 | 0.487 | 0.281 | 1.02E-33 | 5 | Ciita | 0.206 |
| 4.20E-38 | 0.2740498 | 0.601 | 0.364 | 1.30E-33 | 5 | Il10ra | 0.237 |
| 4.90E-38 | 0.2134279 | 0.258 | 0.114 | 1.52E-33 | 5 | Tspan4 | 0.144 |
| 5.24E-38 | 0.1830067 | 0.204 | 0.081 | 1.63E-33 | 5 | Maged1 | 0.123 |
| 5.58E-38 | 0.259817 | 0.974 | 0.724 | 1.73E-33 | 5 | F13a1 | 0.25 |
| 5.77E-38 | 0.2393503 | 0.715 | 0.452 | 1.79E-33 | 5 | Scamp2 | 0.263 |
| 6.22E-38 | 0.2218077 | 0.263 | 0.119 | 1.93E-33 | 5 | Pcyox1 | 0.144 |
| 6.29E-38 | 0.2219774 | 0.389 | 0.203 | 1.95E-33 | 5 | Dennd1a | 0.186 |
| 7.02E-38 | 0.3613358 | 0.876 | 0.635 | 2.18E-33 | 5 | Calr | 0.241 |
| 8.11E-38 | 0.2299755 | 0.308 | 0.149 | 2.52E-33 | 5 | H2-Q4 | 0.159 |
| 8.34E-38 | 0.4143557 | 0.614 | 0.394 | 2.59E-33 | 5 | Sqstm1 | 0.22 |
| 9.14E-38 | 0.2057397 | 0.255 | 0.114 | 2.84E-33 | 5 | BC037034 | 0.141 |
| 9.24E-38 | 0.2162965 | 0.42 | 0.228 | 2.87E-33 | 5 | Sipa1 | 0.192 |
| 1.00E-37 | 0.1831923 | 0.326 | 0.16 | 3.12E-33 | 5 | Ip6k1 | 0.166 |
| 1.04E-37 | 0.2027631 | 0.456 | 0.253 | 3.22E-33 | 5 | Rbm47 | 0.203 |
| 1.17E-37 | 0.241932 | 0.437 | 0.241 | 3.65E-33 | 5 | Mon2 | 0.196 |
| 1.22E-37 | 0.2246424 | 0.541 | 0.319 | 3.78E-33 | 5 | Pias1 | 0.222 |
| 1.29E-37 | 0.1987683 | 0.255 | 0.113 | 4.01E-33 | 5 | Plscr3 | 0.142 |

| | | | | | | | |
|---|---|---|---|---|---|---|---|
| 1.31E-37 | 0.1716832 | 0.193 | 0.075 | 4.06E-33 | 5 | Abcb1b | 0.118 |
| 1.55E-37 | 0.2656983 | 0.714 | 0.46 | 4.83E-33 | 5 | Jmjd1c | 0.254 |
| 1.87E-37 | 0.2600021 | 0.617 | 0.382 | 5.82E-33 | 5 | Pnisr | 0.235 |
| 1.88E-37 | 0.1696282 | 0.609 | 0.362 | 5.84E-33 | 5 | Gng2 | 0.247 |
| 2.51E-37 | 0.2676531 | 0.415 | 0.227 | 7.80E-33 | 5 | Rapgef6 | 0.188 |
| 3.19E-37 | 0.2852367 | 0.756 | 0.498 | 9.90E-33 | 5 | Vsir | 0.258 |
| 3.34E-37 | 0.2171967 | 0.479 | 0.269 | 1.04E-32 | 5 | Bin1 | 0.21 |
| 4.47E-37 | 0.1907675 | 0.547 | 0.32 | 1.39E-32 | 5 | Ski | 0.227 |
| 5.31E-37 | 0.2176447 | 0.571 | 0.337 | 1.65E-32 | 5 | Lrp6 | 0.234 |
| 6.49E-37 | 0.1921545 | 0.259 | 0.117 | 2.02E-32 | 5 | Tlr4 | 0.142 |
| 6.79E-37 | 0.246067 | 0.363 | 0.192 | 2.11E-32 | 5 | Hs6st1 | 0.171 |
| 6.92E-37 | 0.2998429 | 0.892 | 0.668 | 2.15E-32 | 5 | Picalm | 0.224 |
| 7.24E-37 | 0.3994824 | 0.988 | 0.899 | 2.25E-32 | 5 | H3f3b | 0.089 |
| 7.46E-37 | 0.2299992 | 0.443 | 0.248 | 2.32E-32 | 5 | Sirt2 | 0.195 |
| 7.56E-37 | 0.2486901 | 0.278 | 0.132 | 2.35E-32 | 5 | Vps18 | 0.146 |
| 8.75E-37 | 0.2539876 | 0.452 | 0.255 | 2.72E-32 | 5 | Specc1l | 0.197 |
| 9.18E-37 | 0.1672939 | 0.344 | 0.171 | 2.85E-32 | 5 | Rhoc | 0.173 |
| 1.07E-36 | 0.238084 | 0.686 | 0.432 | 3.33E-32 | 5 | Slc43a2 | 0.254 |
| 1.10E-36 | 0.1746878 | 0.315 | 0.152 | 3.42E-32 | 5 | Snx13 | 0.163 |
| 1.72E-36 | 0.185852 | 0.592 | 0.356 | 5.35E-32 | 5 | Mafg | 0.236 |
| 1.85E-36 | 0.1979074 | 0.392 | 0.209 | 5.76E-32 | 5 | Ifnar2 | 0.183 |
| 1.89E-36 | 0.1951454 | 0.266 | 0.122 | 5.88E-32 | 5 | Dennd1c | 0.144 |
| 1.91E-36 | 0.250961 | 0.766 | 0.497 | 5.92E-32 | 5 | Creg1 | 0.269 |
| 1.96E-36 | 0.2117015 | 0.273 | 0.127 | 6.09E-32 | 5 | Plxnc1 | 0.146 |
| 2.03E-36 | 0.2091545 | 0.63 | 0.389 | 6.29E-32 | 5 | Inpp5d | 0.241 |
| 2.57E-36 | 0.1626047 | 0.234 | 0.1 | 7.97E-32 | 5 | Cbx7 | 0.134 |
| 2.73E-36 | 0.1997202 | 0.351 | 0.18 | 8.48E-32 | 5 | Kcnk6 | 0.171 |
| 2.84E-36 | 0.1854731 | 0.119 | 0.036 | 8.83E-32 | 5 | Trpm2 | 0.083 |
| 6.57E-36 | 0.2102848 | 0.278 | 0.131 | 2.04E-31 | 5 | Hck | 0.147 |
| 7.41E-36 | 0.1677914 | 0.137 | 0.045 | 2.30E-31 | 5 | Fer | 0.092 |
| 7.95E-36 | 0.2472912 | 0.409 | 0.224 | 2.47E-31 | 5 | Hfe | 0.185 |
| 8.90E-36 | 0.2468994 | 0.205 | 0.086 | 2.76E-31 | 5 | Tmem260 | 0.119 |
| 9.16E-36 | 0.2154928 | 0.26 | 0.118 | 2.84E-31 | 5 | Ddx60 | 0.142 |
| 9.26E-36 | 0.1724926 | 0.4 | 0.212 | 2.87E-31 | 5 | 8-Sep | 0.188 |
| 1.04E-35 | 0.1908212 | 0.347 | 0.177 | 3.23E-31 | 5 | Micu1 | 0.17 |
| 1.13E-35 | 0.1752458 | 0.351 | 0.18 | 3.52E-31 | 5 | Clasp2 | 0.171 |
| 1.16E-35 | 0.1934941 | 0.615 | 0.376 | 3.60E-31 | 5 | Scpep1 | 0.239 |
| 1.32E-35 | 0.1565579 | 0.207 | 0.085 | 4.09E-31 | 5 | Pld1 | 0.122 |
| 1.39E-35 | 0.2396513 | 0.602 | 0.37 | 4.31E-31 | 5 | Cndp2 | 0.232 |
| 1.42E-35 | 0.2602314 | 0.647 | 0.406 | 4.41E-31 | 5 | P2rx4 | 0.241 |
| 1.43E-35 | 0.2072813 | 0.351 | 0.181 | 4.44E-31 | 5 | Ogfrl1 | 0.17 |
| 1.68E-35 | 0.1960106 | 0.55 | 0.326 | 5.22E-31 | 5 | Cd164 | 0.224 |
| 1.78E-35 | 0.2319332 | 0.266 | 0.125 | 5.53E-31 | 5 | Pde3b | 0.141 |
| 2.13E-35 | 0.1577008 | 0.176 | 0.067 | 6.62E-31 | 5 | Tef | 0.109 |
| 2.22E-35 | 0.5143928 | 0.847 | 0.661 | 6.90E-31 | 5 | Fosb | 0.186 |
| 2.78E-35 | 0.5233744 | 0.845 | 0.715 | 8.63E-31 | 5 | Nr4a1 | 0.13 |
| 2.83E-35 | 0.2056817 | 0.418 | 0.228 | 8.79E-31 | 5 | Ocrl | 0.19 |
| 3.24E-35 | 0.1234991 | 0.417 | 0.223 | 1.01E-30 | 5 | Arhgef12 | 0.194 |
| 3.25E-35 | 0.2127304 | 0.433 | 0.242 | 1.01E-30 | 5 | Fnip1 | 0.191 |
| 3.66E-35 | 0.210391 | 0.448 | 0.255 | 1.14E-30 | 5 | Ankfy1 | 0.193 |
| 4.21E-35 | 0.1718772 | 0.138 | 0.046 | 1.31E-30 | 5 | Slc12a2 | 0.092 |
| 4.74E-35 | 0.232573 | 0.235 | 0.104 | 1.47E-30 | 5 | Oasl2 | 0.131 |
| 5.58E-35 | 0.2178208 | 0.257 | 0.12 | 1.73E-30 | 5 | Dctn5 | 0.137 |
| 5.77E-35 | 0.2140881 | 0.178 | 0.07 | 1.79E-30 | 5 | Akr1b10 | 0.108 |
| 7.71E-35 | 0.2296026 | 0.27 | 0.13 | 2.40E-30 | 5 | Sft2d1 | 0.14 |
| 8.24E-35 | 0.1917693 | 0.134 | 0.045 | 2.56E-30 | 5 | Vegfb | 0.089 |

| | | | | | | | |
|---|---|---|---|---|---|---|---|
| 9.25E-35 | 0.2141854 | 0.232 | 0.102 | 2.87E-30 | 5 | Csad | 0.13 |
| 1.04E-34 | 0.2843056 | 0.696 | 0.459 | 3.23E-30 | 5 | Cflar | 0.237 |
| 1.36E-34 | 0.223942 | 0.554 | 0.336 | 4.23E-30 | 5 | Pura | 0.218 |
| 1.53E-34 | 0.2550259 | 0.666 | 0.427 | 4.75E-30 | 5 | Sema4a | 0.239 |
| 1.83E-34 | 0.1688244 | 0.148 | 0.052 | 5.67E-30 | 5 | Adamtsl5 | 0.096 |
| 1.97E-34 | 0.2539713 | 0.418 | 0.238 | 6.12E-30 | 5 | Ttyh3 | 0.18 |
| 2.27E-34 | 0.2003555 | 0.443 | 0.249 | 7.05E-30 | 5 | Cln8 | 0.194 |
| 2.39E-34 | 0.2024487 | 0.654 | 0.409 | 7.43E-30 | 5 | Arap1 | 0.245 |
| 2.47E-34 | 0.2885447 | 0.519 | 0.304 | 7.66E-30 | 5 | Ptger4 | 0.215 |
| 2.51E-34 | 0.2206236 | 0.305 | 0.154 | 7.79E-30 | 5 | Acvrl1 | 0.151 |
| 3.41E-34 | 0.2264543 | 0.452 | 0.26 | 1.06E-29 | 5 | Ppp1r21 | 0.192 |
| 3.57E-34 | 0.1917804 | 0.521 | 0.31 | 1.11E-29 | 5 | Prkacb | 0.211 |
| 3.65E-34 | 0.1683732 | 0.603 | 0.368 | 1.13E-29 | 5 | Vwa5a | 0.235 |
| 3.83E-34 | 0.2241525 | 0.373 | 0.202 | 1.19E-29 | 5 | Chd9 | 0.171 |
| 5.20E-34 | 0.2163582 | 0.528 | 0.314 | 1.61E-29 | 5 | Bmp2k | 0.214 |
| 6.17E-34 | 0.2021841 | 0.322 | 0.164 | 1.92E-29 | 5 | Anks1 | 0.158 |
| 6.68E-34 | 0.1911511 | 0.225 | 0.099 | 2.08E-29 | 5 | Pfkfb4 | 0.126 |
| 7.19E-34 | 0.1946409 | 0.136 | 0.046 | 2.23E-29 | 5 | Gm4951 | 0.09 |
| 7.71E-34 | 0.234592 | 0.357 | 0.192 | 2.39E-29 | 5 | Plbd2 | 0.165 |
| 1.04E-33 | 0.2291855 | 0.206 | 0.089 | 3.22E-29 | 5 | Sipa1l1 | 0.117 |
| 1.21E-33 | 0.2110236 | 0.27 | 0.128 | 3.76E-29 | 5 | Gm34084 | 0.142 |
| 1.39E-33 | 0.2337891 | 0.402 | 0.226 | 4.31E-29 | 5 | Os9 | 0.176 |
| 1.41E-33 | 0.2525677 | 0.473 | 0.279 | 4.39E-29 | 5 | Ppp1cc | 0.194 |
| 1.60E-33 | 0.1593667 | 0.139 | 0.048 | 4.98E-29 | 5 | )31425F14R | 0.091 |
| 1.90E-33 | 0.2109734 | 0.223 | 0.099 | 5.91E-29 | 5 | Milr1 | 0.124 |
| 2.00E-33 | 0.1938062 | 0.184 | 0.075 | 6.21E-29 | 5 | Txndc12 | 0.109 |
| 2.03E-33 | 0.1592003 | 0.269 | 0.128 | 6.29E-29 | 5 | Lmbrd2 | 0.141 |
| 2.15E-33 | 0.1668607 | 0.57 | 0.342 | 6.66E-29 | 5 | Foxn3 | 0.228 |
| 2.23E-33 | 0.505672 | 0.26 | 0.129 | 6.92E-29 | 5 | Hsph1 | 0.131 |
| 3.07E-33 | 0.2150632 | 0.307 | 0.157 | 9.54E-29 | 5 | Ifi27 | 0.15 |
| 3.16E-33 | 0.1750148 | 0.495 | 0.29 | 9.82E-29 | 5 | Nfic | 0.205 |
| 3.43E-33 | 0.1284551 | 0.15 | 0.054 | 1.06E-28 | 5 | Fhl3 | 0.096 |
| 3.89E-33 | 0.1441451 | 0.507 | 0.292 | 1.21E-28 | 5 | Srgap2 | 0.215 |
| 4.00E-33 | 0.2410614 | 0.797 | 0.532 | 1.24E-28 | 5 | Dhrs3 | 0.265 |
| 5.79E-33 | 0.2258716 | 0.432 | 0.245 | 1.80E-28 | 5 | Parp14 | 0.187 |
| 6.62E-33 | 0.1593895 | 0.115 | 0.036 | 2.06E-28 | 5 | Hrh1 | 0.079 |
| 7.07E-33 | 0.3653482 | 0.575 | 0.363 | 2.20E-28 | 5 | Irf1 | 0.212 |
| 7.41E-33 | 0.1585876 | 0.54 | 0.329 | 2.30E-28 | 5 | Arl8a | 0.211 |
| 7.57E-33 | 0.2470622 | 0.54 | 0.334 | 2.35E-28 | 5 | Hexb | 0.206 |
| 8.41E-33 | 0.2363072 | 0.295 | 0.146 | 2.61E-28 | 5 | Pdgfb | 0.149 |
| 9.03E-33 | 0.1648564 | 0.194 | 0.08 | 2.81E-28 | 5 | Zdhhc14 | 0.114 |
| 9.58E-33 | 0.2000983 | 0.227 | 0.103 | 2.98E-28 | 5 | 530001G21F | 0.124 |
| 1.01E-32 | 0.1791409 | 0.362 | 0.193 | 3.14E-28 | 5 | Tnfrsf11a | 0.169 |
| 1.07E-32 | 0.1514103 | 0.515 | 0.302 | 3.33E-28 | 5 | Atp6v0a1 | 0.213 |
| 1.27E-32 | 0.1535663 | 0.121 | 0.04 | 3.94E-28 | 5 | Abcc3 | 0.081 |
| 1.40E-32 | 0.1676826 | 0.756 | 0.471 | 4.36E-28 | 5 | Ms4a6c | 0.285 |
| 1.57E-32 | 0.1573161 | 0.179 | 0.072 | 4.87E-28 | 5 | Ampd3 | 0.107 |
| 1.64E-32 | 0.3427595 | 0.974 | 0.915 | 5.08E-28 | 5 | Dusp1 | 0.059 |
| 1.79E-32 | 0.2347366 | 0.489 | 0.292 | 5.56E-28 | 5 | Comt | 0.197 |
| 1.83E-32 | 0.2077714 | 0.525 | 0.316 | 5.69E-28 | 5 | Trim30a | 0.209 |
| 1.93E-32 | 0.1968105 | 0.556 | 0.336 | 6.01E-28 | 5 | )30111J21Ri | 0.22 |
| 2.08E-32 | 0.1800098 | 0.266 | 0.125 | 6.46E-28 | 5 | Rcn3 | 0.141 |
| 2.11E-32 | 0.2194399 | 0.275 | 0.136 | 6.54E-28 | 5 | Glb1 | 0.139 |
| 2.29E-32 | 0.2048838 | 0.829 | 0.575 | 7.10E-28 | 5 | Itm2c | 0.254 |
| 2.43E-32 | 0.2048862 | 0.353 | 0.189 | 7.54E-28 | 5 | Snx30 | 0.164 |
| 2.59E-32 | 0.1807042 | 0.127 | 0.043 | 8.04E-28 | 5 | Per3 | 0.084 |

| | | | | | | | |
|---|---|---|---|---|---|---|---|
| 2.63E-32 | 0.1991332 | 0.842 | 0.582 | 8.18E-28 | 5 | Stk17b | 0.26 |
| 3.30E-32 | 0.1779607 | 0.43 | 0.244 | 1.03E-27 | 5 | Tbc1d14 | 0.186 |
| 3.76E-32 | 0.2087025 | 0.198 | 0.085 | 1.17E-27 | 5 | Zfp512 | 0.113 |
| 4.14E-32 | 0.1625492 | 0.12 | 0.04 | 1.29E-27 | 5 | Fam213a | 0.08 |
| 4.20E-32 | 0.2033582 | 0.531 | 0.323 | 1.30E-27 | 5 | Prex1 | 0.208 |
| 4.21E-32 | 0.6159754 | 0.644 | 0.43 | 1.31E-27 | 5 | Hmox1 | 0.214 |
| 4.69E-32 | 0.1589002 | 0.385 | 0.209 | 1.46E-27 | 5 | Idh2 | 0.176 |
| 5.31E-32 | 0.4366244 | 0.957 | 0.87 | 1.65E-27 | 5 | Fos | 0.087 |
| 5.32E-32 | 0.1408376 | 0.177 | 0.071 | 1.65E-27 | 5 | Cd99l2 | 0.106 |
| 5.33E-32 | 0.1262021 | 0.272 | 0.13 | 1.65E-27 | 5 | Slc28a2 | 0.142 |
| 5.41E-32 | 0.1954499 | 0.231 | 0.106 | 1.68E-27 | 5 | Flcn | 0.125 |
| 5.74E-32 | 0.2435079 | 0.7 | 0.466 | 1.78E-27 | 5 | Top1 | 0.234 |
| 6.07E-32 | 0.180715 | 0.322 | 0.165 | 1.89E-27 | 5 | Alox5 | 0.157 |
| 6.46E-32 | 0.2483829 | 0.421 | 0.245 | 2.01E-27 | 5 | Tank | 0.176 |
| 6.74E-32 | 0.2088938 | 0.318 | 0.166 | 2.09E-27 | 5 | Lyl1 | 0.152 |
| 7.86E-32 | 0.1917313 | 0.213 | 0.095 | 2.44E-27 | 5 | 930402H24R | 0.118 |
| 8.06E-32 | 0.2112307 | 0.492 | 0.296 | 2.50E-27 | 5 | Crlf3 | 0.196 |
| 8.10E-32 | 0.2157175 | 0.803 | 0.545 | 2.51E-27 | 5 | Zcchc6 | 0.258 |
| 8.42E-32 | 0.2403434 | 0.385 | 0.215 | 2.62E-27 | 5 | Trip11 | 0.17 |
| 8.56E-32 | 0.2138263 | 0.327 | 0.173 | 2.66E-27 | 5 | Mctp1 | 0.154 |
| 8.58E-32 | 0.208298 | 0.469 | 0.276 | 2.66E-27 | 5 | Arhgef2 | 0.193 |
| 9.41E-32 | 0.1516097 | 0.191 | 0.08 | 2.92E-27 | 5 | Rubcn | 0.111 |
| 1.02E-31 | 0.1583096 | 0.322 | 0.166 | 3.17E-27 | 5 | Slc23a2 | 0.156 |
| 1.20E-31 | 0.2093873 | 0.84 | 0.598 | 3.73E-27 | 5 | Mat2a | 0.242 |
| 1.43E-31 | 0.1609919 | 0.168 | 0.066 | 4.43E-27 | 5 | Aldh7a1 | 0.102 |
| 1.56E-31 | 0.2986352 | 0.419 | 0.244 | 4.84E-27 | 5 | Slc15a3 | 0.175 |
| 1.82E-31 | 0.1705564 | 0.694 | 0.441 | 5.66E-27 | 5 | Abca9 | 0.253 |
| 2.51E-31 | 0.2127666 | 0.198 | 0.086 | 7.80E-27 | 5 | S1pr2 | 0.112 |
| 2.63E-31 | 0.2182517 | 0.757 | 0.519 | 8.17E-27 | 5 | Ccdc50 | 0.238 |
| 2.74E-31 | 0.1892816 | 0.229 | 0.105 | 8.51E-27 | 5 | Tubgcp5 | 0.124 |
| 3.29E-31 | 0.2071222 | 0.413 | 0.238 | 1.02E-26 | 5 | Aldh9a1 | 0.175 |
| 3.43E-31 | 0.1533939 | 0.214 | 0.095 | 1.07E-26 | 5 | Cpne2 | 0.119 |
| 3.57E-31 | 0.2209612 | 0.432 | 0.253 | 1.11E-26 | 5 | Rap1gds1 | 0.179 |
| 3.63E-31 | 0.2509274 | 0.388 | 0.221 | 1.13E-26 | 5 | Slc35b2 | 0.167 |
| 3.88E-31 | 0.158911 | 0.17 | 0.068 | 1.20E-26 | 5 | Tmem2 | 0.102 |
| 4.28E-31 | 0.2448524 | 0.906 | 0.69 | 1.33E-26 | 5 | Lamp2 | 0.216 |
| 4.31E-31 | 0.2386413 | 0.674 | 0.445 | 1.34E-26 | 5 | Hcls1 | 0.229 |
| 4.40E-31 | 0.2095199 | 1 | 0.97 | 1.37E-26 | 5 | mt-Nd1 | 0.03 |
| 4.66E-31 | 0.2117849 | 0.184 | 0.078 | 1.45E-26 | 5 | Slc11a2 | 0.106 |
| 4.68E-31 | 0.2024506 | 0.411 | 0.238 | 1.45E-26 | 5 | Safb2 | 0.173 |
| 5.39E-31 | 0.1663546 | 0.223 | 0.101 | 1.67E-26 | 5 | Arsk | 0.122 |
| 5.56E-31 | 0.1563327 | 0.475 | 0.277 | 1.73E-26 | 5 | Tmem109 | 0.198 |
| 7.62E-31 | 0.1600395 | 0.373 | 0.203 | 2.36E-26 | 5 | Zfp652 | 0.17 |
| 9.19E-31 | 0.1554962 | 0.147 | 0.055 | 2.85E-26 | 5 | Hs1bp3 | 0.092 |
| 1.05E-30 | 0.1699983 | 0.395 | 0.223 | 3.26E-26 | 5 | Fkbp15 | 0.172 |
| 1.15E-30 | 0.2159517 | 0.407 | 0.23 | 3.58E-26 | 5 | Il4ra | 0.177 |
| 1.26E-30 | 0.1856907 | 0.734 | 0.49 | 3.91E-26 | 5 | Snx2 | 0.244 |
| 1.37E-30 | 0.1343673 | 0.115 | 0.038 | 4.25E-26 | 5 | Pxdc1 | 0.077 |
| 1.43E-30 | 0.2228861 | 0.488 | 0.298 | 4.43E-26 | 5 | Rnpep | 0.19 |
| 1.68E-30 | 0.2313771 | 0.739 | 0.491 | 5.21E-26 | 5 | Pirb | 0.248 |
| 1.97E-30 | 0.1348743 | 0.111 | 0.036 | 6.12E-26 | 5 | Gdpd1 | 0.075 |
| 2.31E-30 | 0.1738371 | 0.18 | 0.076 | 7.18E-26 | 5 | Nagpa | 0.104 |
| 2.80E-30 | 0.2261076 | 0.565 | 0.354 | 8.70E-26 | 5 | Nckap1l | 0.211 |
| 2.81E-30 | 0.2009967 | 0.359 | 0.198 | 8.73E-26 | 5 | Pik3c2a | 0.161 |
| 2.88E-30 | 0.1883569 | 0.21 | 0.094 | 8.95E-26 | 5 | Slc7a7 | 0.116 |
| 2.91E-30 | 0.1673475 | 0.18 | 0.075 | 9.02E-26 | 5 | Tslp | 0.105 |

| | | | | | | | |
|---|---|---|---|---|---|---|---|
| 3.19E-30 | 0.1938123 | 0.408 | 0.236 | 9.91E-26 | 5 | Wbp2 | 0.172 |
| 3.29E-30 | 0.181893 | 0.124 | 0.043 | 1.02E-25 | 5 | Fpr2 | 0.081 |
| 3.37E-30 | 0.2328496 | 0.512 | 0.316 | 1.05E-25 | 5 | Nufip2 | 0.196 |
| 3.56E-30 | 0.1398164 | 0.426 | 0.241 | 1.11E-25 | 5 | Slc7a8 | 0.185 |
| 3.57E-30 | 0.2572166 | 0.92 | 0.68 | 1.11E-25 | 5 | Hexa | 0.24 |
| 3.71E-30 | 0.4481318 | 0.882 | 0.677 | 1.15E-25 | 5 | Mt1 | 0.205 |
| 3.76E-30 | 0.1886479 | 0.65 | 0.418 | 1.17E-25 | 5 | Ptpra | 0.232 |
| 3.95E-30 | 0.1718517 | 0.345 | 0.185 | 1.23E-25 | 5 | Igf1r | 0.16 |
| 4.54E-30 | 0.1776609 | 0.33 | 0.177 | 1.41E-25 | 5 | Gigyf1 | 0.153 |
| 4.63E-30 | 0.1501086 | 0.175 | 0.072 | 1.44E-25 | 5 | Cep68 | 0.103 |
| 4.77E-30 | 0.2062968 | 0.618 | 0.391 | 1.48E-25 | 5 | Tbxas1 | 0.227 |
| 5.55E-30 | 0.1479274 | 0.131 | 0.046 | 1.72E-25 | 5 | Spaca6 | 0.085 |
| 6.01E-30 | 0.155258 | 0.449 | 0.261 | 1.87E-25 | 5 | Etnk1 | 0.188 |
| 6.27E-30 | 0.1908431 | 0.2 | 0.089 | 1.95E-25 | 5 | Acsl3 | 0.111 |
| 6.37E-30 | 0.1822239 | 0.264 | 0.131 | 1.98E-25 | 5 | Enox2 | 0.133 |
| 6.68E-30 | 0.1853875 | 0.503 | 0.305 | 2.07E-25 | 5 | Blnk | 0.198 |
| 6.78E-30 | 0.1868317 | 0.193 | 0.085 | 2.11E-25 | 5 | Nt5c2 | 0.108 |
| 7.05E-30 | 0.1387231 | 0.592 | 0.371 | 2.19E-25 | 5 | Galnt1 | 0.221 |
| 8.87E-30 | 0.2299152 | 0.427 | 0.25 | 2.76E-25 | 5 | Tsc22d2 | 0.177 |
| 9.45E-30 | 0.1737404 | 0.301 | 0.155 | 2.93E-25 | 5 | Phf20 | 0.146 |
| 9.70E-30 | 0.1905568 | 0.173 | 0.072 | 3.01E-25 | 5 | Cd82 | 0.101 |
| 1.05E-29 | 0.2046564 | 0.266 | 0.133 | 3.28E-25 | 5 | Ephx1 | 0.133 |
| 1.20E-29 | 0.1264099 | 0.245 | 0.116 | 3.74E-25 | 5 | Lpar6 | 0.129 |
| 1.26E-29 | 0.2690025 | 0.884 | 0.657 | 3.93E-25 | 5 | Ifitm2 | 0.227 |
| 1.29E-29 | 0.2527658 | 0.245 | 0.118 | 4.00E-25 | 5 | 530028O10R | 0.127 |
| 1.41E-29 | 0.1682241 | 0.122 | 0.043 | 4.36E-25 | 5 | Arid3b | 0.079 |
| 1.45E-29 | 0.1906825 | 0.487 | 0.298 | 4.52E-25 | 5 | Ppp1r12c | 0.189 |
| 1.61E-29 | 0.1792025 | 0.377 | 0.212 | 4.99E-25 | 5 | Phc3 | 0.165 |
| 1.71E-29 | 0.154301 | 0.478 | 0.287 | 5.31E-25 | 5 | Trpc4ap | 0.191 |
| 1.77E-29 | 0.1609351 | 0.261 | 0.128 | 5.50E-25 | 5 | Ly9 | 0.133 |
| 2.04E-29 | 0.1815676 | 0.195 | 0.086 | 6.34E-25 | 5 | Scn1b | 0.109 |
| 2.08E-29 | 0.1841173 | 0.636 | 0.411 | 6.46E-25 | 5 | B4galt6 | 0.225 |
| 2.29E-29 | 0.298934 | 0.16 | 0.065 | 7.10E-25 | 5 | Rrad | 0.095 |
| 2.44E-29 | 0.1806205 | 0.771 | 0.514 | 7.57E-25 | 5 | Mef2a | 0.257 |
| 2.51E-29 | 0.1931053 | 0.525 | 0.326 | 7.78E-25 | 5 | Pten | 0.199 |
| 2.53E-29 | 0.1855865 | 0.315 | 0.168 | 7.85E-25 | 5 | Tmem173 | 0.147 |
| 3.15E-29 | 0.1435585 | 0.474 | 0.283 | 9.77E-25 | 5 | Ifngr2 | 0.191 |
| 3.76E-29 | 0.1579932 | 0.566 | 0.352 | 1.17E-24 | 5 | Susd6 | 0.214 |
| 4.12E-29 | 0.172185 | 0.228 | 0.107 | 1.28E-24 | 5 | Hlx | 0.121 |
| 4.63E-29 | 0.1688779 | 0.58 | 0.367 | 1.44E-24 | 5 | Arid1a | 0.213 |
| 4.90E-29 | 0.1744575 | 0.19 | 0.083 | 1.52E-24 | 5 | Pml | 0.107 |
| 5.07E-29 | 0.1331343 | 0.624 | 0.394 | 1.57E-24 | 5 | Rassf2 | 0.23 |
| 5.46E-29 | 0.1751509 | 0.21 | 0.096 | 1.70E-24 | 5 | Pnpla7 | 0.114 |
| 5.56E-29 | 0.1685487 | 0.215 | 0.099 | 1.73E-24 | 5 | Spire1 | 0.116 |
| 6.01E-29 | 0.1981934 | 0.266 | 0.136 | 1.87E-24 | 5 | Cnpy3 | 0.13 |
| 6.26E-29 | 0.1197487 | 0.119 | 0.041 | 1.94E-24 | 5 | Cers4 | 0.078 |
| 7.16E-29 | 0.1626119 | 0.229 | 0.108 | 2.22E-24 | 5 | Txndc16 | 0.121 |
| 9.11E-29 | 0.1840498 | 0.331 | 0.182 | 2.83E-24 | 5 | Sh2b3 | 0.149 |
| 9.29E-29 | 0.1687068 | 0.35 | 0.193 | 2.88E-24 | 5 | Ago3 | 0.157 |
| 9.29E-29 | 0.4537472 | 0.759 | 0.591 | 2.88E-24 | 5 | Tnfaip3 | 0.168 |
| 1.04E-28 | 0.1545913 | 0.327 | 0.175 | 3.23E-24 | 5 | Galc | 0.152 |
| 1.07E-28 | 0.149949 | 0.556 | 0.347 | 3.31E-24 | 5 | Crebbp | 0.209 |
| 1.12E-28 | 0.1180046 | 0.705 | 0.465 | 3.48E-24 | 5 | Bsg | 0.24 |
| 1.24E-28 | 0.1166445 | 0.483 | 0.288 | 3.85E-24 | 5 | Wls | 0.195 |
| 1.43E-28 | 0.1863331 | 0.139 | 0.053 | 4.43E-24 | 5 | Chst7 | 0.086 |
| 1.56E-28 | 0.1969513 | 0.645 | 0.424 | 4.85E-24 | 5 | Rabac1 | 0.221 |

| | | | | | | | |
|---|---|---|---|---|---|---|---|
| 1.62E-28 | 0.1597198 | 0.277 | 0.142 | 5.03E-24 | 5 | Uhrf1bp1l | 0.135 |
| 1.68E-28 | 0.1985029 | 0.372 | 0.212 | 5.23E-24 | 5 | Dmxl1 | 0.16 |
| 1.75E-28 | 0.1739878 | 0.361 | 0.203 | 5.45E-24 | 5 | Efcab14 | 0.158 |
| 1.76E-28 | 0.1452886 | 0.507 | 0.311 | 5.48E-24 | 5 | Zfand6 | 0.196 |
| 2.13E-28 | 0.1746501 | 0.352 | 0.197 | 6.61E-24 | 5 | Marf1 | 0.155 |
| 2.15E-28 | 0.1767488 | 0.281 | 0.145 | 6.67E-24 | 5 | Armcx3 | 0.136 |
| 2.16E-28 | 0.2042405 | 0.175 | 0.076 | 6.71E-24 | 5 | Tle1 | 0.099 |
| 2.25E-28 | 0.1611224 | 0.162 | 0.067 | 6.98E-24 | 5 | Cib2 | 0.095 |
| 2.40E-28 | 0.1504063 | 0.147 | 0.057 | 7.44E-24 | 5 | Cd79b | 0.09 |
| 2.70E-28 | 0.2184688 | 0.938 | 0.731 | 8.37E-24 | 5 | Sirpa | 0.207 |
| 3.33E-28 | 0.2590158 | 0.867 | 0.647 | 1.03E-23 | 5 | Ddx3x | 0.22 |
| 3.36E-28 | 0.1922592 | 0.303 | 0.161 | 1.04E-23 | 5 | Mfsd11 | 0.142 |
| 3.72E-28 | 0.166131 | 0.305 | 0.162 | 1.15E-23 | 5 | Helz | 0.143 |
| 4.01E-28 | 0.1988042 | 0.418 | 0.247 | 1.24E-23 | 5 | Tnfrsf1a | 0.171 |
| 4.15E-28 | 0.1472971 | 0.754 | 0.508 | 1.29E-23 | 5 | G3bp2 | 0.246 |
| 4.60E-28 | 0.1642424 | 0.278 | 0.144 | 1.43E-23 | 5 | Fbxw4 | 0.134 |
| 5.14E-28 | 0.1925786 | 0.429 | 0.257 | 1.60E-23 | 5 | Twf1 | 0.172 |
| 5.68E-28 | 0.2131651 | 0.567 | 0.365 | 1.76E-23 | 5 | Phip | 0.202 |
| 5.78E-28 | 0.1416833 | 0.262 | 0.13 | 1.79E-23 | 5 | Plekhg5 | 0.132 |
| 6.22E-28 | 0.1627687 | 0.183 | 0.08 | 1.93E-23 | 5 | Fam118a | 0.103 |
| 6.46E-28 | 0.241581 | 0.516 | 0.331 | 2.01E-23 | 5 | Skil | 0.185 |
| 7.00E-28 | 0.1529515 | 0.456 | 0.274 | 2.17E-23 | 5 | Tnrc6b | 0.182 |
| 7.07E-28 | 0.1350018 | 0.321 | 0.174 | 2.20E-23 | 5 | Atraid | 0.147 |
| 7.27E-28 | 0.202736 | 0.28 | 0.149 | 2.26E-23 | 5 | Ahsa1 | 0.131 |
| 7.30E-28 | 0.1612475 | 0.505 | 0.313 | 2.27E-23 | 5 | Snap23 | 0.192 |
| 7.49E-28 | 0.1082104 | 0.101 | 0.033 | 2.32E-23 | 5 | Ikzf2 | 0.068 |
| 7.84E-28 | 0.1306189 | 0.21 | 0.097 | 2.43E-23 | 5 | Grap | 0.113 |
| 8.47E-28 | 0.139361 | 0.219 | 0.104 | 2.63E-23 | 5 | Bmyc | 0.115 |
| 9.40E-28 | 0.161691 | 0.431 | 0.257 | 2.92E-23 | 5 | Fndc3a | 0.174 |
| 9.41E-28 | 0.101988 | 0.69 | 0.463 | 2.92E-23 | 5 | Zbtb20 | 0.227 |
| 9.75E-28 | 0.1334549 | 0.599 | 0.379 | 3.03E-23 | 5 | Dennd5a | 0.22 |
| 9.97E-28 | 0.1656701 | 0.234 | 0.114 | 3.10E-23 | 5 | Fam53b | 0.12 |
| 1.02E-27 | 0.2398051 | 0.355 | 0.204 | 3.15E-23 | 5 | Bcl3 | 0.151 |
| 1.03E-27 | 0.1830217 | 0.274 | 0.145 | 3.19E-23 | 5 | Slc38a10 | 0.129 |
| 1.19E-27 | 0.1378262 | 0.575 | 0.362 | 3.69E-23 | 5 | Ppp1r9b | 0.213 |
| 1.19E-27 | 0.1634474 | 0.203 | 0.094 | 3.70E-23 | 5 | Hk3 | 0.109 |
| 1.35E-27 | 0.1494907 | 0.232 | 0.112 | 4.20E-23 | 5 | Slc16a7 | 0.12 |
| 1.37E-27 | 0.1549811 | 0.329 | 0.182 | 4.25E-23 | 5 | Atp2c1 | 0.147 |
| 1.43E-27 | 0.1688481 | 0.261 | 0.133 | 4.43E-23 | 5 | Parp8 | 0.128 |
| 2.73E-27 | 0.1268312 | 0.148 | 0.059 | 8.47E-23 | 5 | Slc46a3 | 0.089 |
| 3.20E-27 | 0.1534947 | 0.224 | 0.109 | 9.92E-23 | 5 | Tmem205 | 0.115 |
| 3.41E-27 | 0.1251523 | 0.343 | 0.19 | 1.06E-22 | 5 | Rybp | 0.153 |
| 5.07E-27 | 0.1614962 | 0.331 | 0.183 | 1.57E-22 | 5 | Fam234a | 0.148 |
| 5.14E-27 | 0.1621778 | 0.216 | 0.102 | 1.60E-22 | 5 | Slc38a6 | 0.114 |
| 5.30E-27 | 0.2003013 | 0.257 | 0.133 | 1.65E-22 | 5 | Ctbp2 | 0.124 |
| 5.82E-27 | 0.1396442 | 0.461 | 0.278 | 1.81E-22 | 5 | Cmtm7 | 0.183 |
| 6.27E-27 | 0.1692401 | 0.223 | 0.108 | 1.95E-22 | 5 | [illegible]10037D02R | 0.115 |
| 6.31E-27 | 0.1597466 | 0.462 | 0.281 | 1.96E-22 | 5 | Iqgap2 | 0.181 |
| 6.31E-27 | 0.1839862 | 0.517 | 0.325 | 1.96E-22 | 5 | Ash1l | 0.192 |
| 6.53E-27 | 0.212085 | 0.424 | 0.259 | 2.03E-22 | 5 | Ilk | 0.165 |
| 6.66E-27 | 0.141203 | 0.293 | 0.156 | 2.07E-22 | 5 | Rab5b | 0.137 |
| 7.26E-27 | 0.1378328 | 0.252 | 0.127 | 2.25E-22 | 5 | Fez2 | 0.125 |
| 8.65E-27 | 0.1701213 | 0.254 | 0.129 | 2.69E-22 | 5 | Plagl2 | 0.125 |
| 9.20E-27 | 0.1635165 | 0.488 | 0.301 | 2.86E-22 | 5 | Crk | 0.187 |
| 9.75E-27 | 0.2042423 | 0.232 | 0.116 | 3.03E-22 | 5 | Phf11b | 0.116 |
| 1.14E-26 | 0.2216544 | 0.403 | 0.244 | 3.54E-22 | 5 | Mknk2 | 0.159 |

| | | | | | | | |
|---|---|---|---|---|---|---|---|
| 1.25E-26 | 0.1681241 | 0.392 | 0.228 | 3.88E-22 | 5 | Phf21a | 0.164 |
| 1.27E-26 | 0.1651371 | 0.283 | 0.151 | 3.94E-22 | 5 | Zfp950 | 0.132 |
| 1.28E-26 | 0.1462771 | 0.35 | 0.198 | 3.96E-22 | 5 | Pcf11 | 0.152 |
| 1.54E-26 | 0.2017945 | 0.47 | 0.29 | 4.79E-22 | 5 | Rnf19b | 0.18 |
| 1.80E-26 | 0.1706038 | 0.179 | 0.08 | 5.57E-22 | 5 | Fam3a | 0.099 |
| 2.01E-26 | 0.146434 | 0.105 | 0.036 | 6.24E-22 | 5 | Tnfrsf25 | 0.069 |
| 2.07E-26 | 0.1528439 | 0.608 | 0.4 | 6.42E-22 | 5 | Wsb1 | 0.208 |
| 2.34E-26 | 0.1272685 | 0.35 | 0.195 | 7.25E-22 | 5 | Nacc2 | 0.155 |
| 2.44E-26 | 0.1879291 | 0.464 | 0.285 | 7.58E-22 | 5 | Trim8 | 0.179 |
| 2.83E-26 | 0.160788 | 0.167 | 0.073 | 8.80E-22 | 5 | Wdr81 | 0.094 |
| 3.03E-26 | 0.1738301 | 0.513 | 0.327 | 9.41E-22 | 5 | Ttc14 | 0.186 |
| 3.18E-26 | 0.1635801 | 0.312 | 0.17 | 9.89E-22 | 5 | Tmem87b | 0.142 |
| 3.39E-26 | 0.2434135 | 0.919 | 0.726 | 1.05E-21 | 5 | Rsrp1 | 0.193 |
| 3.53E-26 | 0.1512028 | 0.524 | 0.331 | 1.10E-21 | 5 | Mapk14 | 0.193 |
| 3.67E-26 | 0.2646961 | 0.923 | 0.726 | 1.14E-21 | 5 | Hspa5 | 0.197 |
| 3.92E-26 | 0.1667028 | 0.591 | 0.384 | 1.22E-21 | 5 | Cep170 | 0.207 |
| 3.95E-26 | 0.5443571 | 0.706 | 0.509 | 1.23E-21 | 5 | Dnaja1 | 0.197 |
| 4.19E-26 | 0.175186 | 0.207 | 0.099 | 1.30E-21 | 5 | Tmem243 | 0.108 |
| 4.20E-26 | 0.1082782 | 0.18 | 0.08 | 1.31E-21 | 5 | Chpt1 | 0.1 |
| 4.61E-26 | 0.1480986 | 0.602 | 0.388 | 1.43E-21 | 5 | Tet3 | 0.214 |
| 5.06E-26 | 0.2238868 | 0.356 | 0.207 | 1.57E-21 | 5 | Nfkbie | 0.149 |
| 6.01E-26 | 0.1830832 | 0.164 | 0.072 | 1.87E-21 | 5 | Sil1 | 0.092 |
| 6.02E-26 | 0.1545743 | 0.326 | 0.183 | 1.87E-21 | 5 | Gtpbp2 | 0.143 |
| 6.75E-26 | 0.1493315 | 0.609 | 0.397 | 2.10E-21 | 5 | Synj1 | 0.212 |
| 7.14E-26 | 0.1076319 | 0.154 | 0.064 | 2.22E-21 | 5 | Aig1 | 0.09 |
| 7.83E-26 | 0.1771664 | 0.395 | 0.234 | 2.43E-21 | 5 | Dyrk2 | 0.161 |
| 8.12E-26 | 0.1882267 | 0.431 | 0.264 | 2.52E-21 | 5 | Zmynd8 | 0.167 |
| 9.58E-26 | 0.1219882 | 0.22 | 0.107 | 2.97E-21 | 5 | Smpd1 | 0.113 |
| 9.69E-26 | 0.1420279 | 0.202 | 0.094 | 3.01E-21 | 5 | Cracr2b | 0.108 |
| 1.03E-25 | 0.1474092 | 0.364 | 0.211 | 3.19E-21 | 5 | Snx27 | 0.153 |
| 1.18E-25 | 0.1457664 | 0.158 | 0.068 | 3.68E-21 | 5 | Dpp7 | 0.09 |
| 1.19E-25 | 0.1159436 | 0.486 | 0.298 | 3.68E-21 | 5 | Hprt | 0.188 |
| 1.50E-25 | 0.1257027 | 0.166 | 0.072 | 4.66E-21 | 5 | Gkap1 | 0.094 |
| 1.72E-25 | 0.1162791 | 0.42 | 0.252 | 5.35E-21 | 5 | Acat1 | 0.168 |
| 1.87E-25 | 0.130317 | 0.231 | 0.115 | 5.80E-21 | 5 | Pgm2l1 | 0.116 |
| 2.01E-25 | 0.1963477 | 0.395 | 0.236 | 6.25E-21 | 5 | H2-Q7 | 0.159 |
| 2.43E-25 | 0.1331324 | 0.345 | 0.195 | 7.55E-21 | 5 | Atxn7 | 0.15 |
| 2.49E-25 | 0.154816 | 0.331 | 0.188 | 7.72E-21 | 5 | Sav1 | 0.143 |
| 2.53E-25 | 0.1292654 | 0.243 | 0.123 | 7.85E-21 | 5 | Frmd4a | 0.12 |
| 2.71E-25 | 0.1360043 | 0.742 | 0.49 | 8.42E-21 | 5 | Gas7 | 0.252 |
| 3.04E-25 | 0.1406993 | 0.155 | 0.066 | 9.43E-21 | 5 | Sfmbt1 | 0.089 |
| 3.49E-25 | 0.1396991 | 0.147 | 0.061 | 1.08E-20 | 5 | Pgap1 | 0.086 |
| 3.66E-25 | 0.1984536 | 0.231 | 0.117 | 1.14E-20 | 5 | Tnfaip8l2 | 0.114 |
| 3.81E-25 | 0.1664499 | 0.317 | 0.177 | 1.18E-20 | 5 | Ints6l | 0.14 |
| 4.25E-25 | 0.1599374 | 0.629 | 0.415 | 1.32E-20 | 5 | Rnf13 | 0.214 |
| 4.29E-25 | 0.2539999 | 0.697 | 0.48 | 1.33E-20 | 5 | Ehd1 | 0.217 |
| 4.33E-25 | 0.3160832 | 0.399 | 0.243 | 1.35E-20 | 5 | Cd209d | 0.156 |
| 4.59E-25 | 0.1770757 | 0.268 | 0.143 | 1.43E-20 | 5 | Dpf2 | 0.125 |
| 4.71E-25 | 0.1997638 | 0.293 | 0.163 | 1.46E-20 | 5 | Crtc3 | 0.13 |
| 4.79E-25 | 0.1532813 | 0.31 | 0.174 | 1.49E-20 | 5 | Zdhhc9 | 0.136 |
| 5.11E-25 | 0.124873 | 0.232 | 0.116 | 1.59E-20 | 5 | Itpripl1 | 0.116 |
| 5.17E-25 | 0.1463832 | 0.507 | 0.322 | 1.60E-20 | 5 | Psenen | 0.185 |
| 5.85E-25 | 0.1476928 | 0.354 | 0.206 | 1.82E-20 | 5 | Lilrb4a | 0.148 |
| 5.89E-25 | 0.1466661 | 0.661 | 0.436 | 1.83E-20 | 5 | Gnaq | 0.225 |
| 8.88E-25 | 0.1978424 | 0.429 | 0.266 | 2.76E-20 | 5 | Hip1 | 0.163 |
| 9.22E-25 | 0.1564156 | 0.224 | 0.112 | 2.86E-20 | 5 | Myo7a | 0.112 |

| | | | | | | | |
|---|---|---|---|---|---|---|---|
| 1.00E-24 | 0.1449405 | 0.26 | 0.136 | 3.11E-20 | 5 | Atp7a | 0.124 |
| 1.01E-24 | 0.1103739 | 0.207 | 0.098 | 3.12E-20 | 5 | Pbx1 | 0.109 |
| 1.07E-24 | 0.1211029 | 0.128 | 0.05 | 3.33E-20 | 5 | Bcl9 | 0.078 |
| 1.11E-24 | 0.1599356 | 0.509 | 0.325 | 3.46E-20 | 5 | Bptf | 0.184 |
| 1.13E-24 | 0.1639417 | 0.324 | 0.184 | 3.50E-20 | 5 | Gmip | 0.14 |
| 1.13E-24 | 0.162778 | 0.303 | 0.167 | 3.51E-20 | 5 | Tpcn1 | 0.136 |
| 1.15E-24 | 0.1350081 | 0.39 | 0.229 | 3.56E-20 | 5 | Rreb1 | 0.161 |
| 1.25E-24 | 0.1780523 | 0.551 | 0.355 | 3.89E-20 | 5 | Rell1 | 0.196 |
| 1.91E-24 | 0.1643857 | 0.187 | 0.088 | 5.94E-20 | 5 | Dusp7 | 0.099 |
| 2.11E-24 | 0.2076329 | 0.947 | 0.763 | 6.55E-20 | 5 | Irf2bp2 | 0.184 |
| 2.14E-24 | 0.1484925 | 0.241 | 0.125 | 6.63E-20 | 5 | Foxo3 | 0.116 |
| 2.14E-24 | 0.1350875 | 0.406 | 0.244 | 6.65E-20 | 5 | 1-Mar | 0.162 |
| 2.16E-24 | 0.1387934 | 0.31 | 0.174 | 6.71E-20 | 5 | Smarca2 | 0.136 |
| 2.23E-24 | 0.1909171 | 0.406 | 0.25 | 6.92E-20 | 5 | Nucb1 | 0.156 |
| 2.35E-24 | 0.1856152 | 0.271 | 0.147 | 7.30E-20 | 5 | Slc31a2 | 0.124 |
| 3.26E-24 | 0.1482214 | 0.262 | 0.14 | 1.01E-19 | 5 | Tpst2 | 0.122 |
| 4.17E-24 | 0.1463696 | 0.253 | 0.134 | 1.29E-19 | 5 | Cpq | 0.119 |
| 4.74E-24 | 0.2258541 | 0.589 | 0.391 | 1.47E-19 | 5 | Slamf9 | 0.198 |
| 5.18E-24 | 0.1640436 | 0.291 | 0.163 | 1.61E-19 | 5 | Gripap1 | 0.128 |
| 5.32E-24 | 0.1074032 | 0.553 | 0.358 | 1.65E-19 | 5 | Rp2 | 0.195 |
| 6.39E-24 | 0.1493226 | 0.153 | 0.066 | 1.99E-19 | 5 | Lrba | 0.087 |
| 6.42E-24 | 0.1307515 | 0.248 | 0.129 | 1.99E-19 | 5 | Rnf2 | 0.119 |
| 7.18E-24 | 0.1791831 | 0.675 | 0.456 | 2.23E-19 | 5 | Ptpre | 0.219 |
| 7.50E-24 | 0.1464992 | 0.557 | 0.362 | 2.33E-19 | 5 | Ptpn6 | 0.195 |
| 8.11E-24 | 0.1346127 | 0.403 | 0.247 | 2.52E-19 | 5 | Srsf7 | 0.156 |
| 8.92E-24 | 0.1481745 | 0.366 | 0.219 | 2.77E-19 | 5 | )32438A13R | 0.147 |
| 1.02E-23 | 0.1526048 | 0.231 | 0.118 | 3.17E-19 | 5 | Ago4 | 0.113 |
| 1.02E-23 | 0.1544573 | 0.395 | 0.241 | 3.17E-19 | 5 | Pdxk | 0.154 |
| 1.14E-23 | 0.1268224 | 0.109 | 0.041 | 3.54E-19 | 5 | Iqsec2 | 0.068 |
| 1.17E-23 | 0.1770465 | 0.234 | 0.123 | 3.62E-19 | 5 | Ccm2 | 0.111 |
| 1.17E-23 | 0.2263338 | 0.475 | 0.299 | 3.63E-19 | 5 | Clec4b1 | 0.176 |
| 1.62E-23 | 0.1334155 | 0.681 | 0.461 | 5.04E-19 | 5 | Scp2 | 0.22 |
| 1.91E-23 | 0.1396985 | 0.224 | 0.115 | 5.93E-19 | 5 | Fbrs | 0.109 |
| 2.17E-23 | 0.1099959 | 0.289 | 0.16 | 6.75E-19 | 5 | Ncoa2 | 0.129 |
| 2.35E-23 | 0.1018853 | 0.564 | 0.365 | 7.31E-19 | 5 | Epn1 | 0.199 |
| 2.37E-23 | 0.1439683 | 0.465 | 0.294 | 7.36E-19 | 5 | Atf6 | 0.171 |
| 2.98E-23 | 0.1778344 | 0.159 | 0.072 | 9.26E-19 | 5 | Gm46224 | 0.087 |
| 3.09E-23 | 0.1052003 | 0.541 | 0.35 | 9.61E-19 | 5 | Sdf4 | 0.191 |
| 3.23E-23 | 0.1686422 | 0.335 | 0.197 | 1.00E-18 | 5 | Cebpa | 0.138 |
| 3.38E-23 | 0.1691652 | 0.384 | 0.236 | 1.05E-18 | 5 | Ncstn | 0.148 |
| 3.55E-23 | 0.1305707 | 0.223 | 0.114 | 1.10E-18 | 5 | Zscan26 | 0.109 |
| 3.66E-23 | 0.1348952 | 0.244 | 0.129 | 1.14E-18 | 5 | Slc25a45 | 0.115 |
| 3.83E-23 | 0.131255 | 0.451 | 0.285 | 1.19E-18 | 5 | Pkn1 | 0.166 |
| 4.59E-23 | 0.2113755 | 0.489 | 0.319 | 1.43E-18 | 5 | Ppib | 0.17 |
| 4.62E-23 | 0.124731 | 0.516 | 0.33 | 1.43E-18 | 5 | Mxd4 | 0.186 |
| 5.95E-23 | 0.1362407 | 0.562 | 0.37 | 1.85E-18 | 5 | Atp6v1a | 0.192 |
| 6.10E-23 | 0.1164014 | 0.353 | 0.21 | 1.89E-18 | 5 | Slc25a36 | 0.143 |
| 7.81E-23 | 0.2618432 | 0.347 | 0.21 | 2.43E-18 | 5 | Gm6377 | 0.137 |
| 8.33E-23 | 0.1162518 | 0.189 | 0.09 | 2.59E-18 | 5 | Gab1 | 0.099 |
| 8.39E-23 | 0.1359858 | 0.516 | 0.333 | 2.61E-18 | 5 | Fuca1 | 0.183 |
| 1.01E-22 | 0.2372253 | 0.728 | 0.524 | 3.13E-18 | 5 | Tob2 | 0.204 |
| 1.13E-22 | 0.124513 | 0.579 | 0.383 | 3.51E-18 | 5 | Cmtm6 | 0.196 |
| 1.14E-22 | 0.1402751 | 0.29 | 0.165 | 3.55E-18 | 5 | Chordc1 | 0.125 |
| 1.19E-22 | 0.1538537 | 0.246 | 0.131 | 3.70E-18 | 5 | Nbr1 | 0.115 |
| 1.20E-22 | 0.151047 | 0.205 | 0.103 | 3.71E-18 | 5 | Ticam2 | 0.102 |
| 1.27E-22 | 0.1198605 | 0.298 | 0.168 | 3.96E-18 | 5 | Dnase1l1 | 0.13 |

| | | | | | | | |
|---|---|---|---|---|---|---|---|
| 1.52E-22 | 0.1882133 | 0.784 | 0.551 | 4.73E-18 | 5 | Cyth4 | 0.233 |
| 1.66E-22 | 0.1283875 | 0.307 | 0.175 | 5.16E-18 | 5 | Heatr5a | 0.132 |
| 1.73E-22 | 0.1244346 | 0.154 | 0.07 | 5.38E-18 | 5 | Tmem214 | 0.084 |
| 1.83E-22 | 0.1617042 | 0.181 | 0.087 | 5.67E-18 | 5 | Fblim1 | 0.094 |
| 1.95E-22 | 0.105047 | 0.545 | 0.354 | 6.06E-18 | 5 | Rnf141 | 0.191 |
| 2.05E-22 | 0.1125739 | 0.115 | 0.045 | 6.35E-18 | 5 | Rusc2 | 0.07 |
| 2.09E-22 | 0.1657052 | 0.194 | 0.096 | 6.48E-18 | 5 | Kcnk13 | 0.098 |
| 2.13E-22 | 0.1331988 | 0.1 | 0.037 | 6.61E-18 | 5 | Vps37c | 0.063 |
| 2.26E-22 | 0.1465639 | 0.216 | 0.111 | 7.02E-18 | 5 | Hist3h2a | 0.105 |
| 2.35E-22 | 0.1477944 | 0.313 | 0.183 | 7.30E-18 | 5 | Taf6l | 0.13 |
| 2.46E-22 | 0.15793 | 0.505 | 0.326 | 7.63E-18 | 5 | Ifi203 | 0.179 |
| 2.51E-22 | 0.1010532 | 0.53 | 0.349 | 7.80E-18 | 5 | Tnrc18 | 0.181 |
| 2.78E-22 | 0.1395482 | 0.202 | 0.102 | 8.63E-18 | 5 | Abl1 | 0.1 |
| 2.96E-22 | 0.1388584 | 0.143 | 0.063 | 9.20E-18 | 5 | Dcxr | 0.08 |
| 2.98E-22 | 0.1396552 | 0.459 | 0.295 | 9.26E-18 | 5 | Slc29a3 | 0.164 |
| 3.38E-22 | 0.1673843 | 0.461 | 0.297 | 1.05E-17 | 5 | Atxn2l | 0.164 |
| 4.06E-22 | 0.1660298 | 0.23 | 0.122 | 1.26E-17 | 5 | Pak1 | 0.108 |
| 4.32E-22 | 0.1355617 | 0.399 | 0.247 | 1.34E-17 | 5 | Zfp710 | 0.152 |
| 4.49E-22 | 0.1066888 | 0.112 | 0.044 | 1.40E-17 | 5 | Hnmt | 0.068 |
| 4.56E-22 | 0.1424546 | 0.236 | 0.125 | 1.42E-17 | 5 | Mast3 | 0.111 |
| 5.91E-22 | 0.1052995 | 0.102 | 0.038 | 1.84E-17 | 5 | Fam110b | 0.064 |
| 6.26E-22 | 0.1154414 | 0.578 | 0.379 | 1.94E-17 | 5 | Notch2 | 0.199 |
| 6.87E-22 | 0.1139837 | 0.558 | 0.362 | 2.13E-17 | 5 | Ssh2 | 0.196 |
| 7.09E-22 | 0.1556252 | 0.326 | 0.193 | 2.20E-17 | 5 | 931406C07R | 0.133 |
| 7.48E-22 | 0.1284671 | 0.45 | 0.29 | 2.32E-17 | 5 | Sumo1 | 0.16 |
| 7.94E-22 | 0.1146577 | 0.162 | 0.075 | 2.47E-17 | 5 | Mpv17 | 0.087 |
| 8.11E-22 | 0.1252374 | 0.522 | 0.345 | 2.52E-17 | 5 | Per1 | 0.177 |
| 8.22E-22 | 0.149417 | 0.373 | 0.223 | 2.55E-17 | 5 | Dusp6 | 0.15 |
| 9.03E-22 | 0.1352559 | 0.286 | 0.162 | 2.80E-17 | 5 | St6galnac4 | 0.124 |
| 9.44E-22 | 0.2223489 | 0.785 | 0.551 | 2.93E-17 | 5 | Ifi27l2a | 0.234 |
| 1.05E-21 | 0.111076 | 0.103 | 0.039 | 3.26E-17 | 5 | Peli2 | 0.064 |
| 1.06E-21 | 0.3882853 | 0.87 | 0.72 | 3.30E-17 | 5 | Txnip | 0.15 |
| 1.19E-21 | 0.1661479 | 0.309 | 0.18 | 3.69E-17 | 5 | Irf9 | 0.129 |
| 1.22E-21 | 0.1150096 | 0.23 | 0.121 | 3.80E-17 | 5 | Gm37494 | 0.109 |
| 1.27E-21 | 0.1712858 | 1 | 0.981 | 3.96E-17 | 5 | Malat1 | 0.019 |
| 1.31E-21 | 0.1673909 | 0.476 | 0.308 | 4.06E-17 | 5 | Fgd4 | 0.168 |
| 1.31E-21 | 0.1350901 | 0.183 | 0.089 | 4.06E-17 | 5 | Selenbp1 | 0.094 |
| 1.53E-21 | 0.1009785 | 0.153 | 0.069 | 4.75E-17 | 5 | Dgkh | 0.084 |
| 1.56E-21 | 0.1536622 | 0.407 | 0.257 | 4.83E-17 | 5 | mt-Atp8 | 0.15 |
| 1.60E-21 | 0.11363 | 0.203 | 0.103 | 4.96E-17 | 5 | Tmem106c | 0.1 |
| 1.64E-21 | 0.1058714 | 0.655 | 0.436 | 5.11E-17 | 5 | Rab5c | 0.219 |
| 1.76E-21 | 0.1253425 | 0.663 | 0.448 | 5.47E-17 | 5 | Lyn | 0.215 |
| 1.84E-21 | 0.1060029 | 0.549 | 0.362 | 5.70E-17 | 5 | Git2 | 0.187 |
| 1.99E-21 | 0.1674633 | 0.732 | 0.516 | 6.19E-17 | 5 | Cbl | 0.216 |
| 2.20E-21 | 0.1037118 | 0.498 | 0.323 | 6.82E-17 | 5 | Acin1 | 0.175 |
| 2.33E-21 | 0.107717 | 0.218 | 0.113 | 7.23E-17 | 5 | Sh3gl1 | 0.105 |
| 2.43E-21 | 0.1126477 | 0.209 | 0.107 | 7.54E-17 | 5 | Gm36161 | 0.102 |
| 2.48E-21 | 0.1007382 | 0.166 | 0.078 | 7.71E-17 | 5 | Rrm2b | 0.088 |
| 3.61E-21 | 0.1111826 | 0.296 | 0.169 | 1.12E-16 | 5 | Insr | 0.127 |
| 3.69E-21 | 0.1264735 | 0.106 | 0.041 | 1.15E-16 | 5 | Timd4 | 0.065 |
| 4.35E-21 | 0.2973535 | 0.92 | 0.784 | 1.35E-16 | 5 | Nfkbia | 0.136 |
| 4.35E-21 | 0.101308 | 0.129 | 0.054 | 1.35E-16 | 5 | Kif3a | 0.075 |
| 4.36E-21 | 0.1423975 | 0.223 | 0.118 | 1.35E-16 | 5 | Igsf9 | 0.105 |
| 4.48E-21 | 0.1255452 | 0.359 | 0.219 | 1.39E-16 | 5 | Pabpn1 | 0.14 |
| 4.77E-21 | 0.1343763 | 0.286 | 0.165 | 1.48E-16 | 5 | Zkscan3 | 0.121 |
| 5.27E-21 | 0.1492712 | 0.2 | 0.104 | 1.64E-16 | 5 | Pilrb1 | 0.096 |

| | | | | | | | |
|---|---|---|---|---|---|---|---|
| 5.28E-21 | 0.1115357 | 0.286 | 0.165 | 1.64E-16 | 5 | Rbfa | 0.121 |
| 6.31E-21 | 0.1648288 | 0.221 | 0.117 | 1.96E-16 | 5 | Dnajb4 | 0.104 |
| 6.43E-21 | 0.1364611 | 0.152 | 0.07 | 2.00E-16 | 5 | Slc38a7 | 0.082 |
| 6.77E-21 | 0.1247517 | 0.283 | 0.161 | 2.10E-16 | 5 | P4ha1 | 0.122 |
| 7.14E-21 | 0.1851605 | 0.206 | 0.109 | 2.22E-16 | 5 | Tmem229b | 0.097 |
| 7.18E-21 | 0.1508523 | 0.136 | 0.059 | 2.23E-16 | 5 | Kank2 | 0.077 |
| 7.21E-21 | 0.2277722 | 0.912 | 0.69 | 2.24E-16 | 5 | Rrbp1 | 0.222 |
| 1.16E-20 | 0.1108927 | 0.595 | 0.403 | 3.60E-16 | 5 | Mfsd1 | 0.192 |
| 1.23E-20 | 0.1212042 | 0.262 | 0.148 | 3.83E-16 | 5 | Dhrs1 | 0.114 |
| 1.24E-20 | 0.1116534 | 0.505 | 0.333 | 3.85E-16 | 5 | Sumo3 | 0.172 |
| 1.28E-20 | 0.125214 | 0.347 | 0.212 | 3.97E-16 | 5 | Lman1 | 0.135 |
| 1.37E-20 | 0.127267 | 0.446 | 0.288 | 4.25E-16 | 5 | Cmtm3 | 0.158 |
| 1.37E-20 | 0.1198672 | 0.51 | 0.334 | 4.25E-16 | 5 | Camk1d | 0.176 |
| 1.50E-20 | 0.115433 | 0.406 | 0.254 | 4.65E-16 | 5 | Fmnl1 | 0.152 |
| 1.59E-20 | 0.1322497 | 0.183 | 0.091 | 4.94E-16 | 5 | Rnf167 | 0.092 |
| 1.76E-20 | 0.1228807 | 0.223 | 0.119 | 5.45E-16 | 5 | Mtmr12 | 0.104 |
| 1.77E-20 | 0.1432088 | 0.21 | 0.109 | 5.51E-16 | 5 | Tgfbrap1 | 0.101 |
| 1.93E-20 | 0.284548 | 0.464 | 0.313 | 5.99E-16 | 5 | Egr2 | 0.151 |
| 2.38E-20 | 0.1139713 | 0.288 | 0.167 | 7.40E-16 | 5 | Rgl2 | 0.121 |
| 2.39E-20 | 0.1596941 | 0.155 | 0.073 | 7.43E-16 | 5 | Rtp4 | 0.082 |
| 2.49E-20 | 0.1482969 | 0.734 | 0.504 | 7.73E-16 | 5 | Tgfb1 | 0.23 |
| 2.55E-20 | 0.2920719 | 0.281 | 0.167 | 7.91E-16 | 5 | Crem | 0.114 |
| 2.66E-20 | 0.1345136 | 0.236 | 0.129 | 8.26E-16 | 5 | Cdip1 | 0.107 |
| 2.72E-20 | 0.1224288 | 0.332 | 0.2 | 8.43E-16 | 5 | Adam9 | 0.132 |
| 2.81E-20 | 0.1202852 | 0.297 | 0.174 | 8.72E-16 | 5 | Aff1 | 0.123 |
| 2.97E-20 | 0.1444101 | 0.559 | 0.378 | 9.21E-16 | 5 | Ppp1r15a | 0.181 |
| 3.05E-20 | 0.1326673 | 0.548 | 0.364 | 9.47E-16 | 5 | Fermt3 | 0.184 |
| 3.29E-20 | 0.1218914 | 0.295 | 0.173 | 1.02E-15 | 5 | Usp48 | 0.122 |
| 3.31E-20 | 0.1100086 | 0.593 | 0.403 | 1.03E-15 | 5 | Dpysl2 | 0.19 |
| 3.54E-20 | 0.1363278 | 0.204 | 0.107 | 1.10E-15 | 5 | Tor3a | 0.097 |
| 3.56E-20 | 0.1098045 | 0.294 | 0.172 | 1.10E-15 | 5 | Tomm34 | 0.122 |
| 3.90E-20 | 0.1477677 | 0.225 | 0.122 | 1.21E-15 | 5 | Ethe1 | 0.103 |
| 4.17E-20 | 0.1310714 | 0.186 | 0.094 | 1.29E-15 | 5 | Nmd3 | 0.092 |
| 4.17E-20 | 0.1383521 | 0.312 | 0.185 | 1.30E-15 | 5 | Rnf213 | 0.127 |
| 4.71E-20 | 0.1587132 | 0.248 | 0.141 | 1.46E-15 | 5 | Coro7 | 0.107 |
| 5.01E-20 | 0.1211943 | 0.318 | 0.191 | 1.56E-15 | 5 | Ccdc47 | 0.127 |
| 5.22E-20 | 0.1236058 | 0.277 | 0.16 | 1.62E-15 | 5 | Ncoa1 | 0.117 |
| 5.53E-20 | 0.1224593 | 0.422 | 0.27 | 1.72E-15 | 5 | Ik | 0.152 |
| 5.73E-20 | 0.1401995 | 0.289 | 0.171 | 1.78E-15 | 5 | Rnf114 | 0.118 |
| 6.35E-20 | 0.1370851 | 0.185 | 0.094 | 1.97E-15 | 5 | Rasgrp2 | 0.091 |
| 8.34E-20 | 0.1476333 | 0.757 | 0.518 | 2.59E-15 | 5 | Rassf4 | 0.239 |
| 1.53E-19 | 0.1013878 | 0.763 | 0.546 | 4.77E-15 | 5 | Snx3 | 0.217 |
| 1.59E-19 | 0.1362795 | 0.415 | 0.269 | 4.92E-15 | 5 | Mrpl23 | 0.146 |
| 1.64E-19 | 0.1074601 | 0.383 | 0.241 | 5.08E-15 | 5 | Papd4 | 0.142 |
| 1.68E-19 | 0.1005239 | 0.388 | 0.244 | 5.22E-15 | 5 | Cpne3 | 0.144 |
| 1.76E-19 | 0.1101043 | 0.219 | 0.119 | 5.48E-15 | 5 | Ppfibp2 | 0.1 |
| 1.78E-19 | 0.1357181 | 0.488 | 0.322 | 5.52E-15 | 5 | Ost4 | 0.166 |
| 1.89E-19 | 0.1561334 | 0.31 | 0.186 | 5.86E-15 | 5 | Arhgef3 | 0.124 |
| 2.16E-19 | 0.1057507 | 0.14 | 0.063 | 6.70E-15 | 5 | Ttc28 | 0.077 |
| 2.34E-19 | 0.129725 | 0.382 | 0.24 | 7.25E-15 | 5 | Trafd1 | 0.142 |
| 2.46E-19 | 0.1097105 | 0.39 | 0.249 | 7.63E-15 | 5 | Rragc | 0.141 |
| 2.91E-19 | 0.204428 | 0.256 | 0.15 | 9.03E-15 | 5 | Polr2l | 0.106 |
| 2.95E-19 | 0.1164316 | 0.2 | 0.105 | 9.15E-15 | 5 | Trpv2 | 0.095 |
| 2.97E-19 | 0.1231506 | 0.265 | 0.154 | 9.22E-15 | 5 | Vrk2 | 0.111 |
| 3.11E-19 | 0.1311045 | 0.181 | 0.093 | 9.66E-15 | 5 | Wdr11 | 0.088 |
| 3.21E-19 | 0.1466863 | 0.236 | 0.133 | 9.98E-15 | 5 | Kdm5c | 0.103 |

| | | | | | | | |
|---|---|---|---|---|---|---|---|
| 3.31E-19 | 0.1338764 | 0.138 | 0.063 | 1.03E-14 | 5 | Rmi1 | 0.075 |
| 3.36E-19 | 0.1471825 | 0.303 | 0.183 | 1.04E-14 | 5 | Rps6ka1 | 0.12 |
| 3.57E-19 | 0.1057928 | 0.211 | 0.112 | 1.11E-14 | 5 | Ctso | 0.099 |
| 3.73E-19 | 0.1070797 | 0.263 | 0.15 | 1.16E-14 | 5 | Rcsd1 | 0.113 |
| 3.78E-19 | 0.1337532 | 0.411 | 0.265 | 1.17E-14 | 5 | Arid4a | 0.146 |
| 3.87E-19 | 0.1165934 | 0.12 | 0.052 | 1.20E-14 | 5 | Usp21 | 0.068 |
| 3.88E-19 | 0.1328565 | 0.207 | 0.111 | 1.21E-14 | 5 | Tk2 | 0.096 |
| 3.92E-19 | 0.1268868 | 0.14 | 0.064 | 1.22E-14 | 5 | Pdgfa | 0.076 |
| 4.44E-19 | 0.1453899 | 0.995 | 0.967 | 1.38E-14 | 5 | Jund | 0.028 |
| 4.45E-19 | 0.1004867 | 0.274 | 0.16 | 1.38E-14 | 5 | Dync1li2 | 0.114 |
| 4.53E-19 | 0.1862858 | 0.209 | 0.114 | 1.41E-14 | 5 | Mfap3 | 0.095 |
| 4.73E-19 | 0.1007333 | 0.368 | 0.233 | 1.47E-14 | 5 | Snw1 | 0.135 |
| 4.85E-19 | 0.1052555 | 0.707 | 0.491 | 1.50E-14 | 5 | Rab1a | 0.216 |
| 5.16E-19 | 0.1369045 | 0.11 | 0.047 | 1.60E-14 | 5 | Cep85 | 0.063 |
| 5.57E-19 | 0.1267769 | 0.142 | 0.066 | 1.73E-14 | 5 | Slc25a10 | 0.076 |
| 6.01E-19 | 0.1531197 | 0.167 | 0.084 | 1.87E-14 | 5 | Adcy9 | 0.083 |
| 6.12E-19 | 0.1148044 | 0.15 | 0.071 | 1.90E-14 | 5 | Tbc1d31 | 0.079 |
| 6.46E-19 | 0.1099276 | 0.287 | 0.17 | 2.01E-14 | 5 | Tbc1d23 | 0.117 |
| 6.51E-19 | 0.1073092 | 0.65 | 0.443 | 2.02E-14 | 5 | Raph1 | 0.207 |
| 6.51E-19 | 0.1823301 | 0.968 | 0.818 | 2.02E-14 | 5 | Mbnl1 | 0.15 |
| 6.61E-19 | 0.1076667 | 0.244 | 0.137 | 2.05E-14 | 5 | Pbx3 | 0.107 |
| 7.13E-19 | 0.1652014 | 0.221 | 0.122 | 2.21E-14 | 5 | Cbx4 | 0.099 |
| 7.60E-19 | 0.146812 | 0.384 | 0.249 | 2.36E-14 | 5 | Kmt2c | 0.135 |
| 1.04E-18 | 0.1127098 | 0.778 | 0.568 | 3.23E-14 | 5 | Ogt | 0.21 |
| 1.05E-18 | 0.1541322 | 0.833 | 0.625 | 3.27E-14 | 5 | Nrros | 0.208 |
| 1.05E-18 | 0.1369622 | 0.142 | 0.067 | 3.27E-14 | 5 | Map3k3 | 0.075 |
| 1.12E-18 | 0.1020555 | 0.486 | 0.324 | 3.47E-14 | 5 | Stx4a | 0.162 |
| 1.19E-18 | 0.1369726 | 0.332 | 0.206 | 3.68E-14 | 5 | Stx16 | 0.126 |
| 1.25E-18 | 0.105581 | 0.387 | 0.248 | 3.88E-14 | 5 | Zfp148 | 0.139 |
| 1.33E-18 | 0.1428332 | 0.363 | 0.232 | 4.12E-14 | 5 | Crlf2 | 0.131 |
| 1.38E-18 | 0.1218478 | 0.127 | 0.057 | 4.28E-14 | 5 | Ccdc134 | 0.07 |
| 1.47E-18 | 0.1450435 | 0.181 | 0.095 | 4.55E-14 | 5 | Icosl | 0.086 |
| 1.49E-18 | 0.1385677 | 0.414 | 0.272 | 4.64E-14 | 5 | Mvp | 0.142 |
| 1.76E-18 | 0.1266319 | 0.193 | 0.103 | 5.47E-14 | 5 | Mgat4a | 0.09 |
| 1.88E-18 | 0.1175228 | 0.411 | 0.267 | 5.83E-14 | 5 | Washc2 | 0.144 |
| 1.89E-18 | 0.1021922 | 0.143 | 0.067 | 5.86E-14 | 5 | Uba7 | 0.076 |
| 2.11E-18 | 0.1168441 | 0.351 | 0.223 | 6.54E-14 | 5 | Cyb5a | 0.128 |
| 2.13E-18 | 0.1026433 | 0.296 | 0.178 | 6.61E-14 | 5 | Xpo6 | 0.118 |
| 2.19E-18 | 0.1041388 | 0.854 | 0.633 | 6.79E-14 | 5 | Qk | 0.221 |
| 2.50E-18 | 0.1189733 | 0.307 | 0.186 | 7.76E-14 | 5 | D1Ertd622e | 0.121 |
| 2.59E-18 | 0.2029029 | 0.383 | 0.245 | 8.04E-14 | 5 | Mt2 | 0.138 |
| 2.70E-18 | 0.1324705 | 0.172 | 0.088 | 8.38E-14 | 5 | Erf | 0.084 |
| 2.96E-18 | 0.1068237 | 0.592 | 0.408 | 9.20E-14 | 5 | Tmed7 | 0.184 |
| 3.07E-18 | 0.1523566 | 0.352 | 0.225 | 9.53E-14 | 5 | Impact | 0.127 |
| 3.29E-18 | 0.230848 | 0.642 | 0.471 | 1.02E-13 | 5 | Csrnp1 | 0.171 |
| 3.88E-18 | 0.1354989 | 0.287 | 0.173 | 1.21E-13 | 5 | Nfya | 0.114 |
| 4.16E-18 | 0.1547759 | 0.877 | 0.654 | 1.29E-13 | 5 | Celf2 | 0.223 |
| 4.24E-18 | 0.1037481 | 0.11 | 0.047 | 1.32E-13 | 5 | Sgcb | 0.063 |
| 4.55E-18 | 0.1048812 | 0.344 | 0.214 | 1.41E-13 | 5 | Tle4 | 0.13 |
| 4.66E-18 | 0.1131653 | 0.179 | 0.093 | 1.45E-13 | 5 | Soga1 | 0.086 |
| 4.66E-18 | 0.1117964 | 0.561 | 0.387 | 1.45E-13 | 5 | Ankrd11 | 0.174 |
| 4.81E-18 | 0.1281752 | 0.135 | 0.063 | 1.49E-13 | 5 | R3hcc1l | 0.072 |
| 4.84E-18 | 0.1000479 | 0.331 | 0.203 | 1.50E-13 | 5 | Nceh1 | 0.128 |
| 5.73E-18 | 0.119747 | 0.155 | 0.077 | 1.78E-13 | 5 | Ipo9 | 0.078 |
| 5.81E-18 | 0.1160912 | 0.215 | 0.119 | 1.80E-13 | 5 | Rbbp8 | 0.096 |
| 5.86E-18 | 0.1316378 | 0.67 | 0.479 | 1.82E-13 | 5 | Tra2a | 0.191 |

| | | | | | | | |
|---|---|---|---|---|---|---|---|
| 7.96E-18 | 0.1148562 | 0.229 | 0.129 | 2.47E-13 | 5 | Vkorc1 | 0.1 |
| 8.36E-18 | 0.1326127 | 0.272 | 0.163 | 2.60E-13 | 5 | Dguok | 0.109 |
| 8.64E-18 | 0.1017756 | 0.242 | 0.139 | 2.68E-13 | 5 | Epc2 | 0.103 |
| 8.90E-18 | 0.113497 | 0.638 | 0.45 | 2.77E-13 | 5 | Atp6ap2 | 0.188 |
| 9.27E-18 | 0.1190728 | 0.27 | 0.16 | 2.88E-13 | 5 | Nfkb2 | 0.11 |
| 9.35E-18 | 0.1703173 | 0.672 | 0.477 | 2.90E-13 | 5 | Litaf | 0.195 |
| 9.52E-18 | 0.1030446 | 0.271 | 0.16 | 2.96E-13 | 5 | Renbp | 0.111 |
| 9.71E-18 | 0.1013036 | 0.325 | 0.202 | 3.02E-13 | 5 | Ubr2 | 0.123 |
| 1.04E-17 | 0.1011546 | 0.24 | 0.136 | 3.22E-13 | 5 | Cpeb2 | 0.104 |
| 1.06E-17 | 0.1540308 | 0.199 | 0.11 | 3.29E-13 | 5 | Pilrb2 | 0.089 |
| 1.09E-17 | 0.1159998 | 0.293 | 0.179 | 3.38E-13 | 5 | Ebp | 0.114 |
| 1.17E-17 | 0.1024382 | 0.252 | 0.147 | 3.63E-13 | 5 | D5Ertd579e | 0.105 |
| 1.18E-17 | 0.1239483 | 0.151 | 0.075 | 3.66E-13 | 5 | Ppp5c | 0.076 |
| 1.42E-17 | 0.1375505 | 0.143 | 0.069 | 4.40E-13 | 5 | Ptpn23 | 0.074 |
| 1.44E-17 | 0.1495424 | 0.465 | 0.317 | 4.47E-13 | 5 | Hivep2 | 0.148 |
| 1.59E-17 | 0.1521885 | 0.261 | 0.157 | 4.93E-13 | 5 | Trp53 | 0.104 |
| 1.89E-17 | 0.100097 | 0.315 | 0.196 | 5.86E-13 | 5 | 510039O18R | 0.119 |
| 2.13E-17 | 0.2033039 | 0.294 | 0.184 | 6.61E-13 | 5 | Cd180 | 0.11 |
| 2.40E-17 | 0.1073222 | 0.215 | 0.12 | 7.46E-13 | 5 | Stx5a | 0.095 |
| 2.81E-17 | 0.1163838 | 0.476 | 0.321 | 8.74E-13 | 5 | Cysltr1 | 0.155 |
| 3.10E-17 | 0.1123108 | 0.164 | 0.084 | 9.61E-13 | 5 | Dok3 | 0.08 |
| 3.28E-17 | 0.1476497 | 0.219 | 0.124 | 1.02E-12 | 5 | Eif2ak2 | 0.095 |
| 3.28E-17 | 0.1400523 | 0.226 | 0.131 | 1.02E-12 | 5 | Hmgcl | 0.095 |
| 3.62E-17 | 0.1018318 | 0.288 | 0.176 | 1.12E-12 | 5 | Cpsf7 | 0.112 |
| 3.67E-17 | 0.105718 | 0.354 | 0.227 | 1.14E-12 | 5 | Slc4a7 | 0.127 |
| 3.86E-17 | 0.1031105 | 0.239 | 0.138 | 1.20E-12 | 5 | Pisd | 0.101 |
| 4.40E-17 | 0.1112154 | 0.159 | 0.08 | 1.37E-12 | 5 | Zbtb37 | 0.079 |
| 4.44E-17 | 0.1321973 | 0.269 | 0.164 | 1.38E-12 | 5 | Brox | 0.105 |
| 4.73E-17 | 0.1046594 | 0.578 | 0.399 | 1.47E-12 | 5 | Tsc22d4 | 0.179 |
| 4.89E-17 | 0.1244357 | 0.138 | 0.067 | 1.52E-12 | 5 | Ivd | 0.071 |
| 5.50E-17 | 0.1089653 | 0.258 | 0.153 | 1.71E-12 | 5 | Maff | 0.105 |
| 5.98E-17 | 0.1135096 | 0.815 | 0.574 | 1.86E-12 | 5 | Pdia3 | 0.241 |
| 6.01E-17 | 0.1106887 | 0.305 | 0.19 | 1.87E-12 | 5 | Ankhd1 | 0.115 |
| 6.25E-17 | 0.1150136 | 0.263 | 0.158 | 1.94E-12 | 5 | Tlnrd1 | 0.105 |
| 6.70E-17 | 0.1065552 | 0.325 | 0.204 | 2.08E-12 | 5 | Atxn2 | 0.121 |
| 7.23E-17 | 0.1383449 | 0.171 | 0.09 | 2.25E-12 | 5 | Fgd6 | 0.081 |
| 7.74E-17 | 0.1183957 | 0.351 | 0.227 | 2.40E-12 | 5 | Fes | 0.124 |
| 8.12E-17 | 0.1312942 | 0.448 | 0.302 | 2.52E-12 | 5 | Coq10b | 0.146 |
| 8.13E-17 | 0.1222267 | 0.291 | 0.181 | 2.53E-12 | 5 | Rgs19 | 0.11 |
| 8.62E-17 | 0.1369199 | 0.173 | 0.092 | 2.68E-12 | 5 | Plxna4 | 0.081 |
| 9.13E-17 | 0.1219551 | 0.158 | 0.081 | 2.84E-12 | 5 | Ctbs | 0.077 |
| 9.38E-17 | 0.1064291 | 0.419 | 0.278 | 2.91E-12 | 5 | Kpnb1 | 0.141 |
| 1.14E-16 | 0.112701 | 0.188 | 0.102 | 3.53E-12 | 5 | Rcbtb2 | 0.086 |
| 1.14E-16 | 0.1186656 | 0.155 | 0.079 | 3.54E-12 | 5 | Ctc1 | 0.076 |
| 1.20E-16 | 0.1034719 | 0.286 | 0.176 | 3.72E-12 | 5 | Dennd4b | 0.11 |
| 1.22E-16 | 0.1030444 | 0.204 | 0.114 | 3.80E-12 | 5 | Cep120 | 0.09 |
| 1.34E-16 | 0.1179027 | 0.582 | 0.403 | 4.16E-12 | 5 | Evi2a | 0.179 |
| 1.36E-16 | 0.1142607 | 0.22 | 0.126 | 4.24E-12 | 5 | Jmjd6 | 0.094 |
| 1.55E-16 | 0.1081973 | 0.349 | 0.225 | 4.81E-12 | 5 | Dnajb9 | 0.124 |
| 2.10E-16 | 0.1218298 | 0.485 | 0.327 | 6.52E-12 | 5 | Tiparp | 0.158 |
| 2.29E-16 | 0.11788 | 0.116 | 0.053 | 7.10E-12 | 5 | Igtp | 0.063 |
| 2.35E-16 | 0.1138933 | 0.179 | 0.096 | 7.28E-12 | 5 | Aph1b | 0.083 |
| 2.47E-16 | 0.1092485 | 0.117 | 0.054 | 7.68E-12 | 5 | Man2b2 | 0.063 |
| 2.59E-16 | 0.1215581 | 0.315 | 0.198 | 8.03E-12 | 5 | Atp13a2 | 0.117 |
| 2.64E-16 | 0.1153175 | 0.241 | 0.142 | 8.19E-12 | 5 | Kdm6a | 0.099 |
| 2.71E-16 | 0.1251846 | 0.185 | 0.101 | 8.42E-12 | 5 | Aak1 | 0.084 |

| | | | | | | | |
|---|---|---|---|---|---|---|---|
| 2.86E-16 | 0.1077588 | 0.221 | 0.127 | 8.88E-12 | 5 | Filip1l | 0.094 |
| 3.10E-16 | 0.1158549 | 0.276 | 0.17 | 9.61E-12 | 5 | Ә33434E20R | 0.106 |
| 3.18E-16 | 0.109771 | 0.232 | 0.136 | 9.86E-12 | 5 | Hnrnph3 | 0.096 |
| 3.58E-16 | 0.1010231 | 0.219 | 0.126 | 1.11E-11 | 5 | Zufsp | 0.093 |
| 4.01E-16 | 0.1253833 | 0.591 | 0.409 | 1.24E-11 | 5 | Taok3 | 0.182 |
| 4.30E-16 | 0.1040677 | 0.216 | 0.123 | 1.33E-11 | 5 | Trim26 | 0.093 |
| 4.88E-16 | 0.1376723 | 0.918 | 0.723 | 1.51E-11 | 5 | Tmbim6 | 0.195 |
| 5.49E-16 | 0.149588 | 0.915 | 0.721 | 1.70E-11 | 5 | mt-Nd5 | 0.194 |
| 6.88E-16 | 0.1412173 | 0.163 | 0.086 | 2.14E-11 | 5 | Tmem86a | 0.077 |
| 8.12E-16 | 0.1106568 | 0.761 | 0.544 | 2.52E-11 | 5 | Tacc1 | 0.217 |
| 1.04E-15 | 0.1163715 | 0.108 | 0.049 | 3.23E-11 | 5 | Nos1ap | 0.059 |
| 1.08E-15 | 0.109082 | 0.184 | 0.101 | 3.37E-11 | 5 | Zfyve16 | 0.083 |
| 1.16E-15 | 0.1209724 | 0.284 | 0.178 | 3.59E-11 | 5 | Slc9a3r1 | 0.106 |
| 1.20E-15 | 0.1149679 | 0.118 | 0.056 | 3.72E-11 | 5 | Fgd3 | 0.062 |
| 2.01E-15 | 0.1355924 | 0.211 | 0.123 | 6.25E-11 | 5 | Zfp871 | 0.088 |
| 2.04E-15 | 0.1059293 | 0.262 | 0.162 | 6.34E-11 | 5 | Tmem127 | 0.1 |
| 2.07E-15 | 0.1012381 | 0.322 | 0.208 | 6.42E-11 | 5 | Fam91a1 | 0.114 |
| 2.51E-15 | 0.3133419 | 0.209 | 0.124 | 7.79E-11 | 5 | Slc5a3 | 0.085 |
| 2.54E-15 | 0.1550591 | 0.254 | 0.157 | 7.89E-11 | 5 | Ints6 | 0.097 |
| 2.60E-15 | 0.3603243 | 0.401 | 0.28 | 8.06E-11 | 5 | Phlda1 | 0.121 |
| 4.44E-15 | 0.1018205 | 0.357 | 0.237 | 1.38E-10 | 5 | Stt3a | 0.12 |
| 4.83E-15 | 0.1310805 | 0.153 | 0.08 | 1.50E-10 | 5 | Tlr1 | 0.073 |
| 4.89E-15 | 0.1159394 | 0.116 | 0.056 | 1.52E-10 | 5 | Fkrp | 0.06 |
| 5.13E-15 | 0.1175106 | 0.104 | 0.048 | 1.59E-10 | 5 | Donson | 0.056 |
| 5.51E-15 | 0.1118013 | 0.328 | 0.216 | 1.71E-10 | 5 | Szrd1 | 0.112 |
| 6.08E-15 | 0.1247888 | 0.201 | 0.116 | 1.89E-10 | 5 | Dmtf1 | 0.085 |
| 6.41E-15 | 0.1139611 | 0.252 | 0.156 | 1.99E-10 | 5 | Dgkz | 0.096 |
| 1.02E-14 | 0.1305817 | 0.261 | 0.164 | 3.16E-10 | 5 | Abcd1 | 0.097 |
| 1.10E-14 | 0.1054427 | 0.773 | 0.562 | 3.42E-10 | 5 | Rab14 | 0.211 |
| 1.31E-14 | 0.1934396 | 0.117 | 0.057 | 4.08E-10 | 5 | Rcan1 | 0.06 |
| 1.50E-14 | 0.1329123 | 0.219 | 0.13 | 4.65E-10 | 5 | Slc45a4 | 0.089 |
| 1.67E-14 | 0.1001784 | 0.446 | 0.307 | 5.18E-10 | 5 | Ifnar1 | 0.139 |
| 1.76E-14 | 0.1022822 | 0.213 | 0.126 | 5.45E-10 | 5 | Fam193a | 0.087 |
| 2.15E-14 | 0.1658833 | 0.265 | 0.167 | 6.67E-10 | 5 | Kcnq1ot1 | 0.098 |
| 2.78E-14 | 0.1259382 | 0.423 | 0.29 | 8.63E-10 | 5 | Icam1 | 0.133 |
| 2.78E-14 | 0.1089089 | 0.132 | 0.067 | 8.64E-10 | 5 | Siae | 0.065 |
| 3.33E-14 | 0.2351219 | 0.858 | 0.723 | 1.03E-09 | 5 | Pim1 | 0.135 |
| 3.55E-14 | 0.1096428 | 0.161 | 0.088 | 1.10E-09 | 5 | Plcb3 | 0.073 |
| 3.66E-14 | 0.1190463 | 0.929 | 0.75 | 1.14E-09 | 5 | Sdcbp | 0.179 |
| 3.69E-14 | 0.1221185 | 0.318 | 0.212 | 1.15E-09 | 5 | Aup1 | 0.106 |
| 4.40E-14 | 0.103345 | 0.212 | 0.126 | 1.37E-09 | 5 | Mesd | 0.086 |
| 6.70E-14 | 0.1040052 | 0.866 | 0.656 | 2.08E-09 | 5 | App | 0.21 |
| 8.69E-14 | 0.1323024 | 0.185 | 0.107 | 2.70E-09 | 5 | Plk3 | 0.078 |
| 1.01E-13 | 0.1112488 | 0.234 | 0.144 | 3.15E-09 | 5 | Prdm2 | 0.09 |
| 1.09E-13 | 0.1058039 | 0.105 | 0.05 | 3.38E-09 | 5 | Fktn | 0.055 |
| 1.14E-13 | 0.1183715 | 0.186 | 0.108 | 3.55E-09 | 5 | Pknox1 | 0.078 |
| 1.21E-13 | 0.1048907 | 0.311 | 0.204 | 3.77E-09 | 5 | Dhx40 | 0.107 |
| 1.23E-13 | 0.1019986 | 0.201 | 0.119 | 3.83E-09 | 5 | Tmem88 | 0.082 |
| 1.28E-13 | 0.1006309 | 0.183 | 0.106 | 3.98E-09 | 5 | Taf11 | 0.077 |
| 1.33E-13 | 0.1164446 | 0.238 | 0.149 | 4.12E-09 | 5 | Dlgap4 | 0.089 |
| 1.52E-13 | 0.118633 | 0.14 | 0.075 | 4.73E-09 | 5 | Mid1 | 0.065 |
| 1.68E-13 | 0.1016864 | 0.17 | 0.096 | 5.23E-09 | 5 | Pkd1 | 0.074 |
| 1.71E-13 | 0.1034739 | 0.246 | 0.154 | 5.30E-09 | 5 | Slc35f6 | 0.092 |
| 1.87E-13 | 0.1605703 | 0.627 | 0.466 | 5.81E-09 | 5 | Trib1 | 0.161 |
| 1.88E-13 | 0.1264032 | 0.201 | 0.121 | 5.83E-09 | 5 | Btaf1 | 0.08 |
| 2.34E-13 | 0.1315729 | 0.436 | 0.307 | 7.26E-09 | 5 | Lfng | 0.129 |

| | | | | | | | |
|---|---|---|---|---|---|---|---|
| 2.42E-13 | 0.1004701 | 0.105 | 0.051 | 7.51E-09 | 5 | Myc | 0.054 |
| 2.82E-13 | 0.1015348 | 0.186 | 0.11 | 8.75E-09 | 5 | Ilvbl | 0.076 |
| 3.05E-13 | 0.1350595 | 0.197 | 0.119 | 9.46E-09 | 5 | Sec24a | 0.078 |
| 3.05E-13 | 0.1056331 | 0.269 | 0.174 | 9.48E-09 | 5 | Atxn7l1 | 0.095 |
| 3.24E-13 | 0.1039862 | 0.394 | 0.277 | 1.00E-08 | 5 | Samd4b | 0.117 |
| 5.10E-13 | 0.1116428 | 0.176 | 0.102 | 1.58E-08 | 5 | Ddx58 | 0.074 |
| 5.13E-13 | 0.1019016 | 0.14 | 0.075 | 1.59E-08 | 5 | Rab10os | 0.065 |
| 5.40E-13 | 0.1279017 | 0.339 | 0.233 | 1.68E-08 | 5 | Eif4e | 0.106 |
| 6.55E-13 | 0.1572556 | 0.979 | 0.938 | 2.03E-08 | 5 | Junb | 0.041 |
| 7.96E-13 | 0.1732622 | 0.856 | 0.664 | 2.47E-08 | 5 | Cfp | 0.192 |
| 8.77E-13 | 0.6456276 | 0.149 | 0.084 | 2.72E-08 | 5 | Il10 | 0.065 |
| 1.00E-12 | 0.1289468 | 0.235 | 0.149 | 3.12E-08 | 5 | Rabgef1 | 0.086 |
| 1.16E-12 | 0.1101557 | 0.12 | 0.062 | 3.59E-08 | 5 | Zfyve26 | 0.058 |
| 1.21E-12 | 0.1014147 | 0.173 | 0.1 | 3.75E-08 | 5 | Mxi1 | 0.073 |
| 1.24E-12 | 0.1239013 | 0.194 | 0.118 | 3.85E-08 | 5 | Vav2 | 0.076 |
| 1.44E-12 | 0.1265039 | 0.181 | 0.107 | 4.48E-08 | 5 | Swt1 | 0.074 |
| 1.79E-12 | 0.1003669 | 0.155 | 0.088 | 5.57E-08 | 5 | Tmub2 | 0.067 |
| 2.88E-12 | 0.1136377 | 0.147 | 0.082 | 8.96E-08 | 5 | Zbtb21 | 0.065 |
| 3.16E-12 | 0.1241964 | 0.268 | 0.175 | 9.80E-08 | 5 | Rasgef1b | 0.093 |
| 3.92E-12 | 0.2822739 | 0.261 | 0.173 | 1.22E-07 | 5 | Dusp2 | 0.088 |
| 9.62E-12 | 0.1042241 | 0.142 | 0.08 | 2.99E-07 | 5 | Ctps2 | 0.062 |
| 1.04E-11 | 0.1107192 | 0.139 | 0.077 | 3.24E-07 | 5 | Sgsh | 0.062 |
| 1.22E-11 | 0.1157993 | 0.146 | 0.083 | 3.78E-07 | 5 | Tesk1 | 0.063 |
| 2.10E-11 | 0.1073289 | 0.252 | 0.167 | 6.52E-07 | 5 | Sde2 | 0.085 |
| 2.90E-11 | 0.1400449 | 0.793 | 0.609 | 8.99E-07 | 5 | Wfdc17 | 0.184 |
| 3.39E-11 | 0.1014552 | 0.114 | 0.061 | 1.05E-06 | 5 | Caml | 0.053 |
| 3.43E-11 | 0.1364263 | 0.973 | 0.763 | 1.06E-06 | 5 | Tyrobp | 0.21 |
| 4.00E-11 | 0.1031115 | 0.777 | 0.587 | 1.24E-06 | 5 | Pnrc1 | 0.19 |
| 4.69E-11 | 0.1035669 | 0.156 | 0.092 | 1.46E-06 | 5 | Cep250 | 0.064 |
| 1.74E-10 | 0.1521776 | 0.949 | 0.832 | 5.41E-06 | 5 | Neat1 | 0.117 |
| 5.27E-10 | 0.1004248 | 0.163 | 0.101 | 1.64E-05 | 5 | Mfsd10 | 0.062 |
| 1.04E-09 | 0.1717511 | 0.554 | 0.425 | 3.23E-05 | 5 | Sbno2 | 0.129 |
| 2.27E-09 | 0.1091942 | 0.747 | 0.57 | 7.05E-05 | 5 | Plek | 0.177 |
| 1.99E-08 | 0.4070088 | 0.632 | 0.556 | 0.0006177 | 5 | Il1b | 0.076 |
| 3.83E-08 | 0.1057828 | 0.249 | 0.175 | 0.0011897 | 5 | Chka | 0.074 |
| 4.73E-08 | 0.1069589 | 0.913 | 0.783 | 0.0014695 | 5 | Cebpb | 0.13 |
| 9.31E-08 | 0.1168062 | 0.226 | 0.158 | 0.0028917 | 5 | Arl5c | 0.068 |
| 1.32E-07 | 0.361921 | 0.424 | 0.349 | 0.0040894 | 5 | Id2 | 0.075 |
| 2.96E-07 | 0.114867 | 0.203 | 0.142 | 0.0091871 | 5 | Rnd3 | 0.061 |
| 7.00E-07 | 0.1437924 | 0.899 | 0.768 | 0.0217506 | 5 | Zfp36l2 | 0.131 |
| 5.26E-06 | 0.1036624 | 0.201 | 0.147 | 0.1634704 | 5 | Gadd45g | 0.054 |
| 2.56E-05 | 0.1029086 | 0.956 | 0.865 | 0.7943038 | 5 | Klf6 | 0.091 |
| 2.82E-05 | 0.3448634 | 0.113 | 0.078 | 0.8744727 | 5 | Cxcl10 | 0.035 |
