## Supplementary material for "Nerve-associated macrophages control adipose homeostasis across lifespan and restrain age-related inflammation": Table S3

Tnfsf9
Ccl12
Ccl7
St3gal6
Col14a1
Marcksl1
Vcam1
Blnk
Bmp2
Ccr2
Egr1
Fnbp1l
Cbr2
Cyr61
F13a1
Fcrls
Gas7
Igfbp4
Lifr
Mgl2
Pmp22
Hivep2
Rbpj
Ccnd1
Il10
Itga6
Cd163
Cfh
Il21r
Rgs18
Apold1
S100a4
Akt3
Gpr165
Serpinf1
Ccl2
Ccl4
Ccl9
Cp
Il1rl1
Kitl
Clec10a
Lat2

Pf4
Tagap
Ch25h
Mef2c
Plxna4
Folr2
Gas6
Lyve1
Sdc4
Clec12a
Wwp1
Arap3
Ptgs2
Bend4
Gm8995
Nfkbiz
Socs3
Skil
Sesn1
1700025G04Rik
Arhgap22
Cdk6
Cdr2
Dennd2c
Ptpro
Susd3
Atp8b4
Rtp4
Cxcl1
Cxcl2
Dlc1
Etv1
Ier3
Tln2
Tnfaip3
Zfp37
Abca9
Cd209a
Retnla
Mmp13
Olfml3
Man1a
Stard8

1810011O10Rik
Cd40
Hpgd
Itgam
Tlr1
