## Supplementary material for "Nerve-associated macrophages control adipose homeostasis across lifespan and restrain age-related inflammation": Table S5

| Gene Name | Forward Primer |
|---|---|
| *Actb* | CTC TGG CTC CTA GCA CCA TGA AGA |
| *Hprt1* | ACA GGC CAG ACT TTG TTG GA |
| *Adrb3* | GGA AGC TTG CTT GAT CCC CAT |
| *Pparg* | TCC AGC ATT TCT GCT CCA CA |
| *Gdf3* | TCT CCC AGA CCA GGG TTT TT |
| *Nlrp3* | GCT AAG AAG GAC CAG CCA GA |
| *Asc* | TAC AGC CAG AAC AGG ACA CTT T |
| *Casp1* | GGA CCC TCA AGT TTT GCC CT |
| *Il1b* | GGT CAA AGG TTTGGA AGC |
| *Il18* | GAC AGC CTG TGT TCG AGG AT |
| *Il10* | GGG TTG CCA AGC CTT ATC GGA AAT |
| *Il6* | AGA CAA AGC CAG AGT CCT TCA GAG |
| *Tnf* | TCT CAG CCT CTT CTC ATT |

| Reverse Primer |
|---|
| GTA AAA CGC AGC TCA GTA ACA GTC CG |
| ACT TGC GCT CAT CTT AGG CT |
| AGT TGA GCT TCC AGG GGA CT |
| AAG GTG GAG ATG CAG GTT CT |
| AAT AGA GGA CCT TCT GGA GAC A |
| CAG CAA ACC CAT CCA CTC TT |
| AAA GCA TCC AGC ACT CCG TC |
| AGA CGT GTA CGA GTG GTT GT |
| TGT GAA ATG CCA CCT TTT |
| CAG TCT GGT CTG GGG TTC AC |
| TCT TCA GCT TCT CAC CCA GGG AAT |
| TTG GTC CTT AGC CAC TCC TTC TGT |
| AGA ACT GAT GAG AGG GAG |
